# Clinically observed *RASA1* missense mutants exhibit diverse RasGAP protein behaviors

**DOI:** 10.1101/2025.10.03.680326

**Authors:** Maxum E. Paul, Rediet B. Delelegne, Jocelyn E. Chau, Titus J. Boggon

## Abstract

The *RASA1* gene is mutated in cerebrovascular disorders and cancer, yet how the resulting mutations in the GTPase Activating Protein, RasGAP (p120RasGAP, RASA1) dysregulate signaling remains poorly understood. Here, we catalogue currently reported disease-associated mutations in *RASA1* and assess their impact on RasGAP protein *in vitro*. On mapping these mutations onto experimental structures and structural models of RasGAP we identify regions that suggest functional impact. We assess key mutations within these regions for their effects on protein expression, thermal stability, and their interactions with a known binding partner, p190RasGAP. We then assess Michaelis-Menten kinetics of the mutant RasGAP proteins towards Ras. Together, we find that disease-associated RasGAP mutations classify into a panel of distinct classes based on their mode of dysregulation. We demonstrate that protein stability is necessary but not sufficient for full catalytic activity and that destabilizing mutations across the length of the protein can disrupt this function, but that the C2 domain appears to be unique in its role of regulating GAP activity by mechanisms other than destabilization involving the interactions of specific residues.

## Introduction

Small GTPases cycle between GTP-bound and GDP-bound states and generally signal to downstream pathways only when bound to GTP (1–3). For the Ras superfamily, proper regulation of this GTP-dependent signaling is essential for cellular processes that include growth, proliferation, and differentiation (4, 5). This regulation is in large part achieved by the actions of guanine nucleotide exchange factors (GEF) which initiate signaling by displacing GDP to allow GTP binding to the GTPase, and GTPase Activating Proteins (GAP) which effectively turn off downstream signaling by accelerating GTPase hydrolysis of GTP (1, 6, 7) (**Figure 1A**). Thus, the balance of GEF and GAP activity is essential to control the amount and duration of GTP-bound GTPase, and appropriate downstream signaling. Importantly, cellular processes are affected when this balance is disrupted, so alterations in GAP activity have the potential to drive disease (3, 4).

**Figure 1.**
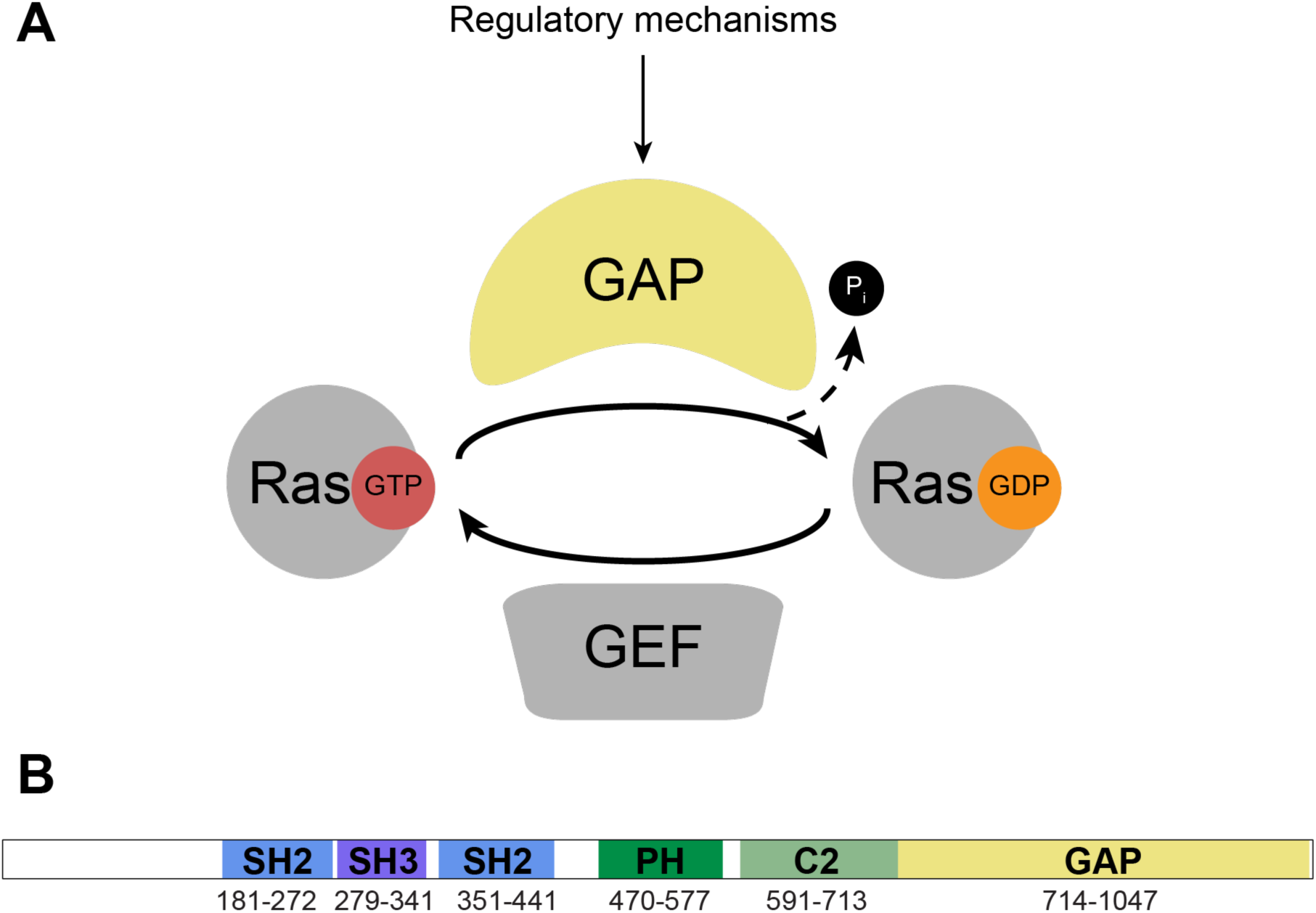
The small GTPase cycle, and the RasGAP protein. **A)** The GTPase cycle illustrating the conversion of GTP-bound Ras to GDP-bound Ras by GAP stimulation, and return to its GTP-bound form via GEF. Selected inputs that alter GAP activity are indicated. **B)** Domain architecture of RasGAP. Domain folds are indicated as: SH2, Src homology-2; SH3, Src homology-3; PH, pleckstrin homology; C2, protein kinase C domain 2; GAP, GTPase activating protein. Residue numbers indicating approximate domain boundaries are shown.

RasGAP (p120RasGAP, RASA1; gene: *RASA1*) was the first GAP to be identified (8), crystallized (9), and co-crystallized with a small GTPase (10, 11). These studies revealed that it, and other GAP proteins, accelerate GTPase hydrolysis of GTP by optimizing the chemistry of hydrolysis. They achieve this by insertion of an ‘arginine finger’ from the GAP into the catalytic cleft of the GTP bound small GTPase, the importance of which is illustrated by mutations in this arginine (R789 in RasGAP) resulting in complete disruption of GAP-mediated acceleration of hydrolysis (10, 12). In cells, such mutations result in increased GTP-bound small GTPase and consequent downstream signaling (13, 14). Similarly, other mutations in RasGAP can impact GAP activity and are associated with increased Ras signaling (15). Unsurprisingly, therefore, Mendelian or acquired mutations in RasGAP have been found to be associated with disease states, including the vascular disorders Capillary Malformations-Arteriovenous Malformations (CM-AVM) (16–18) and Vein of Galen Malformations (VOGM) (18, 19), and a range of cancers (20–29) (**Supplementary Tables 1 and 2**).

RasGAP is a multi-domain protein, comprising a disordered N-terminal region, a Src Homology-2 domain, (N-SH2), a Src Homology 3 domain (SH3), a second SH2 domain (C-SH2), a pleckstrin homology domain (PH), a C2 domain that does not bind calcium, and a C-terminal catalytic GAP domain (**Figure 1B**). These domains perform a range of functions, including recruitment to phosphotyrosine-containing partner proteins by the SH2 domains (30–34), membrane localization by the PH domain (35–37), and cross talk with and regulation of RhoGAP proteins by both the SH2 and SH3 domains (38–42). Disease-associated mutations may therefore result in functional consequences that impact RasGAP signaling by a variety of mechanisms, including potential alterations in enzymatic activity, localization, binding to partner proteins, interactions with small GTPase, or alterations of protein stability (**Figure 1A**). As has been shown in other signaling pathways (43, 44), understanding these differences may be important to better understand the mechanisms that drive individual cases, or groups of, disease.

Recently, we revealed new insights into the function of wild type RasGAP and the mechanism by which a disease mutation disrupts this function. Analyses of a VOGM-associated mutation in the C2 domain of RasGAP, arginine 707, were key to demonstrate that this domain augments GAP activity towards Ras. This finding seems applicable across the entire family of GAP proteins that target Ras and thus provides new understanding of signaling in this class of GAP proteins (45). As the roles of the RasGAP domains are not fully understood, it is likely that analysis of disease mutations can inform on the function of the wild type protein as well as the mechanism by which the disease mutation may disrupt this function.

In this study, we assess disease-associated mutations in RasGAP. We compile an extensive catalogue of published missense mutants found in both vascular malformations and in cancer and analyze these mutations based on crystal structures and AlphaFold models of RasGAP. In these analyses we find a range of different classes of mutations, and we functionally analyze twelve that we consider to be representative of the different classes. Enzymatic assays on purified RasGAP are used to obtain Michaelis-Menten kinetics profiles for both wild type and mutant proteins. These analyses reveal that the mutations differentially impact kinetic activity towards Ras. Similarly, protein stability assessment reveals differential impacts. Finally, we find that some SH2 mutants do not alter affinity for known binding partners. Together, we observe that disease-associated RasGAP mutations classify into different groups, revealing the variety of *in vitro* behaviors that can be associated with disease phenotypes *in vivo*.

## Results

### Structure-based assessment of clinically observed RasGAP mutations

Clinically-observed mutations in *RASA1* are relatively well documented, particularly in the vascular disorders CM-AVM (16–18) and VOGM (18, 19), as well as in cancer (20–29). We therefore began by conducting a search for mutations in the *RASA1* gene using the literature and publicly available databases. We focused on codon changing (missense) mutations because these can be particularly informative based on their specific location of the protein. In contrast, frame shifts and truncations generally result in protein degradation. In the case of RasGAP truncations the loss of the C-terminal GAP domain will necessarily result in a catalytically inactive protein, we therefore did not assess frame shifts or truncations. We compiled lists of clinically observed *RASA1* mutations found in vascular malformations (**Supplementary Table 1**), and in cancers (**Supplementary Table 2**), and manually removed database duplications. We noted that *RASA1* mutations documented in vascular malformations are predominantly found in individual studies in the literature, but that *RASA1* mutations in cancers are more frequently found in database compilations.

Analysis of the compiled tables reveals that both vascular malformation and cancer mutations occur across the length of the RasGAP protein (**Figure 2**). We find that some mutations are frequently observed; for example, mutations at R427, R591 and R749 are each found over ten times. We view this as significant despite the observation that in cancer genome sequencing databases arginine is the most commonly mutated residue (46, 47). When assessing both lists, we find that mutations in a total of nine codons are observed in both vascular disorders and in cancer (**Supplementary Table 3**). Because disorders that harbor *RASA1* mutations vary in phenotype, we reasoned that differences in functional effects might be expected to occur for mutations in different regions of RasGAP.

**Figure 2.**
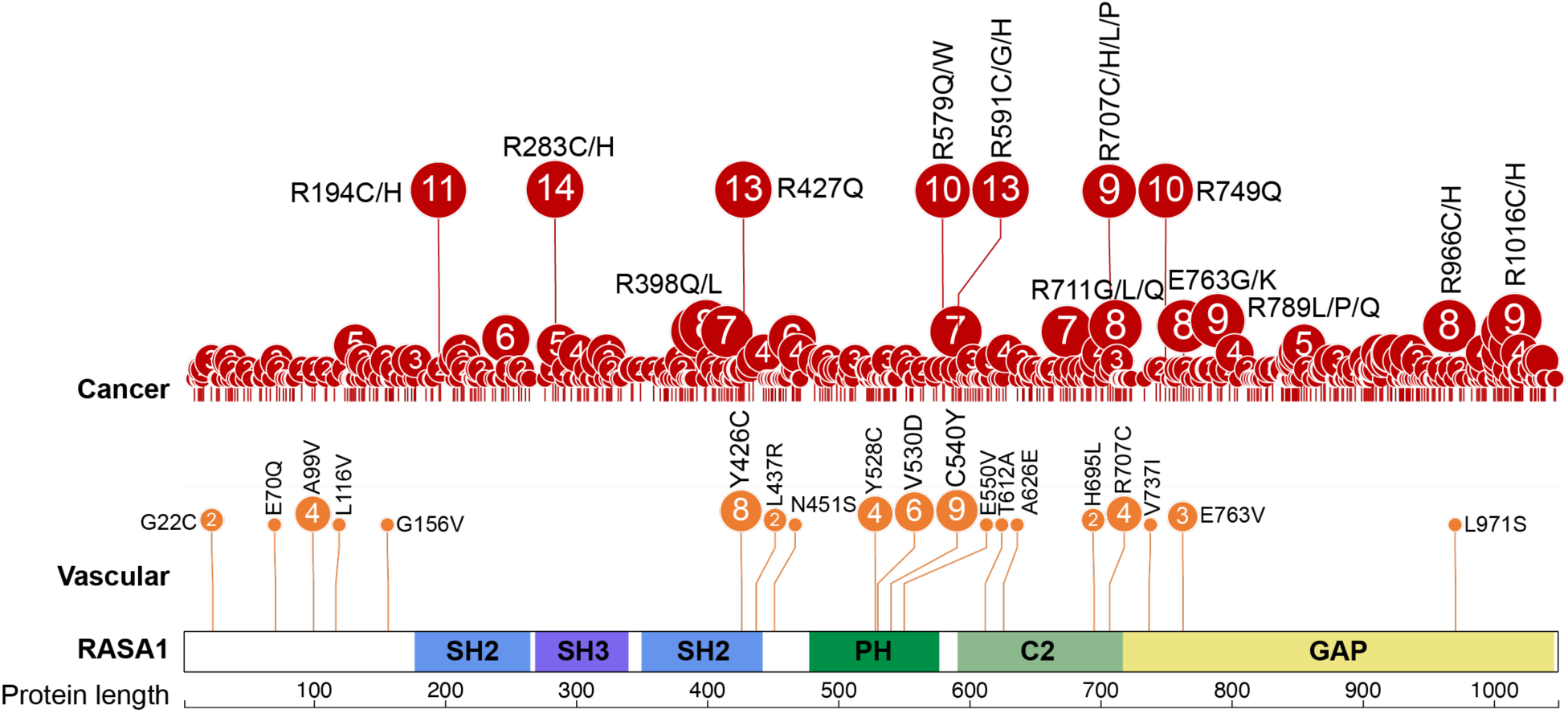
Map of *RASA1* missense mutations. *RASA1* missense mutations reported in the literature are indicated on the domain map of RasGAP. Circle size indicates frequency of a given mutation from either cancer (top, red) or vascular disorders (bottom, orange).

We assessed the AlphaFold model for RasGAP (AlphaFold DB accession code: AF-P20936-F1-v4) in combination with the two multi-domain crystal structures of RasGAP, of the SH2-SH3-SH2 region (Protein Data Bank accession code: 8DGQ (33)) and the C2-GAP region (Protein Data Bank accession code: 9BZ4 (45)) (**Figure 3ABC, Supplementary Figure 1ABC**). On each of these structures we mapped the surface electrostatics to allow us to visualize regions which are positively charged (blue), negatively charged (red) and hydrophobic (white) (**Figure 3DEF, Supplementary Figure 1DEF**). We next generated a sequence alignment of RasGAP across 209 species, from sponges to humans and used this sequence alignment to assess conservation on the surface of RasGAP by coloring highly conserved residues blue and poorly conserved residues white (**Figure 3GHI, Supplementary Figure 1GHI, Supplementary Figure 2**). Together these structural assessments provide a clear understanding of predicted domain-domain interactions and potential partner binding sites.

We next mapped our panel of mutations onto the three structures, with a color scheme that indicates frequency of observation (**Figure 3JKL, Supplementary Figure 1JKL**). Mutations observed at a low frequency are observed across the surface of the predicted and experimental RasGAP structures, perhaps indicating some level of database noise or deleterious functional effects across the protein. In contrast, mutations that are observed frequently (we chose a cut-off of four observations) cluster to regions of RasGAP with high conservation throughout evolution (**Supplementary Figure 3**). A patch of disease mutations is observed on the surface of the GAP domain in a region known to interact directly with Ras (**Supplementary Figure 3**)(10). We reasoned that mutations in the GAP domain likely interrupt Ras binding and would naturally lead to changes in activity, similar to mutations in the arginine finger, R789, so we did not further study these sites.

Instead, we focused on mutations in the accessory domains of RasGAP. We selected twelve codons that are frequently mutated in cancer or mutated in vascular malformations, two of which are mutated in both cancer and vascular malformations, Y528 and R707. Of the twelve mutated codons, one is found in each of the N-SH2, SH3, and C-SH2 domains (R194, R283, and R427, respectively), one is in the PH domain (Y528), and eight are in the C2 domain (R591, H604, T612, A626, W689, H695, S705 and R707) (**Figure 3MNO, Supplementary Figure 1MNO**). All of these residues, except H695, are very highly conserved over evolution (**Supplementary Figure 3)**. Each of these mutations were selected for their predicted role and /or frequency in disease.

We selected the SH2 and SH3 domain mutants R194C, R283H and R427Q which were highly conserved and frequently observed in cancer, possibly suggesting an important role. Two of these residues (R194 and R283) are distant from known sites of binding or protein-protein interactions, raising the possibility that they may have previously unappreciated functions. Conversely, R427 is located adjacent to the phosphotyrosine site of C-SH2, potentially suggesting an impact on binding (**Figure 3M-N, Supplementary Figure 4A-C**). Similarly, we selected the PH domain mutant, Y528C, because of its potential to impact inter-domain interactions (**Figure 3M**, **Supplementary Figure 4D**). The C2 domain mutants were selected to address potential roles of this domain (**Figure 3M,O, Supplementary Figure 4E-I**). R591C was chosen because it forms a salt bridge to the GAP domain, potentially impacting the relative orientations of the C2 and GAP domains. T612A was chosen because it is in a location that is sometimes important for calcium binding in C2 domains, A626E because it was predicted to interrupt the hydrophobic core of the C2 domain, and H695L because it is a loop residue at the surface of the C2 domain contiguous with a conserved basic patch in the PH domain and distal from T612A. In previous studies, the C2 domain was found to augment Ras activity by interacting with Ras, so we selected mutants predicted to disrupt this augmentation, including R707H (distinct from the previously studied R707C), H604N, W689R, S705F. Together these mutations provide a range of possible effects on RasGAP function (**Figure 3M-O, Supplementary Figure 4**).

**Figure 3.**
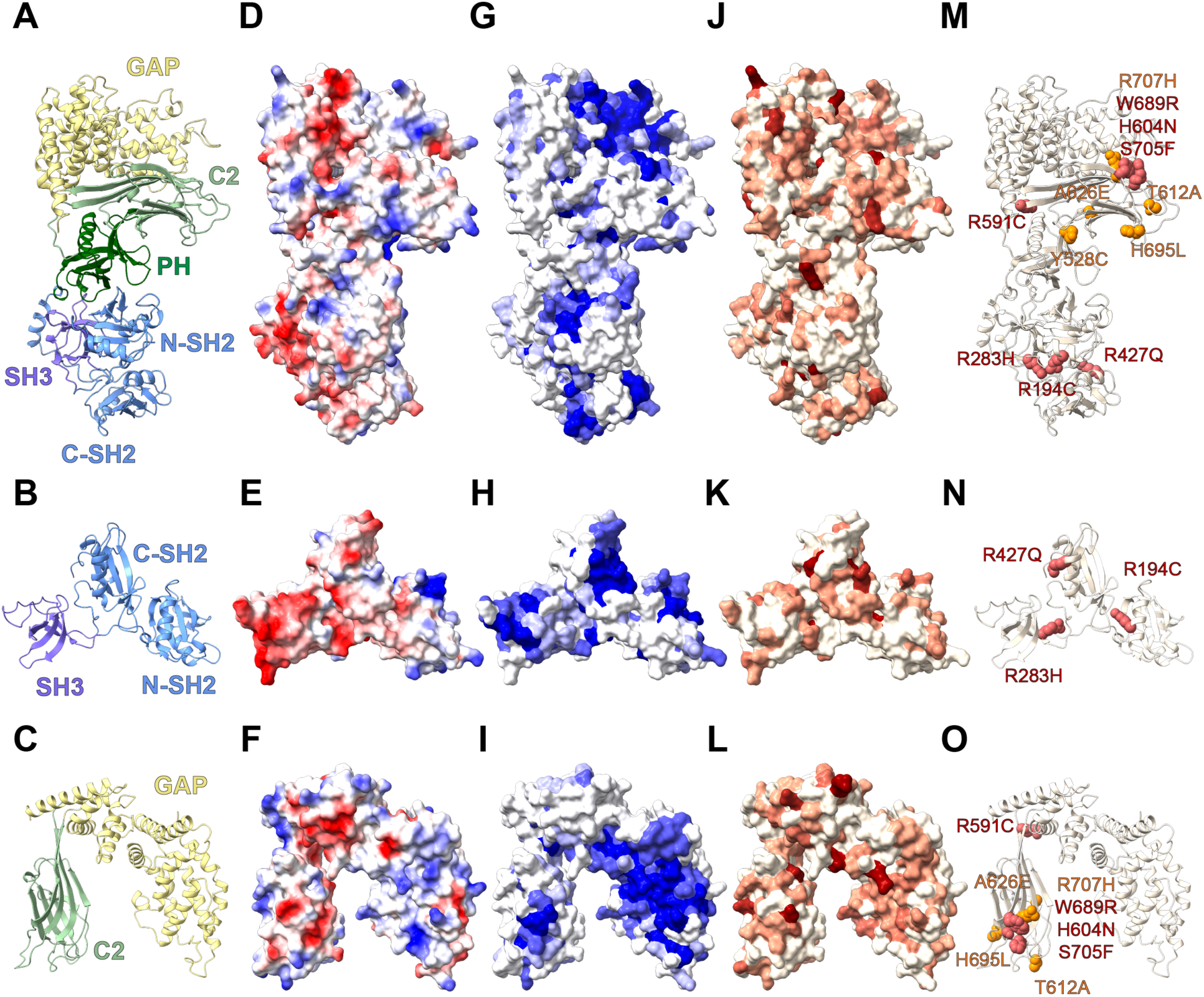
Mapping *RASA1* mutations onto RasGAP. The figure depicts the AlphaFold prediction of ordered domains of RasGAP (**A**, **D**, **G**, **J**, **M**, top row) (AlphaFold ID: AF-P20936-F1-v4), crystal structure of the SH2-SH3-SH2 region (**B**, **E**, **H**, **K**, **N**, middle row) (PDB accession number: 8DGQ (20)), and crystal structure of the C2-GAP region (**C**, **F**, **I**, **L**, **O**, bottom row) (PDB accession number: 9BZ4 (30)). **A-C)** Ribbon diagrams colored by domain. **D-F)** Electrostatics of the protein surface. Positive (blue) and negative (red) charge indicated. **G-I)** Sequence conservation from low (white) to high (blue) mapped onto the protein surface. **J-L)** *RASA1* mutations mapped onto the protein surface. No reported mutations colored white, most frequent colored dark red. **M-O)** Mutations chosen for further study shown as spheres. Red indicates cancer-associated, orange indicates vascular malformations associated.

### Impact of mutations on protein stability

We decided to assess the stability of the panel of RasGAP mutants identified by our structural analysis. Loss of protein stability is a potential mechanism for reduced GAP activity and may be consistent with observations of the studied mutations in human disease. We therefore expressed and purified each of these mutants in constructs coding for the six ordered domains of p120RasGAP using our *E. coli* expression system and FPLC purification protocol (34, 45). We reasoned that significant loss of stability in an *in vitro* setting might indicate rationale for loss of function in cells, and consequently in disease. We find that of the twelve mutant proteins, eleven are soluble from *E. coli* and can be purified using standard protocols. In contrast, mutation A626E, located in the hydrophobic core of the C2 domain (**Supplementary Figure 4G**) results in an insoluble protein (**Supplementary Figure 5**). This raises the possibility that protein destabilization is the mechanism by which this A626E mutation impacts RasGAP activity in disease.

To assess protein stability in more detail we evaluated the melting temperatures of each of the remaining eleven mutants. We did this using thermal shift assays, a quantitative method to assess protein denaturation using differential scanning fluorimetry and determined the melting temperature of each mutant compared to wild type (**Figure 4**, **Supplementary Table 4**). We find that melting temperature does not significantly change for a majority of mutants (R427Q, R591C, H604N, T612A, W689R, H695L, and S705F), suggesting that degraded protein stability may not drive disease impacts for these variants. In contrast, we find that R194C mutation in the N-SH2 domain, R283H in the SH3 domain, and Y528C in the PH domain, as well as R707H in the C2 domain, all deleteriously impact protein stability. We hypothesize that these observed instabilities could, at least in part, impact catalytic activity.

**Figure 4:**
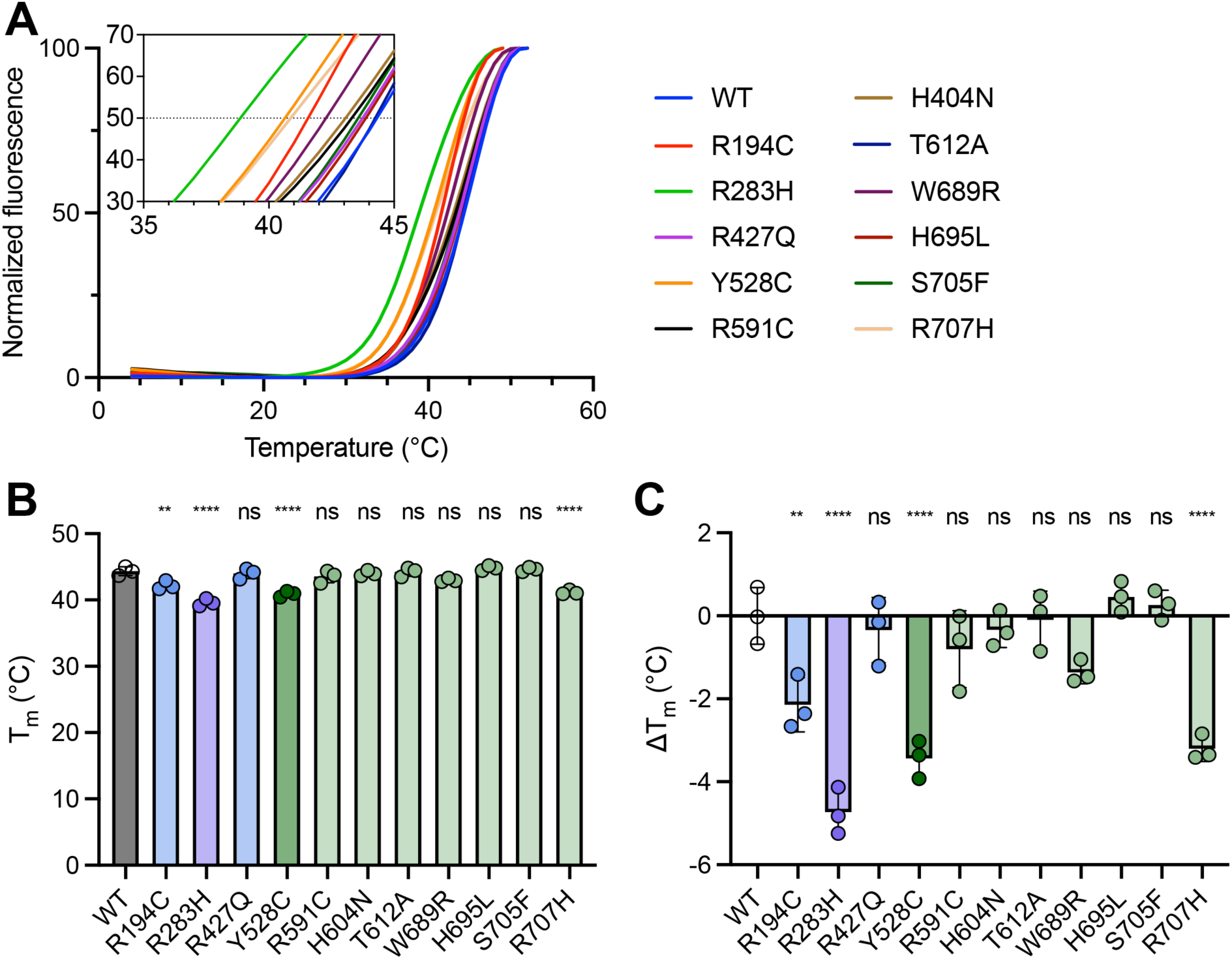
Thermal stability of RasGAP disease mutants. **A)** Melting curves for each RasGAP construct. One representative replicate shown for each. Inset shows closeup of curves around the inflection point. **B)** Melting temperature (Tm) of RasGAP constructs. Bars indicate mean ± SD (n = 3 technical replicates). **C)** Comparison to wild type Tm (44.35°C) for each construct (ΔTm). Comparisons to wild type using ordinary one-way ANOVA with Dunnett correction for multiple comparisons shown: ** indicates *P* < 0.01, **** indicates *P* < 0.0001, ns indicates not significant. *F* = 25.82, overall *P* < 0.0001, *R*^2^ = 0.9221.

### Protein-protein interactions

We next asked whether the R194C and R427Q mutations would affect the ability of the SH2 domains to bind tyrosine-phosphorylated peptides. We therefore assessed the interactions of RasGAP with one of its known phosphorylated binding partners, p190RhoGAP (ARHGAP35) (33, 38, 48). We conducted isothermal titration calorimetry (ITC) to assess the binding of a synthesized phosphopeptide corresponding to p190RhoGAP residues 1083-1111 (phosphorylated on pY1087 and pY1105) with a RasGAP construct encoding the N-SH2, SH3 and C-SH2 domains (**Supplementary Figure 6A-G** and **Supplementary Table 5**). In our ITC experiments we observe no significant difference in affinity for the mutants compared to wild type, suggesting no impact on the ability of the SH2 domains to engage phosphotyrosine binding partners.

### Michaelis-Menten kinetics of RasGAP mutants

We then turned to kinetic analysis of RasGAP activity towards Ras hydrolysis of GTP. We reasoned this might facilitate a better understanding of the mechanisms by which *RASA1* mutants might be associated with disease. We conducted *in vitro* GAP assays using the fluorescent Phosphate Sensor system (49, 50) to monitor real-time phosphate release (45, 51–53). H-Ras pre-loaded with GTP (54) was used as the substrate and Michaelis-Menten kinetics for wild type RasGAP and the eleven soluble mutants assessed allowing comparison of kcat, KM and catalytic efficiency (kcat/KM) for each RasGAP mutant.

Our analyses reveal that wild type RasGAP exhibits kcat on the order of 20 s^-1^, KM on the order of 50 µM, and catalytic efficiency on the order of 400,000 M^-1^s^-1^, parameters similar to those observed in previous studies (45) (**Figure 5, Supplementary Table 6**). This baseline allows comparison of mutants with wild type protein. We find that the R194C mutation in N-SH2 and R283H in the SH3 domain both show reduced catalytic efficiency, primarily driven by weaker KM of GAP interaction with GTP-bound Ras (**Figure 5**). We interpret this to potentially result from the reduced stability of these mutant proteins which we observed in thermal stability assays (**Figure 4**). In contrast, R427Q in the C-SH2 domain is not significantly different from wild type protein in catalytic efficiency, suggesting that effects of this mutation may not directly alter the catalytic activity of RasGAP (**Figure 5**). The PH domain mutant Y528C shows moderately impaired catalytic efficiency, again potentially resulting from the observed destabilization for this mutation (**Figure 5**).

**Figure 5.**
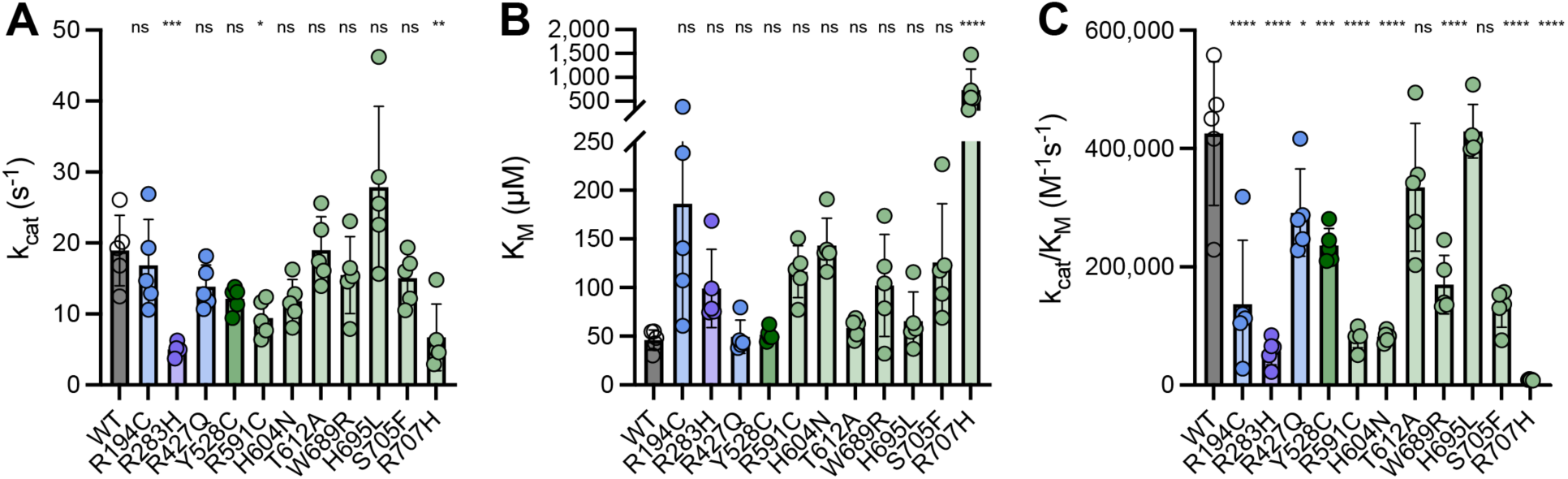
Michaelis-Menten kinetics for disease-associated RasGAP mutants. Michaelis-Menten parameters from single-turnover phosphate release assays. **A)** kcat, **B)** KM, and **C)** catalytic efficiency (kcat/KM) shown for wild-type and each mutant. Bars indicate mean ± SD (n = 5 technical replicates). Bars are colored according to the domain in which the mutation is found. SH2, blue; SH3, purple; PH, dark green; C2, light green. Statistical significance determined via ordinary one-way ANOVA with Dunnett’s multiple comparisons test. *P* values for comparisons of each mutant against WT are represented: ns, not significant; ****, *P* < 0.0001; ***, *P* < 0.001; **, *P* < 0.01; *, *P* < 0.05; ns indicates not significant. ANOVA for kcat values in **A**: *F* = 7.249, overall *P* < 0.0001, *R*^2^ = 0.6242. ANOVA for KM values in **B**: *F* = 9.606, overall *P* < 0.0001, *R*^2^ = 0.6876. ANOVA for kcat/KM values in **C**: *F* = 24.07, overall *P* < 0.0001, *R*^2^ = 0.8465.

For the mutants found in the C2 domain we observe a range of effects. Two, T612A and H695L, located on the periphery of the C2 domain (**Figure 3**) show little effect on activity compared to wild type (**Figure 5**). This correlates with their stability (**Figure 4**) and the finding that the T612A equivalent is not embryonic lethal in mice (45), implying that the disease-associated impacts of these mutations may be orthogonal to either protein stability or catalytic activity. In contrast, R591C also shows negligible change in stability (**Figure 4**) but a significant reduction in catalytic efficiency (**Figure 5**). Our analysis of the crystal and AlphaFold structures suggests that this residue forms a salt bridge between the C2 and GAP domains by interaction with D990 of the GAP domain (**Supplementary Figure 4E**). This is one of a limited number of direct contacts between the GAP and C2 domains; thus we interpret that disrupting this salt bridge may alter kinetics, potentially by unfavorably affecting the relative position or orientation of the C2 domain relative to the GAP domain.

The remaining mutations reside on the surface of the C2 domain predicted to interact with the allosteric lobe of Ras (**Figure 3**). Three of these (H604N, W689R, and S705F) show little to no impact on protein stability and little impact on kcat. In contrast, each of these has a higher (weaker) KM for GTP-bound H-Ras and consequently significantly reduced catalytic efficiency (**Figure 5**). We interpret these results to show that the extended predicted C2/Ras surface is important for RasGAP signaling. Finally, we find that mutation R707H shows both a very significant reduction in kcat and an increased KM, consistent with previous observations for mutations at this site (**Figure 5**). Protein stability for this mutant is not as degraded as for other mutations (**Figure 4**), but activity is significantly impaired; thus we hypothesize that both protein stability and enzymatic activity may be impacted by this mutant.

Finally, we assessed catalytic efficiency in the context of melting temperature. We plotted these values against one another to assess whether protein stability systematically impacts catalytic efficiency (**Figure 6**). We observe that although lower melting temperature does correlate with lower activity, outsized effects of individual mutations can significantly impact catalytic activity. We also examined these results in search of a trend in catalytic efficiency and /or stability compared to the disease origin of each mutation (cancer or vascular malformations). No trend was observed in either case, with wide variations in both kcat/KM and Tm for mutations from both sets of disease literature. This indicates that reduction of GAP activity and /or stability may be correlated with both groups of diseases, as may other potential alterations in RasGAP function.

**Figure 6.**
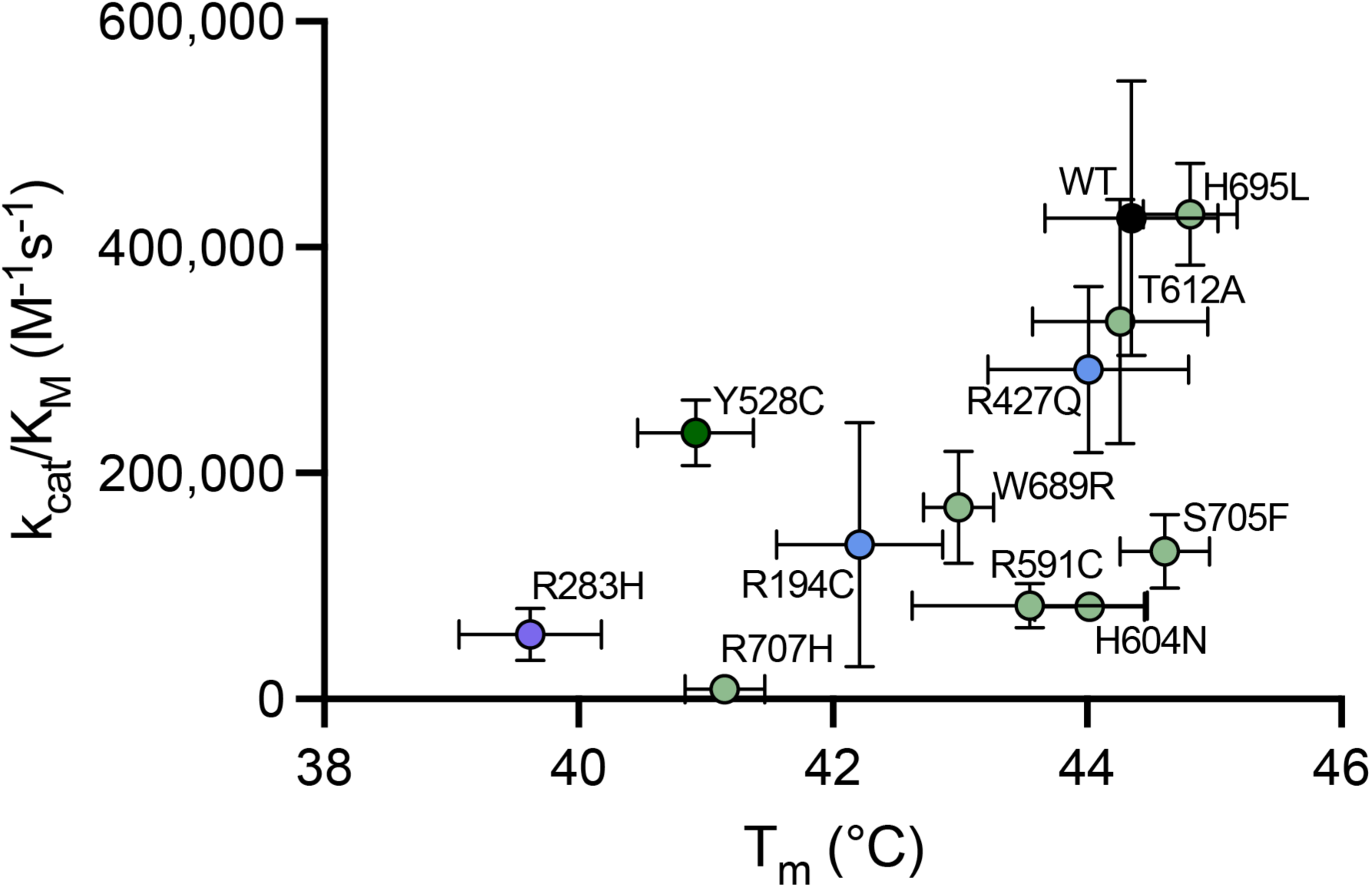
Comparison of catalytic efficiency and protein stability for RasGAP mutants. Plot of catalytic efficiency (kcat/KM, Figure 5C) against melting temperature (Tm, Figure 4B) for each RasGAP construct. Error bars indicate mean ± SD (n = 5 for GAP assays, n = 3 for thermal shift assays). Points are colored according to the domain in which the mutation is found. SH2, blue; SH3, purple; PH, dark green; C2, light green.

## Discussion

In this study we conducted biochemical and biophysical analyses to assess if disease-associated mutations in *RASA1* might result in a range of functional effects for the purified RasGAP protein. Our study catalogued the known codon-changing mutations in the resulting RasGAP protein and used a structure-based approach to rationalize these mutations. This assessment yielded clear regions on the surface of RasGAP protein that are highly mutated in disease, a finding that led us to ask whether these clusters have varying impacts on functional outcomes. We find this to be the case and believe that RasGAP mutations can be categorized into groups that i) result in complete loss of protein folding and solubility in an ectopic prokaryotic system, ii) destabilize the expressed protein, iii) reduce catalytic rate (kcat) and /or weaken GTPase binding (KM), or iv) alter RasGAP function by another mechanism (**Figure 7**). These categorizations can help rationalize the clinical variety resulting from *RASA1* mutations.

**Figure 7.**
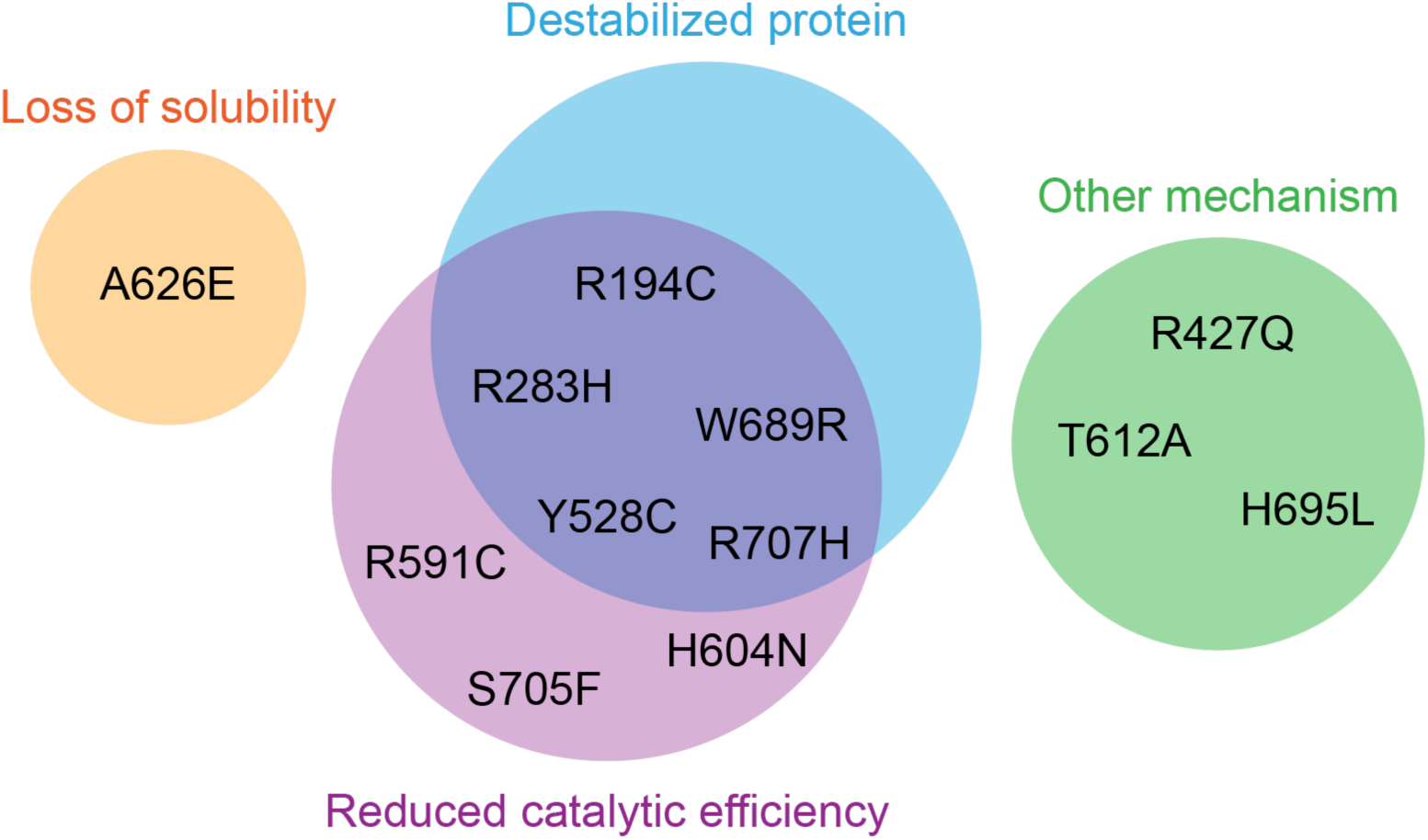
Mechanisms of altered RasGAP signaling. RasGAP mutations categorized into groups that i) result in complete loss of solubility in an ectopic expression system, ii) destabilize the expressed protein, iii) reduce catalytic rate (kcat) and /or weaken GTPase binding (KM), or iv) alter RasGAP function by another mechanism.

As with many genes in RAS signaling pathways, *RASA1* mutations are well represented in the cancer databases. This contrasts with the categorization of *RASA1* mutations associated with vascular disorders, which are in general published as clinical observations or cases and not reported to centralized databases. Our analysis takes this into account (**Supplementary Tables 1 and 2**), and we observe overlap between the cancer and vascular mutations (**Supplementary Table 3**). Mapping this extensive catalogue of mutations onto the multi-domain crystal structures and AlphaFold model resulted in observation of several regions surface regions that are recurrently mutated, which we interpret to be suggestive of diverse alterations in function.

We focused on mutations that occur outside of the well-studied GAP domain, which we reasoned are likely directly interrupt RAS binding. Instead, we assessed the impact of mutations in the accessory domains. Functional understanding of the roles of these domains has recently been expanded with the observation that the C2 domain augments GAP activity, and we wished to better understand if the clusters of mutations within this domain all impact GAP activity in a similar fashion. We find this not to be the case, and that even within this functionally important domain there are varying impacts on catalytic activity. Similarly, our assessment of representative mutations within the SH2, SH3 and PH domains indicate varying impact upon catalytic activity and solubility, strongly suggestive of a range of functional outcomes.

Loss of expression will obviously alter RasGAP signaling, and the major classes of *RASA1* mutation that result in this outcome are the truncations and frame shifts. We did not analyze these as a complete loss of expressed protein may not be functionally informative on the normal signaling of RasGAP. In contrast, we find that one of the twelve selected mutations (A626E) results in a complete loss of soluble protein *in vitro*. We believe this is representative of a class of mutations that disrupt protein folding and /or solubility (i). Similarly, a significant number of our selected mutations result in destabilization of the expressed protein (ii). We used a reduction in melting temperature of 2 °C to delineate ‘destabilizing’ based on our statistical analyses. Although changes in melting temperature have been interpreted as conformational changes (e.g. for SHP2 phosphatase (55)) there is no evidence so far for conformational autoregulation for RasGAP, so we interpret these changes as destabilizing effects. While our cutoff is a somewhat arbitrary threshold, we believe it allows a categorization of mutants that are potentially of reduced solubility in cells, with consequent impacts on GAP activity towards Ras. In the context of a cell, these partially soluble mutants may, however, produce altered protein-protein interactions, localization, or changes in regulation of RasGAP, potential effects that will need to be explored in future studies.

Our Michaelis-Menten analysis of RasGAP mutants yielded an unexpected diversity of impact on the catalytic and Michaelis-Menten constants, kcat and KM. The kinetics of a majority of tested mutants (R194C, R283H, Y528C, R591C, H604N, W689R, S705F, and R707H) were disrupted via a decrease in kcat and /or increase (weakening) of KM (iii), leading to a reduced catalytic efficiency, an effect that will impact Ras signaling. In a cell, however, the functional effects of reduced catalytic rate may differ due to the presence of the membrane or other factors which can be explored in future studies.

Our analyses also show that loss of stability and alterations in kinetics are not mutually exclusive, and that some mutants (R194C, R283H, Y528C, W689R, and R707H) both reduced thermal stability and altered kinetics. This is suggestive that some clinically relevant mutants may result in impacts on signaling by more than one mechanism. Another population was of approximately wild type-level stability but low activity, including R591C, H604N, and S705F. These are the mutations for which changes in activity must necessarily arise due to factors other than destabilization. As previously discussed, R591 forms a salt bridge that mediates the interaction between the C2 and GAP domains, while H604 and S705 are located on the face of the C2 domain adjacent to the Ras binding site, suggesting their modes of action. By contrast, some of the mutants that we studied (R427Q, T612A, and H695L) are not significantly different from wild type in either solubility or kinetics. If these clinically observed mutants are functionally important, we hypothesize this to be due to other effects (iv), including the potential to interrupt interactions with binding partners, though we did not observe any effect on p190RhoGAP binding by R194C or R427Q. Further studies may be needed to reveal the mechanisms by which these effects occur. Lastly, there were no mutants with low stability and high activity, leading us to conclude that adequate stability is necessary but not sufficient for full GAP activity.

Overall, our study demonstrates that clinically associated mutations in the Ras GTPase Activating Protein, RasGAP, classify into groups that are dysregulated in a variety of ways. Through examination of a panel of representative mutants, these findings provide insight into the regulation of RasGAP in normal and disease states and allow for a better understanding of the molecular mechanisms underlying clinical presentations of cancer and vascular malformations linked with *RASA1* mutations.

## Materials and Methods

### Sequence Conservation Analysis

209 sequences of RasGAP were obtained and manually curated based on homology using the Basic Local Alignment Search Tool (BLAST). They were analyzed in Jalview v2.11.4.1 (56) and aligned using Clustal Omega (57). Sequence logos were created using WebLogo v2.8.2 (58). Conservation scores were obtained using the Consurf server (59) by entering the full-length primary amino acid sequence of p120RasGAP as the target sequence and the manual sequence alignment as the sequence alignment to be used.

### Protein Expression and Purification

All constructs were inserted into a modified pET-32 (Novagen) bacterial expression vector containing a hexahistidine (His6) tag followed by a TEV (tobacco etch virus) protease cleavage site using BamHI and XhoI restriction sites. Human RasGAP (p120RasGAP, UniProt ID: P20936) constructs containing residues 174-1047 (near full-length) and 174-444 (SH2-SH3-SH2) contain C236S, C261S, C372S, and C402S mutations to prevent unwanted disulfide bond formation *in vitro* as described in (33). Disease mutations were introduced using the QuikChange Lightning site-directed mutagenesis kit (Agilent) according to manufacturer’s instructions. H-Ras construct (UniProt ID: P01112) codes for residues 1-167 (wild type) of human *HRAS*. Proteins were expressed in Rosetta (*E. coli* DE3) cells grown in Luria Broth (LB) at 37°C in the presence of kanamycin and chloramphenicol to an optical density (OD600) of 0.7, then chilled to 18°C. Expression was induced by addition of 0.2 mM IPTG (isopropyl β-D-thiogalactopyranoside) and incubated for approximately eighteen hours at 18°C. Cultures were subsequently spun down at 4 °C for 30 min at 2,000 rcf and resuspended in 10 mL buffer containing 500 mM NaCl, 50 mM HEPES pH 8. Cells were lysed via addition of 50 µg/mL lysozyme and three successive freeze-thaw cycles followed by sonication. DNase I was added at a concentration of 10 U/nL. Lysate was clarified via centrifugation at 4 °C for 1 hr at 48,000 rcf. Supernatant was applied to a gravity column containing 1 mL Ni-NTA Agarose beads (Qiagen) and rocked at 4 °C for 1 hr. Flowthrough was collected, followed by washes containing successive concentrations of imidazole (20, 40, 100, 250, and 500 mM) in 500 mM NaCl and 50 mM HEPES pH 8.

For near full-length constructs, protein was then dialyzed at 4 °C overnight in 1 L buffer containing 150 mM NaCl and 20 mM Tris pH 8 using tubing with a 10 kDa cutoff. The protein was then diluted to a final salt concentration of 50 mM NaCl and anion exchange chromatography was performed using a 1 mL MonoQ column (GE Healthcare) in a Buffer A with 20 mM Tris pH 8.5 and a Buffer B containing 1 M NaCl, 20 mM Tris pH 8.5. A continuous NaCl gradient from 0-40% Buffer B was used. Relevant fractions were pooled and concentrated using an Amicon Ultra Centrifugal Filter, and size exclusion chromatography was performed using a Superdex 200 Increase 10/300 GL column (GE Healthcare) in a buffer containing 150 mM NaCl, 20 mM Tris pH 8. Relevant fractions were pooled and concentrated. To prepare protein for GAP assays, a stock was made via 1:1 dilution in 100% glycerol to form a 50% glycerol stock and stored at -20 °C. To prepare protein for thermal shift assays, stocks were flash frozen in liquid nitrogen and stored at -80 °C.

For RasGAP SH2-SH3-SH2 constructs and for H-Ras, His6-tagged TEV protease was added to cleave the N-terminal His6 tag and the protein was simultaneously dialyzed at 4 °C overnight in 1 L buffer containing 500 mM NaCl and 50 mM HEPES pH 8 using tubing with a 10 kDa cutoff to remove excess imidazole. The protein was then reapplied to the Ni-NTA Agarose column and rocked at 4 °C for 1 hr. Flowthrough was collected, followed by washes containing successive concentrations of imidazole (20, 40, 100, 250, and 500 mM) in 500 mM NaCl and 50 mM HEPES pH 8. Relevant fractions were pooled, concentrated, then diluted to a final salt concentration of 50 mM NaCl. Anion exchange chromatography was performed using a 1 mL ResourceQ column (Cytiva) in a Buffer A with 20 mM Tris pH 8.5 and a Buffer B containing 1 M NaCl, 20 mM Tris pH 8.5. A continuous NaCl gradient from 0-40% Buffer B was used. Relevant fractions were pooled and concentrated, and size exclusion chromatography was performed using a HiLoad Superdex 75 Prep 16/600 column (GE Healthcare) in a buffer containing 250 mM NaCl (for p120RasGAP SH2-SH3-SH2) or 150 mM NaCl (for H-Ras) and 20 mM Tris pH 8. Relevant fractions were pooled and concentrated.

### Expression and Solubility Tests

Expression of p120RasGAP ΔN A626E was carried out as described above in one liter of LB. A sample was taken from culture before induction with IPTG. A second sample was taken from culture after induction with 0.2mM IPTG and expression for approximately twenty hours at 18°C. Insoluble material in the pellet after centrifugation was resuspended in 6 M urea. Soluble sample in the supernatant was decanted after centrifugation. All samples were prepared in 1x Laemmli buffer (Bio-Rad) and boiled for five minutes. Samples were run on an SDS-PAGE gel (15% acrylamide) and stained with Coomassie blue dye. Bands were examined visually.

### Thermal Shift Assays

Wild type and mutants of RasGAP were prepared at a final concentration of 3 µM (for near full-length) or 10 µM (for SH2-SH3-SH2) with 5x SYPRO Orange dye (Invitrogen) in a buffer of 150 mM NaCl (for near full-length) or 250 mM NaCl (for SH2-SH3-SH2) and 20 mM Tris pH 8, with each reaction having a final volume of 25 µL. Four identical mixtures per construct were prepared per plate. Mixtures were prepared on ice in a white 96-well 0.2 mL capacity qPCR plate, sealed, and spun down for three minutes at 500 rcf at 4 °C to remove bubbles. The plate was then inserted into a Bio-Rad CFX Connect Real-Time PCR System operated by Bio-Rad CFX Manager software (v3.0). After an initial five-minute incubation at 4 °C, the temperature was increased up to 95 °C at a rate of one degree per minute, at which time a fluorescence reading was taken using the default SYBR/FAM filters. Data were exported and analyzed using GraphPad Prism v10.4.1. Individual traces of fluorescence vs. temperature were normalized and truncated after the peak of fluorescence was reached. These truncated traces were fit to a sigmoidal fit (4PL) and the temperature of the inflection point was taken as the melting temperature (Tm). Melting temperatures for the four mixtures were averaged to form one replicate. Three independent technical replicates were conducted for each construct, and differences from wild type were analyzed using ordinary one-way ANOVA with Dunnett’s test for multiple comparisons. To obtain the change in melting temperature relative to wild type (ΔTm), the average Tm taken from the wild type replicates was subtracted from each individual replicate from each construct.

### Peptide Synthesis

Synthetic peptide corresponding to residues 1083-1111 of p190RhoGAP (UniProt ID: Q9NRY4) was purchased from GenScript (Piscataway, NJ). The peptide has a length of 29 amino acids with the sequence DPSDpY(1087)AEPMDAVVKPRNEEENIpY(1105)SVPHDS, is phosphorylated at residues corresponding to p190RhoGAP pY1087 and pY1105 and has N-terminal acetylation and C-terminal amidation. Lyophilized peptide was reconstituted in water at 10 mM (35.1 mg/mL), flash frozen in liquid nitrogen in single-use aliquots, and stored at -80 °C.

### Isothermal Titration Calorimetry

Isothermal titration calorimetry was conducted using a Nano ITC Low Volume calorimeter (TA Instruments) as described in (33). Experiments were carried out at 25 °C in a buffer containing 250 mM NaCl, 20 mM Tris pH 8, and 1 mM TCEP (tris(2-carboxyethyl)phosphine). Proteins were purified as described above. Each RasGAP SH2-SH3-SH2 construct (wild type, R194C, and R427Q) was dialyzed using a Thermo Scientific Slide-A-Lyzer Dialysis Cassette (Extra Strength), 10,000 Da MWCO, 0.5-3mL capacity, Product #66380. p190RhoGAP peptide was dialyzed using a Spectrum Laboratories Spectra/Por Micro Float-A-Lyzer Dialysis Devices, MWCO 100-500 Da, Color: Green, Volume: 400-500 µL, Part Number F235061. Protein and peptide were dialyzed simultaneously in one liter of buffer at 4°C overnight. Concentrations were confirmed pre- and post-dialysis using a Thermo Scientific NanoDrop Lite Spectrophotometer at 280 nm with extinction coefficients of 45840 M^-1^cm^-1^ for all protein constructs and 404 M^-1^cm^-1^ for the peptide. Samples were then degassed in a vacuum desiccator for five minutes. The 182 µL sample cell was loaded with approximately 5 µM protein and the 50 µL syringe was loaded with approximately 25 µM peptide (see **Supplementary Table 5**). The instrument was operated using TA Instruments Nano ITCRun software (v3.7.0.0). After a 300 s baseline collection, injections of 2.5 µL each were added every 300 s for a total of twenty injections. Buffer-buffer and peptide-buffer controls were run to ensure that there were no anomalous dose-dependent heat effects. Experiments were analyzed using TA Instruments NanoAnalyze software (v3.11.0). Default baseline guides and integration regions were adjusted manually as necessary. The first injection was excluded from analysis for each replicate due to evaporation effects. Blank constant correction was applied to the obtained curves, which were then fit to the Independent model. Two to three technical replicates were performed per protein construct, and the averages and standard deviations reported.

### GTP Loading of H-Ras

Loading of Ras with GTP was conducted as described in (45, 54). Protein was incubated with 100-fold molar excess GTP and 10 mM EDTA (ethylenediaminetetraacetic acid) at 37 °C for 10 minutes. The mixture was returned to ice and spiked with 15 mM MgCl2, then centrifuged at 4 °C for 10 minutes at 16,900 rcf in a microcentrifuge tube. Size exclusion chromatography was then performed using a Superdex 75 10/300 GL column (GE Healthcare) in a buffer containing 10 mM EDTA and 20 mM Tris pH 8 in order to remove excess unbound nucleotide. Relevant fractions were pooled and concentrated.

To check the loading efficiency, a sample containing 10 nmol of protein was taken, boiled for 15 minutes, diluted to 1 mL in 20 mM Tris pH 8.5, then centrifuged at 4 °C for 10 minutes at 16,900 rcf. This sample, which now contained the free nucleotide released from the denatured and pelleted protein, was then subjected to anion exchange chromatography on a MonoQ column (GE Healthcare) in a Buffer A with 20 mM Tris pH 8.5 and a Buffer B containing 1 M NaCl, 20 mM Tris pH 8.5. A continuous gradient of 0-100% Buffer B was used to separate the GDP and GTP peaks. These peaks were then integrated, and the percentage of GTP was determined and used as the loading efficiency. This quotient was then applied to the total H-Ras concentration in order to determine the effective concentration of GTP loaded H-Ras. Typical efficiency ranges from 70-80%. Aliquots were flash frozen in liquid nitrogen and stored at -80 °C.

### GAP Assays

Assays were conducted as described in (45, 53). Briefly, reactions were conducted at 30 °C in a BioTek (Agilent) Synergy H1 Microplate Reader, along with the associated Gen5 software (v3.11.19) in fluorescence mode with an excitation wavelength of 430 nm, an emission wavelength of 450 nm, and bandwidths of 10 nm. Reactions were set up in black 384-well round-bottom microplates with a total reaction volume of 20 µL. Reactions contained 10 µM Phosphate Sensor (Invitrogen), varying concentrations of GTP loaded H-Ras, and varying concentrations of RasGAP in a final buffer of 12.5 mM NaCl, 20 mM Tris pH 8, 5 mM MgCl2, 2.5 mM EDTA, 1 mM TCEP (Tris(2-carboxyethyl)phosphine hydrochloride), and 0.01% Triton X-100. GTP loaded H-Ras concentrations varied between 10 and 150 µM to obtain Michaelis-Menten curves. Concentrations of RasGAP were different depending on the mutant of interest in order to optimize rates within the readable range, varying from 5 nM for wild type and highly active mutants to 20 nM for mutants with low activity. After an initial fluorescence baseline reading and incubation containing all components except GTP loaded H-Ras, the GTP loaded H-Ras was then added to initiate the reaction, and readings taken every twelve seconds for twenty minutes. In parallel, reactions with all components except pRasGAP were run to obtain rates for intrinsic H-Ras hydrolysis of GTP. Each reaction condition was performed in duplicate during each run, and five independent technical replicates were conducted for each RasGAP mutant.

Data were exported from Gen5 and processed in Microsoft Excel (v16.66.1) and Prism (v10.4.1) (GraphPad Software, MA). Time courses were converted from relative fluorescence units (RFU) to phosphate turnover in µM via a standard curve of inorganic phosphate (from 1mM Phosphate Standard, Millipore Sigma) fit to a single exponential equation. Duplicates conducted at the same time were analyzed together as one curve, and each replicate was analyzed separately. The initial rate from the linear portion of each time course was determined manually and the slope was determined via linear fit in units of µM/s. Rates for intrinsic hydrolysis were subtracted from the RasGAP-stimulated rates at corresponding H-Ras concentrations to account only for phosphate turnover arising from GAP activity. Corrected rates were graphed vs. substrate (GTP loaded H-Ras) concentration at five concentrations and fit to the Michaelis-Menten curve (called “kcat” in Prism) to extract kcat, KM, and kcat/KM. Statistical significance was determined via ordinary one-way ANOVA with Dunnett’s multiple comparisons test.

## Acknowledgments

We thank Amy Stiegler Wyler, Kimberly Vish, Nalini Natarajan, Karen Anderson, Enrique De La Cruz, Ben Turk, and Moitrayee Bhattacharyya for helpful discussions. We thank James Murphy for ITC training and support. M.E.P. supported by T32GM007324 and F31HL167578. This research was supported by NIH Grant R01NS117609 to T.J.B.

## Author Contributions

Conceptualization, Methodology, Writing: M.E.P., T.J.B.

Investigation: M.E.P., R.B.D., J.E.C.

Data Curation, Visualization: M.E.P.

Supervision: T.J.B.

