## Supplemental Information for "Clinically observed *RASA1* missense mutants exhibit diverse RasGAP protein behaviors"

**From:**

[1]Department of Molecular Biophysics and Biochemistry, Yale University, New Haven, CT, USA 06520

[2]Department of Pharmacology, Yale University, New Haven, CT, USA 06510

[3]Yale Cancer Center, Yale University, New Haven, CT, USA 06510

**Supplementary Table 1. *RASA1* missense mutants associated with vascular malformations.** Missense mutations in *RASA1* associated with vascular malformations compiled from the literature. Shown are the nucleotide and amino acid mutations reported, frequency of mutation, phenotype in patients, and source study.

| Codon Change | Protein Mutation | Single Letter Code | # Mutations at residue | Phenotype | Source |
|---|---|---|---|---|---|
| c.64G > T | p.Gly22Cys | G22C | 2 | Cutaneous CM, pial AVM, family history of termination of pregnancy at 7 months for VOGM | (1) |
| c.208G>C | p.Glu70Gln | E70Q | 1 | suspected HHT, lung AVM (VUS) | (2) |
| c.296C>T | p.Ala99Val | A99V | 4 | unilateral CMs, one patient had hypertrophic CM limb, another patient had a limb with CM that had a smaller circumference | (3) |
| c.346C>G | p.Leu116Val | L116V | 1 | CM on thigh, leg, foot, monocystic lymphatic malformation of the buttock, affected limb was hypertrophic and slightly longer (12mm) than other limb, dilated superficial veins | (3) |
| c.467G>T | p.Gly156Val | G156V | 1 | leg and thigh AVMs (VUS) | (2) |
| c.1277A > G | p.Tyr426Cys | Y426C | 8 | Cutaneous CM | (4) |
| c.1310T > G | p.Leu437Arg | L437R | 2 | Multifocal CMs on right palm, chest, and back, history of nosebleeds, son also had variant and multifocal CMs (VUS) | (5) |
| c.1352A>G | p.Asn451Ser | N451S | 1 | CMs on both les, legs had discrepant lengths, lymphedema on both feet | (3) |
| c.1583A > G | p.Tyr528Cys | Y528C | 4 | 1) Multiple blanching telangiectasias on palms, fingers, tongue, and toes, erythematous nasal mucosa with daily epistaxis, family history of epistaxis (suspected HHT), 2) Cutaneous CM | (1,6) |
| c.1589T > A | p.Val530Asp | V530D | 6 | Cutaneous CM, brainstem AVM (brain hemorrhage with facial palsy at 3 weeks), left leg AVM, basal cell carcinoma in 2 individuals | (1) |
| c.1619G > A | p.Cys540Tyr | C540Y | 9 | CM-AVM1 (CM, Nuchal CM, AVF) | (7) |
| c.1649A>T | p.Glu550Val | E550V | 1 | Multiple CMs (VUS) | (2) |
| p.1834A>G | p.Thr612Ala | T612A | 1 | Suspected HHT, brain AVM (VUS) | (2) |
| c.1877C > A | p.Ala626Glu | A626E | 1 | Cutaneous CM | (1) |
| c.2084A > T | p.His695Leu | H695L | 2 | Brain AVF, brain AVM, multifocal CMs on head/ face, mother (non-symptomatic) carries mutation | (5) |
| c.2119C > T | p.Arg707Cys | R707C | 4 | 1) VOGM, 2) Choroidal VOGM in two consecutive pregnancies of unaffected carrier mother | (8,9) |
| c.2209G>A | p.Val737Ile | V737I | 1 | Multiple CMs, telangiectasia (VUS) | (2) |
| c.2288A > T | p.Glu763Val | E763V | 3 | 1) Cutaneous CM, VOGM, heart failure, epilepsy, died rapidly after birth, 2) Cutaneous CM | (1,4) |
| c.2912T > C | p.Leu971Ser | L971S | 1 | VOGM | (8) |

**Supplementary Table 2. Cancer associated *RASA1* missense mutants.** Missense mutations in *RASA1* (associated with cancer and documented in cBioPortal on July 18, 2025). Redundancy is avoided by use of only a curated set of non-redundant studies in the analysis.

| Codon Change | Protein Mutation | Single Letter Code | # Mutations at residue | Cancer Type | Source |
|---|---|---|---|---|---|
| c.20G>T | p.Gly7Val | G7V | 1 | Lung Adenocarcinoma | (10) |
| c.30G>T | p.Glu10Asp | E10D | 1 | Lung Adenocarcinoma | (11) |
| c.32G>A | p.Gly11Asp | G11D | 1 | Colon Adenocarcinoma | (11) |
| c.34G>T | p.Gly12Cys | G12C | 2 | Lung Adenocarcinoma | (11) |
| c.34G>C | p.Gly12Arg | G12R | 2 | Non-Small Cell Lung Cancer | (11) |
| c.37C>T | p.Pro13Ser | P13S | 1 | Breast Invasive Cancer, NOS | (12) |
| c.40G>T | p.Val14Leu | V14L | 2 | Breast Invasive Ductal Carcinoma | (13) |
| c.40G>T | p.Val14Leu | V14L | 2 | Breast Invasive Ductal Carcinoma | (11) |
| c.59G>A | p.Gly20Glu | G20E | 3 | Renal Clear Cell Carcinoma | (13) |
| c.59G>A | p.Gly20Glu | G20E | 3 | Renal Clear Cell Carcinoma | (13) |
| c.59G>T | p.Gly20Val | G20V | 3 | Lung Adenocarcinoma | (11) |
| c.70G>A | p.Ala24Thr | A24T | 1 | Colorectal Adenocarcinoma | (14) |
| c.74C>T | p.Ala25Val | A25V | 1 | Pancreatic Adenocarcinoma | (15) |
| c.77G>A | p.Gly26Asp | G26D | 1 | Prostate Adenocarcinoma | (11) |
| c.92C>A | p.Pro31His | P31H | 1 | Colon Adenocarcinoma | (11) |
| c.98T>C | p.Val33Ala | V33A | 3 | Colon Adenocarcinoma | (11) |
| c.98T>C | p.Val33Ala | V33A | 3 | Rectal Adenocarcinoma | (11) |
| c.97G>C | p.Val33Leu | V33L | 3 | Invasive Breast Carcinoma | (11) |
| c.101G>C | p.Cys34Ser | C34S | 1 | Lung Adenocarcinoma | (11) |
| c.103C>G | p.Arg35Gly | R35G | 3 | Lung Squamous Cell Carcinoma | (16,17) |
| c.104G>T | p.Arg35Leu | R35L | 3 | Lung Adenocarcinoma | (11) |
| c.104G>T | p.Arg35Leu | R35L | 3 | Head and Neck Squamous Cell Carcinoma | (18-23) |
| c.107T>G | p.Val36Gly | V36G | 2 | B-Lymphoblastic Leukemia/Lymphoma | (24) |
| c.107T>G | p.Val36Gly | V36G | 2 | Optic nerve glioma | (24) |
| c.112A>G | p.Ile38Val | I38V | 1 | Head and Neck Squamous Cell Carcinoma of Unknown Primary | (13) |
| c.119C>G | p.Ala40Gly | A40G | 1 | Lung Squamous Cell Carcinoma | (18-23,25) |
| c.133G>A | p.Ala45Thr | A45T | 1 | Cholangiocarcinoma | (26) |
| c.142_143delinsTT | p.Pro48Phe | P48F | 2 | Skin Cancer, Non-Melanoma | (27) |
| c.143C>A | p.Pro48His | P48H | 2 | Fibrolamellar Carcinoma | (13) |
| c.149C>A | p.Pro50His | P50H | 1 | Colon Adenocarcinoma | (11) |
| c.170T>C | p.Val57Ala | V57A | 1 | Colon Adenocarcinoma | (28) |
| c.173C>T | p.Ala58Val | A58V | 1 | Colorectal Adenocarcinoma | (11) |
| c.184G>T | p.Gly62Cys | G62C | 2 | Lung Adenocarcinoma | (13) |
| c.184G>T | p.Gly62Cys | G62C | 2 | Lung Adenocarcinoma | (11) |
| c.193G>A | p.Ala65Thr | A65T | 1 | Uterine Carcinosarcoma/Uterine Malignant Mixed Mullerian Tumor | (29) |
| c.210G>T | p.Glu70Asp | E70D | 3 | Endometrial Carcinoma | (30) |
| c.209A>G | p.Glu70Gly | E70G | 3 | Activated B-cell Type | (31) |
| c.208G>C | p.Glu70Gln | E70Q | 3 | Breast Invasive Ductal Carcinoma | (32) |
| c.214C>A | p.Leu72Ile | L72I | 2 | Cutaneous Melanoma | (18-23) |
| c.214C>A | p.Leu72Ile | L72I | 2 | Colon Adenocarcinoma | (11) |
| c.224G>A | p.Gly75Glu | G75E | 1 | Pancreatic Neuroendocrine Tumor | (11) |
| c.229G>A | p.Val77Met | V77M | 1 | Colon Adenocarcinoma | (11) |
| c.238G>A | p.Ala80Thr | A80T | 1 | Colon Adenocarcinoma | (11) |
| c.244G>C | p.Gly82Arg | G82R | 1 | Prostate Adenocarcinoma | (11) |

| | | | | | |
|---|---|---|---|---|---|
| c.248G>A | p.Gly83Glu | G83E | 1 | Cholangiocarcinoma | (26) |
| c.266G>C | p.Gly89Ala | G89A | 2 | Prostate Adenocarcinoma | (11) |
| c.265G>A | p.Gly89Arg | G89R | 2 | Adrenocortical Carcinoma | (32) |
| c.290G>C | p.Gly97Ala | G97A | 1 | Colorectal Adenocarcinoma | (11) |
| c.296C>T | p.Ala99Val | A99V | 2 | Germinal Center B-Cell Type | (31) |
| c.296C>T | p.Ala99Val | A99V | 2 | Germinal Center B-Cell Type | (31) |
| c.319G>C | p.Val107Leu | V107L | 1 | Poorly Differentiated Non-Small Cell Lung Cancer | (11) |
| c.322G>T | p.Ala108Ser | A108S | 2 | Colon Adenocarcinoma | (13) |
| c.322G>T | p.Ala108Ser | A108S | 2 | Colon Adenocarcinoma | (11) |
| c.341T>C | p.Met114Thr | M114T | 1 | Uterine Mixed Endometrial Carcinoma | (18-23,25) |
| c.365C>T | p.Ser122Leu | S122L | 2 | Bladder Urothelial Carcinoma | (18-23) |
| c.365C>T | p.Ser122Leu | S122L | 2 | Breast Invasive Ductal Carcinoma | (18-23) |
| c.370C>A | p.Leu124Ile | L124I | 1 | Uterine Endometrioid Carcinoma | (18-23,25) |
| c.376G>A | p.Glu126Lys | E126K | 1 | Lung Squamous Cell Carcinoma | (33) |
| c.379A>T | p.Thr127Ser | T127S | 1 | Uterine Endometrioid Carcinoma | (18-23,25) |
| c.383T>C | p.Leu128Pro | L128P | 1 | Endometrial Carcinoma | (34) |
| c.385G>T | p.Gly129Trp | G129W | 1 | Cutaneous Melanoma | (18-23) |
| c.389C>T | p.Pro130Leu | P130L | 5 | Cutaneous Melanoma | (18-23) |
| c.388_389delinsTT | p.Pro130Leu | P130L | 5 | Melanoma | (33) |
| c.388C>T | p.Pro130Ser | P130S | 5 | Cutaneous Melanoma | (18-23) |
| c.388C>T | p.Pro130Ser | P130S | 5 | Rectal Adenocarcinoma | (11) |
| c.388C>T | p.Pro130Ser | P130S | 5 | Colorectal Adenocarcinoma | (14) |
| c.398G>A | p.Gly133Asp | G133D | 1 | Invasive Breast Carcinoma | (35) |
| c.404C>A | p.Pro135His | P135H | 3 | Large Cell Neuroendocrine Carcinoma | (11) |
| c.403C>T | p.Pro135Ser | P135S | 3 | Cutaneous Melanoma | (36) |
| c.403C>T | p.Pro135Ser | P135S | 3 | Cutaneous Squamous Cell Carcinoma | (13) |
| c.407C>G | p.Pro136Arg | P136R | 1 | Lung Adenocarcinoma | (11) |
| c.409C>G | p.Leu137Val | L137V | 3 | Bladder Urothelial Carcinoma | (32) |
| c.409C>G | p.Leu137Val | L137V | 3 | Breast Invasive Lobular Carcinoma | (13) |
| c.409C>G | p.Leu137Val | L137V | 3 | Breast Invasive Lobular Carcinoma | (11) |
| c.422C>T | p.Pro141Leu | P141L | 2 | Small Cell Lung Cancer | (37) |
| c.422C>T | p.Pro141Leu | P141L | 2 | Small Cell Lung Cancer | (37) |
| c.431C>A | p.Pro144His | P144H | 1 | Cutaneous Melanoma | (18-23) |
| c.437T>C | p.Leu146Ser | L146S | 2 | Colon Adenocarcinoma | (18-23,25) |
| c.437T>G | p.Leu146Trp | L146W | 2 | Melanoma | (38) |
| c.439G>A | p.Gly147Arg | G147R | 1 | Renal Clear Cell Carcinoma | (18-23,25) |
| c.458T>C | p.Val153Ala | V153A | 1 | Non-Small Cell Lung Cancer | (11) |
| c.462C>G | p.Asp154Glu | D154E | 3 | Bladder Urothelial Carcinoma | (39) |
| c.462C>G | p.Asp154Glu | D154E | 3 | Bladder Urothelial Carcinoma | (40) |
| c.461A>G | p.Asp154Gly | D154G | 3 | Esophagogastric Adenocarcinoma | (41) |
| c.466G>T | p.Gly156Cys | G156C | 1 | Non-Small Cell Lung Cancer | (11) |
| c.471C>G | p.Asp157Glu | D157E | 1 | Lung Adenocarcinoma | (11) |
| c.473C>G | p.Ser158Cys | S158C | 1 | Lung Adenocarcinoma | (11) |
| c.475C>G | p.Leu159Val | L159V | 2 | Lung Adenocarcinoma | (11) |
| c.475C>G | p.Leu159Val | L159V | 2 | Bladder Urothelial Carcinoma | (40) |
| c.489A>T | p.Glu163Asp | E163D | 1 | Cutaneous Melanoma | (32) |
| c.502G>A | p.Glu168Lys | E168K | 1 | Bladder Urothelial Carcinoma | (40) |
| c.508G>T | p.Ala170Ser | A170S | 3 | Lung Adenocarcinoma | (13) |
| c.508G>T | p.Ala170Ser | A170S | 3 | Lung Adenocarcinoma | (11) |
| c.509C>T | p.Ala170Val | A170V | 3 | Colon Adenocarcinoma | (11) |
| c.519G>T | p.Leu173Phe | L173F | 1 | Lung Adenocarcinoma | (11) |
| c.524C>G | p.Ala175Gly | A175G | 1 | Prostate Adenocarcinoma | (42) |
| c.527C>A | p.Pro176His | P176H | 3 | Colon Adenocarcinoma | (11) |
| c.526C>T | p.Pro176Ser | P176S | 3 | Invasive Breast Carcinoma | (13) |
| c.526C>T | p.Pro176Ser | P176S | 3 | Invasive Breast Carcinoma | (11) |

| | | | | | |
|---|---|---|---|---|---|
| c.550G>A | p.Gly184Arg | G184R | 1 | Breast Invasive Cancer, NOS | (12) |
| c.562A>G | p.Arg188Gly | R188G | 1 | Colon Adenocarcinoma | (11) |
| c.566C>T | p.Thr189Met | T189M | 1 | Uterine Endometrioid Carcinoma | (18-23,25) |
| c.580C>T | p.Arg194Cys | R194C | 11 | Astrocytoma | (18-23) |
| c.580C>T | p.Arg194Cys | R194C | 11 | Colon Adenocarcinoma | (18-23,25) |
| c.580C>T | p.Arg194Cys | R194C | 11 | Endometrial Carcinoma | (30) |
| c.580C>T | p.Arg194Cys | R194C | 11 | Glioblastoma Multiforme | (18-23,25) |
| c.580C>T | p.Arg194Cys | R194C | 11 | Uterine Endometrioid Carcinoma | (18-23,25) |
| c.581G>A | p.Arg194His | R194H | 11 | Colon Adenocarcinoma | (18-23,25) |
| c.581G>A | p.Arg194His | R194H | 11 | Rectal Adenocarcinoma | (11) |
| c.581G>A | p.Arg194His | R194H | 11 | Colon Adenocarcinoma | (11) |
| c.581G>A | p.Arg194His | R194H | 11 | Uterine Serous Carcinoma/Uterine Papillary Serous Carcinoma | (18-23,25) |
| c.581G>A | p.Arg194His | R194H | 11 | Uterine Endometrioid Carcinoma | (18-23,25) |
| c.581G>A | p.Arg194His | R194H | 11 | Cervical Squamous Cell Carcinoma | (18-23,25) |
| c.584T>A | p.Leu195His | L195H | 2 | Non-Small Cell Lung Cancer | (13) |
| c.584T>A | p.Leu195His | L195H | 2 | Non-Small Cell Lung Cancer | (11) |
| c.613C>T | p.Leu205Phe | L205F | 1 | Breast Invasive Lobular Carcinoma | (11) |
| c.616A>G | p.Ile206Val | I206V | 3 | Hepatocellular Carcinoma plus Intrahepatic Cholangiocarcinoma | (43) |
| c.616A>G | p.Ile206Val | I206V | 3 | Hepatocellular Carcinoma plus Intrahepatic Cholangiocarcinoma | (43) |
| c.616A>G | p.Ile206Val | I206V | 3 | Lung Squamous Cell Carcinoma | (11) |
| c.624G>C | p.Glu208Asp | E208D | 1 | Lung Adenocarcinoma | (11) |
| c.632G>T | p.Arg211Leu | R211L | 4 | Endometrial Carcinoma | (30) |
| c.632G>A | p.Arg211Gln | R211Q | 4 | Colon Adenocarcinoma | (13) |
| c.632G>A | p.Arg211Gln | R211Q | 4 | Colon Adenocarcinoma | (11) |
| c.631C>T | p.Arg211Trp | R211W | 4 | Lung Squamous Cell Carcinoma | (11) |
| c.635G>T | p.Arg212Met | R212M | 3 | Uterine Endometrioid Carcinoma | (18-23,25) |
| c.634A>T | p.Arg212Trp | R212W | 3 | Lung Adenocarcinoma | (13) |
| c.634A>T | p.Arg212Trp | R212W | 3 | Lung Adenocarcinoma | (11) |
| c.637C>T | p.Pro213Ser | P213S | 2 | Basaloid Penile Squamous Cell Carcinoma | (13) |
| c.637C>T | p.Pro213Ser | P213S | 2 | Colorectal Adenocarcinoma | (14) |
| c.652C>T | p.Leu218Phe | L218F | 1 | Colorectal Adenocarcinoma | (14) |
| c.656C>T | p.Ser219Leu | S219L | 1 | Bladder Urothelial Carcinoma | (18-23) |
| c.661C>G | p.Leu221Val | L221V | 1 | Lung Adenocarcinoma | (11) |
| c.665G>A | p.Ser222Asn | S222N | 1 | Colorectal Adenocarcinoma | (44) |
| c.672G>T | p.Met224Ile | M224I | 2 | Colon Adenocarcinoma | (11) |
| c.672G>T | p.Met224Ile | M224I | 2 | Uterine Endometrioid Carcinoma | (18-23,25) |
| c.677T>C | p.Val226Ala | V226A | 2 | Intrahepatic Cholangiocarcinoma | (13) |
| c.676G>C | p.Val226Leu | V226L | 2 | Lung Squamous Cell Carcinoma | (11) |
| c.685C>T | p.His229Tyr | H229Y | 1 | Uterine Endometrioid Carcinoma | (18-23,25) |
| c.704T>C | p.Met235Thr | M235T | 1 | Colon Adenocarcinoma | (11) |
| c.710G>C | p.Gly237Ala | G237A | 1 | Ampullary Carcinoma | (13) |
| c.725G>A | p.Gly242Asp | G242D | 1 | Colorectal Adenocarcinoma | (11) |
| c.727G>A | p.Gly243Arg | G243R | 1 | Prostate | (45) |
| c.733C>T | p.Arg245Cys | R245C | 6 | Lung Squamous Cell Carcinoma | (11) |
| c.733C>T | p.Arg245Cys | R245C | 6 | Tubular Stomach Adenocarcinoma | (18-23,25) |
| c.733C>G | p.Arg245Gly | R245G | 6 | Breast Invasive Ductal Carcinoma | (18-23) |
| c.734G>A | p.Arg245His | R245H | 6 | Cutaneous Melanoma | (46) |
| c.734G>A | p.Arg245His | R245H | 6 | Uterine Endometrioid Carcinoma | (13) |
| c.733C>A | p.Arg245Ser | R245S | 6 | Yolk Sac Tumor | (32) |
| c.740C>A | p.Ser247Tyr | S247Y | 2 | Breast Mixed Ductal and Lobular Carcinoma | (13) |
| c.740C>A | p.Ser247Tyr | S247Y | 2 | Breast Mixed Ductal and Lobular Carcinoma | (11) |
| c.751G>C | p.Asp251His | D251H | 1 | Breast Invasive Ductal Carcinoma | (18-23) |
| c.764A>T | p.Tyr255Phe | Y255F | 2 | Breast Invasive Ductal Carcinoma | (18-23) |

| | | | | | |
|---|---|---|---|---|---|
| c.763T>C | p.Tyr255His | Y255H | 2 | Endometrial Carcinoma | (30) |
| c.770G>A | p.Ser257Asn | S257N | 2 | Endometrial Carcinoma | (47) |
| c.769A>C | p.Ser257Arg | S257R | 2 | Lung Adenocarcinoma | (11) |
| c.779C>G | p.Ser260Cys | S260C | 1 | Lung Squamous Cell Carcinoma | (18-23,25) |
| c.787C>T | p.Leu263Phe | L263F | 1 | Hepatocellular Adenoma | (48) |
| c.823C>G | p.Pro275Ala | P275A | 1 | Glioblastoma | (49) |
| c.842G>T | p.Arg281Ile | R281I | 2 | Colon Adenocarcinoma | (18-23,25) |
| c.842G>T | p.Arg281Ile | R281I | 2 | Colorectal Adenocarcinoma | (33) |
| c.847C>T | p.Arg283Cys | R283C | 14 | Cholangiocarcinoma | (26) |
| c.847C>T | p.Arg283Cys | R283C | 14 | Colon Adenocarcinoma | (11) |
| c.847C>T | p.Arg283Cys | R283C | 14 | Colon Adenocarcinoma | (11) |
| c.847C>T | p.Arg283Cys | R283C | 14 | Lung Squamous Cell Carcinoma | (18-23,25) |
| c.848G>A | p.Arg283His | R283H | 14 | Colorectal Adenocarcinoma | (50) |
| c.848G>A | p.Arg283His | R283H | 14 | Colon Adenocarcinoma | (13) |
| c.848G>A | p.Arg283His | R283H | 14 | Rectal Adenocarcinoma | (13) |
| c.848G>A | p.Arg283His | R283H | 14 | Endometrial Carcinoma | (30) |
| c.848G>A | p.Arg283His | R283H | 14 | Colon Adenocarcinoma | (11) |
| c.848G>A | p.Arg283His | R283H | 14 | Rectal Adenocarcinoma | (11) |
| c.848G>A | p.Arg283His | R283H | 14 | Colon Adenocarcinoma | (11) |
| c.848G>A | p.Arg283His | R283H | 14 | Lung Adenocarcinoma | (51) |
| c.848G>A | p.Arg283His | R283H | 14 | Intestinal Type Stomach Adenocarcinoma | (18-23,25) |
| c.848G>A | p.Arg283His | R283H | 14 | Ampullary Carcinoma | (52) |
| c.853C>G | p.Arg285Gly | R285G | 5 | Lung Squamous Cell Carcinoma | (11) |
| c.854G>C | p.Arg285Pro | R285P | 5 | Lung Squamous Cell Carcinoma | (11) |
| c.854G>A | p.Arg285Gln | R285Q | 5 | Rectal Adenocarcinoma | (18-23,25) |
| c.854G>A | p.Arg285Gln | R285Q | 5 | Uterine Endometrioid Carcinoma | (18-23,25) |
| c.854G>A | p.Arg285Gln | R285Q | 5 | Uterine Endometrioid Carcinoma | (18-23,25) |
| c.866C>T | p.Pro289Leu | P289L | 1 | Skin Cancer, Non-Melanoma | (27) |
| c.869A>G | p.Tyr290Cys | Y290C | 2 | Prostate Adenocarcinoma | (11) |
| c.869A>C | p.Tyr290Ser | Y290S | 2 | Pancreatic Neuroendocrine Tumor | (11) |
| c.883G>A | p.Asp295Asn | D295N | 1 | Lung Adenocarcinoma | (13) |
| c.899G>T | p.Ser300Ile | S300I | 4 | Lung Adenosquamous Carcinoma | (16,17) |
| c.899G>T | p.Ser300Ile | S300I | 4 | Lung Adenosquamous Carcinoma | (16,17) |
| c.899G>T | p.Ser300Ile | S300I | 4 | Lung Adenosquamous Carcinoma | (16,17) |
| c.899G>T | p.Ser300Ile | S300I | 4 | Lung Adenosquamous Carcinoma | (16,17) |
| c.903C>A | p.Phe301Leu | F301L | 1 | Bladder Urothelial Carcinoma | (18-23) |
| c.906A>T | p.Leu302Phe | L302F | 1 | Lung Adenocarcinoma | (13) |
| c.915T>G | p.Asp305Glu | D305E | 1 | Mucinous Adenocarcinoma of the Colon and Rectum | (11) |
| c.921C>G | p.Phe307Leu | F307L | 2 | Breast Mixed Ductal and Lobular Carcinoma | (11) |
| c.919T>C | p.Phe307Leu | F307L | 2 | Rectal Adenocarcinoma | (11) |
| c.923T>A | p.Ile308Asn | I308N | 1 | Mucinous Adenocarcinoma of the Colon and Rectum | (11) |
| c.939A>T | p.Leu313Phe | L313F | 1 | Melanoma | (33) |
| c.940G>C | p.Glu314Gln | E314Q | 1 | Glioblastoma Multiforme | (13) |
| c.945T>A | p.Asp315Glu | D315E | 3 | Invasive Breast Carcinoma | (11) |
| c.943G>A | p.Asp315Asn | D315N | 3 | Cutaneous Melanoma | (13) |
| c.943G>A | p.Asp315Asn | D315N | 3 | Cutaneous Melanoma | (13) |
| c.957G>T | p.Trp319Cys | W319C | 1 | Colorectal Adenocarcinoma | (53) |
| c.964A>C | p.Asn322His | N322H | 4 | Mucinous Adenocarcinoma of the Appendix | (13) |
| c.964A>C | p.Asn322His | N322H | 4 | Uterine Endometrioid Carcinoma | (18-23,25) |
| c.966T>G | p.Asn322Lys | N322K | 4 | Colon Adenocarcinoma | (11) |
| c.965A>G | p.Asn322Ser | N322S | 4 | Lung Adenocarcinoma | (51) |
| c.967T>G | p.Leu323Val | L323V | 1 | Colon Adenocarcinoma | (11) |
| c.971G>A | p.Arg324Lys | R324K | 3 | Colon Adenocarcinoma | (11) |
| c.972A>T | p.Arg324Ser | R324S | 3 | Lung Adenocarcinoma | (13) |

| | | | | | |
|---|---|---|---|---|---|
| c.972A>T | p.Arg324Ser | R324S | 3 | Lung Adenocarcinoma | (11) |
| c.974C>G | p.Thr325Arg | T325R | 1 | Breast Invasive Ductal Carcinoma | (18-23) |
| c.976G>A | p.Asp326Asn | D326N | 2 | Hepatocellular Carcinoma | (18-23,25) |
| c.976G>T | p.Asp326Tyr | D326Y | 2 | Uterine Endometrioid Carcinoma | (18-23,25) |
| c.992T>C | p.Ile331Thr | I331T | 1 | Plasma Cell Myeloma | (54) |
| c.997G>C | p.Glu333Gln | E333Q | 1 | Glioblastoma Multiforme | (13) |
| c.1022G>A | p.Arg341Gln | R341Q | 2 | High-Grade Serous Ovarian Cancer | (13) |
| c.1021C>T | p.Arg341Trp | R341W | 2 | Lung Adenocarcinoma | (11) |
| c.1024G>A | p.Glu342Lys | E342K | 2 | Uterine Endometrioid Carcinoma | (13) |
| c.1024G>A | p.Glu342Lys | E342K | 2 | Non-Small Cell Lung Cancer | (11) |
| c.1043G>C | p.Gly348Ala | G348A | 2 | Cutaneous Squamous Cell Carcinoma | (13) |
| c.1042G>A | p.Gly348Arg | G348R | 2 | Uterine Mixed Endometrial Carcinoma | (18-23,25) |
| c.1072A>G | p.Lys358Glu | K358E | 2 | Lung Adenocarcinoma | (13) |
| c.1072A>G | p.Lys358Glu | K358E | 2 | Lung Adenocarcinoma | (11) |
| c.1087A>C | p.Asn363His | N363H | 1 | Hepatocellular Carcinoma plus Intrahepatic Cholangiocarcinoma | (43) |
| c.1098G>C | p.Met366Ile | M366I | 1 | Esophageal Adenocarcinoma | (33) |
| c.1100C>A | p.Thr367Lys | T367K | 1 | Non-Small Cell Lung Cancer | (11) |
| c.1102G>T | p.Val368Phe | V368F | 2 | Cutaneous Melanoma | (18-23) |
| c.1102G>A | p.Val368Ile | V368I | 2 | Lung Squamous Cell Carcinoma | (18-23,25) |
| c.1105G>A | p.Gly369Ser | G369S | 1 | Leiomyosarcoma | (18-23,25) |
| c.1115G>T | p.Cys372Phe | C372F | 1 | Cholangiocarcinoma | (26) |
| c.1123C>G | p.Leu375Val | L375V | 1 | Bladder Urothelial Carcinoma | (18-23) |
| c.1133C>T | p.Pro378Leu | P378L | 1 | Endometrial Carcinoma | (30) |
| c.1136C>T | p.Ser379Leu | S379L | 3 | Non-Small Cell Lung Cancer | (11) |
| c.1136C>T | p.Ser379Leu | S379L | 3 | Lung Adenocarcinoma | (11) |
| c.1136C>T | p.Ser379Leu | S379L | 3 | Lung Adenocarcinoma | (11) |
| c.1139A>G | p.Asp380Gly | D380G | 2 | Paraganglioma | (13) |
| c.1138G>A | p.Asp380Asn | D380N | 2 | Skin | (55-58) |
| c.1145C>G | p.Thr382Ser | T382S | 1 | Uterine Clear Cell Carcinoma | (59) |
| c.1151G>A | p.Gly384Asp | G384D | 1 | Colon Adenocarcinoma | (11) |
| c.1156T>C | p.Tyr386His | Y386H | 1 | Colon Adenocarcinoma | (11) |
| c.1160C>T | p.Ser387Leu | S387L | 1 | Endocervical Adenocarcinoma | (13) |
| c.1163T>G | p.Leu388Arg | L388R | 1 | Lung Squamous Cell Carcinoma | (11) |
| c.1171C>T | p.Arg391Trp | R391W | 7 | Esophageal Adenocarcinoma | (18-23) |
| c.1171C>T | p.Arg391Trp | R391W | 7 | Colon Adenocarcinoma | (13) |
| c.1171C>T | p.Arg391Trp | R391W | 7 | Endometrial Carcinoma | (30) |
| c.1171C>T | p.Arg391Trp | R391W | 7 | Colon Adenocarcinoma | (11) |
| c.1171C>T | p.Arg391Trp | R391W | 7 | Colon Adenocarcinoma | (11) |
| c.1171C>T | p.Arg391Trp | R391W | 7 | Endometrial Carcinoma | (34) |
| c.1171C>T | p.Arg391Trp | R391W | 7 | Colorectal Adenocarcinoma | (14) |
| c.1190A>C | p.Gln397Pro | Q397P | 1 | Invasive Breast Carcinoma | (11) |
| c.1193G>T | p.Arg398Leu | R398L | 8 | Colorectal Adenocarcinoma | (11) |
| c.1193G>A | p.Arg398Gln | R398Q | 8 | Colon Adenocarcinoma | (13) |
| c.1193G>A | p.Arg398Gln | R398Q | 8 | Upper Tract Urothelial Carcinoma | (13) |
| c.1193G>A | p.Arg398Gln | R398Q | 8 | Uterine Endometrioid Carcinoma | (13) |
| c.1193G>A | p.Arg398Gln | R398Q | 8 | Colon Adenocarcinoma | (11) |
| c.1193G>A | p.Arg398Gln | R398Q | 8 | Non-Small Cell Lung Cancer | (11) |
| c.1193G>A | p.Arg398Gln | R398Q | 8 | Uterine Endometrioid Carcinoma | (18-23,25) |
| c.1193G>A | p.Arg398Gln | R398Q | 8 | Uterine Carcinosarcoma/Uterine Malignant Mixed Mullerian Tumor | (29) |
| c.1207C>G | p.Pro403Ala | P403A | 3 | Lung Squamous Cell Carcinoma | (18-23,25) |
| c.1208C>T | p.Pro403Leu | P403L | 3 | Intestinal Type Stomach Adenocarcinoma | (18-23,25) |
| c.1208C>A | p.Pro403Gln | P403Q | 3 | Lung Adenocarcinoma | (11) |
| c.1210A>T | p.Thr404Ser | T404S | 2 | Papillary Renal Cell Carcinoma | (18-23) |
| c.1210A>T | p.Thr404Ser | T404S | 2 | Papillary Renal Cell Carcinoma | (33) |

| | | | | | |
|---|---|---|---|---|---|
| c.1214C>G | p.Pro405Arg | P405R | 1 | Prostate Adenocarcinoma | (11) |
| c.1224G>T | p.Gln408His | Q408H | 1 | Uterine Endometrioid Carcinoma | (13) |
| c.1230G>A | p.Met410Ile | M410I | 1 | Leiomyosarcoma | (18-23,25) |
| c.1234G>A | p.Gly412Arg | G412R | 2 | Spindle Cell Carcinoma of the Lung | (13) |
| c.1234G>A | p.Gly412Arg | G412R | 2 | Spindle Cell Carcinoma of the Lung | (11) |
| c.1241G>T | p.Arg414Leu | R414L | 7 | Lung Adenocarcinoma | (11) |
| c.1241G>A | p.Arg414Gln | R414Q | 7 | Cutaneous Squamous Cell Carcinoma | (13) |
| c.1241G>A | p.Arg414Gln | R414Q | 7 | Lung Adenocarcinoma | (13) |
| c.1241G>A | p.Arg414Gln | R414Q | 7 | Lung Adenocarcinoma | (11) |
| c.1241G>A | p.Arg414Gln | R414Q | 7 | Uterine Endometrioid Carcinoma | (18-23,25) |
| c.1240C>T | p.Arg414Trp | R414W | 7 | Medullary Carcinoma of the Colon | (11) |
| c.1240C>T | p.Arg414Trp | R414W | 7 | Colon Adenocarcinoma | (11) |
| c.1244A>T | p.Tyr415Phe | Y415F | 1 | Poorly Differentiated Thyroid Cancer | (13) |
| c.1251C>A | p.Asn417Lys | N417K | 1 | Lung Adenocarcinoma | (11) |
| c.1253G>C | p.Ser418Thr | S418T | 1 | Lung Adenocarcinoma | (18-23) |
| c.1262A>G | p.Asp421Gly | D421G | 2 | Upper Tract Urothelial Carcinoma | (60) |
| c.1262A>G | p.Asp421Gly | D421G | 2 | Upper Tract Urothelial Carcinoma | (13) |
| c.1268T>C | p.Ile423Thr | I423T | 1 | Hepatocellular Carcinoma | (61) |
| c.1272T>G | p.Asp424Glu | D424E | 2 | Hepatocellular Carcinoma | (61) |
| c.1270G>T | p.Asp424Tyr | D424Y | 2 | Colon Adenocarcinoma | (11) |
| c.1273C>A | p.His425Asn | H425N | 2 | Lung Adenosquamous Carcinoma | (11) |
| c.1273C>T | p.His425Tyr | H425Y | 2 | Lung Adenocarcinoma | (62) |
| c.1277A>G | p.Tyr426Cys | Y426C | 2 | Endometrial Carcinoma | (47) |
| c.1276T>C | p.Tyr426His | Y426H | 2 | Uterine Clear Cell Carcinoma | (13) |
| c.1280G>A | p.Arg427Gln | R427Q | 13 | Endometrial Carcinoma | (47) |
| c.1280G>A | p.Arg427Gln | R427Q | 13 | Mucinous Adenocarcinoma of the Colon and Rectum | (18-23,25) |
| c.1280G>A | p.Arg427Gln | R427Q | 13 | Colon Adenocarcinoma | (63) |
| c.1280G>A | p.Arg427Gln | R427Q | 13 | Colon Adenocarcinoma | (13) |
| c.1280G>A | p.Arg427Gln | R427Q | 13 | Basaloid Penile Squamous Cell Carcinoma | (13) |
| c.1280G>A | p.Arg427Gln | R427Q | 13 | Uterine Endometrioid Carcinoma | (13) |
| c.1280G>A | p.Arg427Gln | R427Q | 13 | Mucinous Adenocarcinoma of the Colon and Rectum | (33) |
| c.1280G>A | p.Arg427Gln | R427Q | 13 | Colon Adenocarcinoma | (11) |
| c.1280G>A | p.Arg427Gln | R427Q | 13 | Non-Small Cell Lung Cancer | (11) |
| c.1280G>A | p.Arg427Gln | R427Q | 13 | Uterine Endometrioid Carcinoma | (18-23,25) |
| c.1280G>A | p.Arg427Gln | R427Q | 13 | Uterine Endometrioid Carcinoma | (18-23,25) |
| c.1280G>A | p.Arg427Gln | R427Q | 13 | Uterine Endometrioid Carcinoma | (18-23,25) |
| c.1280G>A | p.Arg427Gln | R427Q | 13 | Colorectal Adenocarcinoma | (14) |
| c.1292T>C | p.Ile431Thr | I431T | 2 | Uterine Endometrioid Carcinoma | (13) |
| c.1292T>C | p.Ile431Thr | I431T | 2 | Endometrial Carcinoma | (30) |
| c.1301G>A | p.Gly434Glu | G434E | 3 | Renal Clear Cell Carcinoma | (13) |
| c.1301G>A | p.Gly434Glu | G434E | 3 | Non-Small Cell Lung Cancer | (11) |
| c.1301G>T | p.Gly434Val | G434V | 3 | Rectal Adenocarcinoma | (18-23,25) |
| c.1325C>A | p.Pro442Gln | P442Q | 4 | Cutaneous Melanoma | (18-23) |
| c.1325C>A | p.Pro442Gln | P442Q | 4 | Cutaneous Melanoma | (18-23) |
| c.1325C>A | p.Pro442Gln | P442Q | 4 | Cutaneous Melanoma | (18-23) |
| c.1324C>T | p.Pro442Ser | P442S | 4 | Uterine Carcinosarcoma/Uterine Malignant Mixed Mullerian Tumor | (18-23) |
| c.1332G>T | p.Gln444His | Q444H | 1 | Cutaneous Melanoma | (18-23) |
| c.1336C>G | p.Gln446Glu | Q446E | 1 | Head and Neck Squamous Cell Carcinoma | (18-23) |
| c.1343A>G | p.Gln448Arg | Q448R | 1 | Cutaneous Melanoma | (13) |
| c.1345G>T | p.Val449Leu | V449L | 1 | Cervical Squamous Cell Carcinoma | (18-23,25) |
| c.1352A>G | p.Asn451Ser | N451S | 1 | Osteosarcoma | (64) |
| c.1360G>A | p.Val454Met | V454M | 1 | Lung Squamous Cell Carcinoma | (11) |
| c.1363G>T | p.Asp455Tyr | D455Y | 1 | Esophageal Squamous Cell Carcinoma | (18-23) |

| | | | | | |
|---|---|---|---|---|---|
| c.1370A>G | p.Lys457Arg | K457R | 1 | Renal Clear Cell Carcinoma | (65) |
| c.1376T>C | p.Ile459Thr | I459T | 1 | Uterine Clear Cell Carcinoma | (13) |
| c.1390C>T | p.Arg464Cys | R464C | 6 | Endometrial Carcinoma | (30) |
| c.1390C>T | p.Arg464Cys | R464C | 6 | Breast Invasive Carcinoma, NOS | (11) |
| c.1390C>T | p.Arg464Cys | R464C | 6 | Intestinal Type Stomach Adenocarcinoma | (18-23,25) |
| c.1391G>A | p.Arg464His | R464H | 6 | Colon Adenocarcinoma | (11) |
| c.1391G>A | p.Arg464His | R464H | 6 | Colorectal Adenocarcinoma | (14) |
| c.1391G>A | p.Arg464His | R464H | 6 | Colorectal Adenocarcinoma | (14) |
| c.1394G>A | p.Arg465His | R465H | 2 | Colon Adenocarcinoma | (18-23,25) |
| c.1394G>A | p.Arg465His | R465H | 2 | Prostate Adenocarcinoma | (42) |
| c.1400C>G | p.Thr467Arg | T467R | 2 | Breast Invasive Ductal Carcinoma | (13) |
| c.1400C>G | p.Thr467Arg | T467R | 2 | Breast Invasive Ductal Carcinoma | (11) |
| c.1402A>G | p.Lys468Glu | K468E | 4 | Lung Adenocarcinoma | (11) |
| c.1404G>C | p.Lys468Asn | K468N | 4 | Cutaneous Melanoma | (18-23) |
| c.1404G>T | p.Lys468Asn | K468N | 4 | Colon Adenocarcinoma | (18-23,25) |
| c.1402A>C | p.Lys468Gln | K468Q | 4 | Uterine Endometrioid Carcinoma | (18-23,25) |
| c.1405G>A | p.Asp469Asn | D469N | 1 | Skin Cancer, Non-Melanoma | (27) |
| c.1409C>T | p.Ala470Val | A470V | 1 | Glioblastoma | (49) |
| c.1441C>T | p.Leu481Phe | L481F | 3 | Cervical Squamous Cell Carcinoma | (18-23,25) |
| c.1442T>C | p.Leu481Pro | L481P | 3 | Adenoid Cystic Breast Cancer | (66) |
| c.1442T>C | p.Leu481Pro | L481P | 3 | Adenoid Cystic Breast Cancer | (67) |
| c.1453G>T | p.Gly485Cys | G485C | 2 | Colon Adenocarcinoma | (13) |
| c.1453G>T | p.Gly485Cys | G485C | 2 | Colon Adenocarcinoma | (11) |
| c.1460G>A | p.Gly487Glu | G487E | 1 | Prostate Adenocarcinoma | (11) |
| c.1465C>T | p.Arg489Cys | R489C | 3 | Oligoastrocytoma | (18-23) |
| c.1465C>T | p.Arg489Cys | R489C | 3 | Colon Adenocarcinoma | (11) |
| c.1465C>T | p.Arg489Cys | R489C | 3 | Lung Adenocarcinoma | (18-23) |
| c.1471A>G | p.Lys491Glu | K491E | 3 | Rectal Adenocarcinoma | (18-23,25) |
| c.1472A>C | p.Lys491Thr | K491T | 3 | Prostate Adenocarcinoma | (13) |
| c.1472A>C | p.Lys491Thr | K491T | 3 | Prostate Adenocarcinoma | (11) |
| c.1474A>G | p.Asn492Asp | N492D | 1 | Rectal Adenocarcinoma | (11) |
| c.1480T>C | p.Tyr494His | Y494H | 2 | Colon Adenocarcinoma | (13) |
| c.1480T>C | p.Tyr494His | Y494H | 2 | Colon Adenocarcinoma | (11) |
| c.1494G>T | p.Glu498Asp | E498D | 2 | Lung Squamous Cell Carcinoma | (18-23,25) |
| c.1494G>T | p.Glu498Asp | E498D | 2 | Uterine Endometrioid Carcinoma | (18-23,25) |
| c.1502A>G | p.Asp501Gly | D501G | 2 | Papillary Renal Cell Carcinoma | (18-23) |
| c.1501G>T | p.Asp501Tyr | D501Y | 2 | Colon Adenocarcinoma | (11) |
| c.1511T>C | p.Leu504Pro | L504P | 1 | Intrahepatic Cholangiocarcinoma | (68) |
| c.1523A>T | p.Glu508Val | E508V | 1 | Thymoma | (18-23) |
| c.1526G>A | p.Ser509Asn | S509N | 2 | Cutaneous Melanoma | (18-23) |
| c.1526G>A | p.Ser509Asn | S509N | 2 | Melanoma | (33) |
| c.1528G>A | p.Glu510Lys | E510K | 1 | Cutaneous Melanoma | (36) |
| c.1535G>A | p.Arg512Gln | R512Q | 3 | Colon Adenocarcinoma | (13) |
| c.1535G>A | p.Arg512Gln | R512Q | 3 | Colon Adenocarcinoma | (11) |
| c.1535G>A | p.Arg512Gln | R512Q | 3 | Colon Adenocarcinoma | (11) |
| c.1561G>T | p.Asp521Tyr | D521Y | 1 | Uterine Endometrioid Carcinoma | (18-23,25) |
| c.1564C>T | p.Leu522Phe | L522F | 2 | Colorectal Adenocarcinoma | (14) |
| c.1565T>A | p.Leu522His | L522H | 2 | Lung Adenocarcinoma | (11) |
| c.1567A>G | p.Ser523Gly | S523G | 2 | Melanoma | (69) |
| c.1568G>A | p.Ser523Asn | S523N | 2 | Colon Adenocarcinoma | (11) |
| c.1571T>C | p.Val524Ala | V524A | 1 | Colon Adenocarcinoma | (11) |
| c.1573T>C | p.Cys525Arg | C525R | 1 | Lung Adenocarcinoma | (11) |
| c.1577C>A | p.Ser526Tyr | S526Y | 2 | Colorectal Adenocarcinoma | (13) |
| c.1577C>A | p.Ser526Tyr | S526Y | 2 | Colon Adenocarcinoma | (11) |
| c.1582T>C | p.Tyr528His | Y528H | 1 | Intestinal Type Stomach Adenocarcinoma | (18-23,25) |

| | | | | | |
|---|---|---|---|---|---|
| c.1588G>A | p.Val530Ile | V530I | 1 | Undifferentiated Pleomorphic Sarcoma/Malignant Fibrous Histiocytoma/High-Grade Spindle Cell Sarcoma | (13) |
| c.1592A>G | p.His531Arg | H531R | 1 | Endometrial Carcinoma | (47) |
| c.1600C>A | p.Leu534Ile | L534I | 1 | Colon Adenocarcinoma | (18-23,25) |
| c.1604T>C | p.Phe535Ser | F535S | 1 | Small Cell Lung Cancer | (70) |
| c.1607G>A | p.Gly536Asp | G536D | 1 | Prostate Adenocarcinoma | (11) |
| c.1609A>G | p.Arg537Gly | R537G | 3 | Lung Adenocarcinoma | (11) |
| c.1610G>T | p.Arg537Met | R537M | 3 | Lung Squamous Cell Carcinoma | (13) |
| c.1610G>T | p.Arg537Met | R537M | 3 | Lung Squamous Cell Carcinoma | (11) |
| c.1612C>T | p.Pro538Ser | P538S | 1 | Pancreatic Neuroendocrine Tumor | (11) |
| c.1622T>G | p.Phe541Cys | F541C | 1 | Intrahepatic Cholangiocarcinoma | (13) |
| c.1624C>G | p.Gln542Glu | Q542E | 1 | Lung Adenosquamous Carcinoma | (13) |
| c.1629A>G | p.Ile543Met | I543M | 1 | Prostate | (45) |
| c.1653A>C | p.Glu551Asp | E551D | 3 | Colorectal Adenocarcinoma | (13) |
| c.1653A>C | p.Glu551Asp | E551D | 3 | Colon Adenocarcinoma | (11) |
| c.1652A>T | p.Glu551Val | E551V | 3 | Breast Invasive Ductal Carcinoma | (11) |
| c.1662C>G | p.Ile554Met | I554M | 2 | Lung Adenocarcinoma | (13) |
| c.1662C>G | p.Ile554Met | I554M | 2 | Lung Adenocarcinoma | (11) |
| c.1663T>C | p.Phe555Leu | F555L | 1 | Glioblastoma Multiforme | (13) |
| c.1667A>G | p.Tyr556Cys | Y556C | 1 | Colon Adenocarcinoma | (11) |
| c.1684C>G | p.Pro562Ala | P562A | 1 | Lung Adenocarcinoma | (11) |
| c.1690C>A | p.Gln564Lys | Q564K | 2 | Colon Adenocarcinoma | (11) |
| c.1691A>G | p.Gln564Arg | Q564R | 2 | Hepatocellular Carcinoma | (61) |
| c.1714C>A | p.Leu572Met | L572M | 1 | Esophagogastric Adenocarcinoma | (41) |
| c.1720G>A | p.Ala574Thr | A574T | 2 | Lung Adenocarcinoma | (13) |
| c.1720G>A | p.Ala574Thr | A574T | 2 | Lung Adenocarcinoma | (11) |
| c.1736G>A | p.Arg579Gln | R579Q | 10 | Upper Tract Urothelial Carcinoma | (71) |
| c.1736G>A | p.Arg579Gln | R579Q | 10 | Upper Tract Urothelial Carcinoma | (71) |
| c.1736G>A | p.Arg579Gln | R579Q | 10 | Upper Tract Urothelial Carcinoma | (71) |
| c.1736G>A | p.Arg579Gln | R579Q | 10 | Upper Tract Urothelial Carcinoma | (71) |
| c.1736G>A | p.Arg579Gln | R579Q | 10 | Upper Tract Urothelial Carcinoma | (71) |
| c.1736G>A | p.Arg579Gln | R579Q | 10 | Upper Tract Urothelial Carcinoma | (71) |
| c.1735C>T | p.Arg579Trp | R579W | 10 | Colon Adenocarcinoma | (13) |
| c.1735C>T | p.Arg579Trp | R579W | 10 | Adrenocortical Carcinoma | (13) |
| c.1735C>T | p.Arg579Trp | R579W | 10 | Desmoplastic Small-Round-Cell Tumor | (13) |
| c.1735C>T | p.Arg579Trp | R579W | 10 | Endometrial Carcinoma | (30) |
| c.1740A>C | p.Lys580Asn | K580N | 2 | Colon Adenocarcinoma | (13) |
| c.1740A>C | p.Lys580Asn | K580N | 2 | Colon Adenocarcinoma | (11) |
| c.1756T>C | p.Ser586Pro | S586P | 1 | Colon Adenocarcinoma | (11) |
| c.1760A>G | p.Asn587Ser | N587S | 2 | Colorectal Adenocarcinoma | (13) |
| c.1760A>G | p.Asn587Ser | N587S | 2 | Rectal Adenocarcinoma | (11) |
| c.1765C>T | p.Arg589Cys | R589C | 7 | Endometrial Carcinoma | (47) |
| c.1765C>T | p.Arg589Cys | R589C | 7 | Esophageal Adenocarcinoma | (33) |
| c.1766G>A | p.Arg589His | R589H | 7 | Colon Adenocarcinoma | (28) |
| c.1766G>A | p.Arg589His | R589H | 7 | Colon Adenocarcinoma | (11) |
| c.1766G>A | p.Arg589His | R589H | 7 | Prostate Adenocarcinoma | (11) |
| c.1766G>A | p.Arg589His | R589H | 7 | Colon Adenocarcinoma | (11) |
| c.1766G>A | p.Arg589His | R589H | 7 | Uterine Endometrioid Carcinoma | (18-23,25) |
| c.1771C>T | p.Arg591Cys | R591C | 13 | Ganglioglioma | (72) |
| c.1771C>T | p.Arg591Cys | R591C | 13 | Angiosarcoma | (73) |
| c.1771C>T | p.Arg591Cys | R591C | 13 | Colon Adenocarcinoma | (63) |
| c.1771C>T | p.Arg591Cys | R591C | 13 | Angiosarcoma | (73) |
| c.1771C>T | p.Arg591Cys | R591C | 13 | Uterine Endometrioid Carcinoma | (13) |
| c.1771C>T | p.Arg591Cys | R591C | 13 | Colon Adenocarcinoma | (11) |

| | | | | | |
|---|---|---|---|---|---|
| c.1771C>T | p.Arg591Cys | R591C | 13 | Glioblastoma Multiforme | (18-23,25) |
| c.1771C>T | p.Arg591Cys | R591C | 13 | Uterine Endometrioid Carcinoma | (18-23,25) |
| c.1771C>T | p.Arg591Cys | R591C | 13 | Stomach Adenocarcinoma | (18-23,25) |
| c.1771C>G | p.Arg591Gly | R591G | 13 | Hepatocellular Carcinoma | (18-23,25) |
| c.1772G>A | p.Arg591His | R591H | 13 | Angiosarcoma | (13) |
| c.1772G>A | p.Arg591His | R591H | 13 | Colon Adenocarcinoma | (11) |
| c.1772G>A | p.Arg591His | R591H | 13 | Diffuse Type Stomach Adenocarcinoma | (18-23,25) |
| c.1774C>G | p.Gln592Glu | Q592E | 1 | Lung Adenocarcinoma | (74) |
| c.1778T>G | p.Val593Gly | V593G | 2 | Pancreatic Adenocarcinoma | (13) |
| c.1778T>G | p.Val593Gly | V593G | 2 | Pancreatic Adenocarcinoma | (11) |
| c.1786C>A | p.Leu596Ile | L596I | 1 | Colon Adenocarcinoma | (11) |
| c.1793T>C | p.Leu598Ser | L598S | 1 | Esophageal Adenocarcinoma | (18-23) |
| c.1799T>A | p.Ile600Asn | I600N | 1 | Lung Adenocarcinoma | (18-23) |
| c.1806A>C | p.Glu602Asp | E602D | 3 | Cancer of Unknown Primary | (13) |
| c.1806A>C | p.Glu602Asp | E602D | 3 | Colon Adenocarcinoma | (11) |
| c.1804G>A | p.Glu602Lys | E602K | 3 | Colon Adenocarcinoma | (11) |
| c.1810C>A | p.His604Asn | H604N | 1 | Uterine Endometrioid Carcinoma | (75) |
| c.1817T>A | p.Leu606His | L606H | 1 | Head and Neck Squamous Cell Carcinoma | (18-23) |
| c.1819C>T | p.Pro607Ser | P607S | 1 | Pancreatic Adenocarcinoma | (11) |
| c.1825A>C | p.Lys609Gln | K609Q | 1 | Rectal Adenocarcinoma | (18-23,25) |
| c.1837A>G | p.Asn613Asp | N613D | 1 | Colon Adenocarcinoma | (11) |
| c.1840C>T | p.Pro614Ser | P614S | 1 | Bladder Urothelial Carcinoma | (18-23) |
| c.1846T>C | p.Cys616Arg | C616R | 3 | Endometrial Carcinoma | (30) |
| c.1846T>C | p.Cys616Arg | C616R | 3 | Lung Squamous Cell Carcinoma | (11) |
| c.1847G>A | p.Cys616Tyr | C616Y | 3 | Colorectal Adenocarcinoma | (11) |
| c.1858C>A | p.Leu620Met | L620M | 1 | Colon Adenocarcinoma | (11) |
| c.1864A>G | p.Ser622Gly | S622G | 1 | Chondroblastic Osteosarcoma | (24) |
| c.1873G>T | p.Val625Leu | V625L | 1 | Uterine Endometrioid Carcinoma | (18-23,25) |
| c.1881A>C | p.Lys627Asn | K627N | 4 | Hepatocellular Carcinoma | (33) |
| c.1881A>C | p.Lys627Asn | K627N | 4 | Colon Adenocarcinoma | (11) |
| c.1880A>G | p.Lys627Arg | K627R | 4 | Hepatocellular Carcinoma | (61) |
| c.1880A>C | p.Lys627Thr | K627T | 4 | Uterine Endometrioid Carcinoma | (18-23,25) |
| c.1907C>A | p.Pro636Gln | P636Q | 1 | Lung Squamous Cell Carcinoma | (18-23,25) |
| c.1918G>A | p.Glu640Lys | E640K | 3 | Cutaneous Melanoma | (76) |
| c.1918G>A | p.Glu640Lys | E640K | 3 | Cutaneous Melanoma | (62) |
| c.1918G>A | p.Glu640Lys | E640K | 3 | Breast Invasive Ductal Carcinoma | (11) |
| c.1921G>C | p.Glu641Gln | E641Q | 1 | Esophageal Squamous Cell Carcinoma | (77) |
| c.1935T>G | p.Asp645Glu | D645E | 1 | Lung Adenocarcinoma | (11) |
| c.1936G>A | p.Asp646Asn | D646N | 2 | Lung Adenocarcinoma | (13) |
| c.1936G>A | p.Asp646Asn | D646N | 2 | Lung Adenocarcinoma | (11) |
| c.1939C>T | p.Leu647Phe | L647F | 2 | Colon Adenocarcinoma | (63) |
| c.1939C>G | p.Leu647Val | L647V | 2 | Lung Adenocarcinoma | (11) |
| c.1952T>C | p.Ile651Thr | I651T | 2 | Rectal Adenocarcinoma | (11) |
| c.1952T>C | p.Ile651Thr | I651T | 2 | Colon Adenocarcinoma | (11) |
| c.1958G>T | p.Arg653Ile | R653I | 1 | Colon Adenocarcinoma | (11) |
| c.1968A>G | p.Ile656Met | I656M | 1 | Uterine Endometrioid Carcinoma | (13) |
| c.1980T>G | p.Asn660Lys | N660K | 2 | Rectal Adenocarcinoma | (11) |
| c.1978A>T | p.Asn660Tyr | N660Y | 2 | Intrahepatic Cholangiocarcinoma | (68) |
| c.1983A>C | p.Lys661Asn | K661N | 1 | Breast Invasive Ductal Carcinoma | (13) |
| c.1985C>A | p.Thr662Lys | T662K | 2 | Neuroblastoma | (13) |
| c.1984A>C | p.Thr662Pro | T662P | 2 | Colon Adenocarcinoma | (11) |
| c.1989G>T | p.Lys663Asn | K663N | 2 | Colon Adenocarcinoma | (63) |
| c.1987A>C | p.Lys663Gln | K663Q | 2 | Colon Adenocarcinoma | (11) |
| c.2005G>A | p.Asp669Asn | D669N | 1 | Stomach Adenocarcinoma | (18-23,25) |
| c.2013A>C | p.Leu671Phe | L671F | 1 | Prostate Adenocarcinoma | (42) |
| c.2017A>C | p.Met673Leu | M673L | 2 | Pancreatic Adenocarcinoma | (78) |

| | | | | | |
|---|---|---|---|---|---|
| c.2017A>C | p.Met673Leu | M673L | 2 | Pancreatic Adenocarcinoma | (33) |
| c.2020C>T | p.Arg674Cys | R674C | 7 | Prostate Adenocarcinoma | (18-23,25) |
| c.2020C>T | p.Arg674Cys | R674C | 7 | Colon Adenocarcinoma | (11) |
| c.2021G>A | p.Arg674His | R674H | 7 | Colon Adenocarcinoma | (13) |
| c.2021G>A | p.Arg674His | R674H | 7 | Colon Adenocarcinoma | (11) |
| c.2021G>A | p.Arg674His | R674H | 7 | Non-Small Cell Lung Cancer | (11) |
| c.2021G>A | p.Arg674His | R674H | 7 | Intestinal Type Stomach Adenocarcinoma | (18-23,25) |
| c.2021G>C | p.Arg674Pro | R674P | 7 | Uterine Serous Carcinoma/Uterine Papillary Serous Carcinoma | (18-23,25) |
| c.2026C>A | p.Gln676Lys | Q676K | 1 | Lung Adenocarcinoma | (62) |
| c.2036G>A | p.Arg679Gln | R679Q | 2 | Colon Adenocarcinoma | (11) |
| c.2036G>A | p.Arg679Gln | R679Q | 2 | Uterine Endometrioid Carcinoma | (18-23,25) |
| c.2039T>C | p.Leu680Ser | L680S | 1 | Uterine Serous Carcinoma/Uterine Papillary Serous Carcinoma | (18-23,25) |
| c.2054C>G | p.Ala685Gly | A685G | 2 | Uterine Endometrioid Carcinoma | (33) |
| c.2054C>G | p.Ala685Gly | A685G | 2 | Uterine Endometrioid Carcinoma | (18-23,25) |
| c.2056A>G | p.Thr686Ala | T686A | 1 | Rectal Adenocarcinoma | (11) |
| c.2065T>A | p.Trp689Arg | W689R | 1 | Lung Carcinoma | (79) |
| c.2069T>G | p.Phe690Cys | F690C | 2 | Breast Invasive Lobular Carcinoma | (13) |
| c.2069T>G | p.Phe690Cys | F690C | 2 | Breast Invasive Lobular Carcinoma | (11) |
| c.2071C>G | p.Leu691Val | L691V | 1 | Diffuse Type Stomach Adenocarcinoma | (18-23,25) |
| c.2074C>G | p.Leu692Val | L692V | 1 | Bladder Urothelial Carcinoma | (18-23) |
| c.2088A>G | p.Ile696Met | I696M | 3 | Endometrial Carcinoma | (30) |
| c.2087T>C | p.Ile696Thr | I696T | 3 | Colon Adenocarcinoma | (11) |
| c.2086A>G | p.Ile696Val | I696V | 3 | Head and Neck Squamous Cell Carcinoma | (18-23) |
| c.2098G>T | p.Gly700Cys | G700C | 4 | Cutaneous Melanoma | (18-23) |
| c.2098G>T | p.Gly700Cys | G700C | 4 | Serous Ovarian Cancer | (18-23) |
| c.2098G>T | p.Gly700Cys | G700C | 4 | Serous Ovarian Cancer | (33) |
| c.2098G>T | p.Gly700Cys | G700C | 4 | Uterine Serous Carcinoma/Uterine Papillary Serous Carcinoma | (18-23,25) |
| c.2101A>G | p.Ile701Val | I701V | 1 | Endometrial Carcinoma | (30) |
| c.2114C>G | p.Ser705Cys | S705C | 2 | Lung Adenocarcinoma | (80) |
| c.2113T>C | p.Ser705Pro | S705P | 2 | Pancreatic Adenocarcinoma | (18-23) |
| c.2119C>T | p.Arg707Cys | R707C | 9 | Colon Adenocarcinoma | (11) |
| c.2119C>T | p.Arg707Cys | R707C | 9 | Colon Adenocarcinoma | (11) |
| c.2119C>T | p.Arg707Cys | R707C | 9 | Colorectal Adenocarcinoma | (11) |
| c.2120G>A | p.Arg707His | R707H | 9 | Colon Adenocarcinoma | (81) |
| c.2120G>A | p.Arg707His | R707H | 9 | Upper Tract Urothelial Carcinoma | (13) |
| c.2120G>A | p.Arg707His | R707H | 9 | Colorectal Adenocarcinoma | (14) |
| c.2120G>A | p.Arg707His | R707H | 9 | Colorectal Adenocarcinoma | (14) |
| c.2120G>T | p.Arg707Leu | R707L | 9 | Non-Small Cell Lung Cancer | (11) |
| c.2120G>C | p.Arg707Pro | R707P | 9 | Uterine Serous Carcinoma/Uterine Papillary Serous Carcinoma | (18-23,25) |
| c.2126G>A | p.Arg709Gln | R709Q | 2 | Cholangiocarcinoma | (26) |
| c.2126G>A | p.Arg709Gln | R709Q | 2 | Colorectal Adenocarcinoma | (14) |
| c.2131C>G | p.Arg711Gly | R711G | 8 | Endometrial Carcinoma | (30) |
| c.2131C>G | p.Arg711Gly | R711G | 8 | Endometrial Carcinoma | (34) |
| c.2132G>T | p.Arg711Leu | R711L | 8 | Non-Small Cell Lung Cancer | (11) |
| c.2132G>A | p.Arg711Gln | R711Q | 8 | Endometrial Carcinoma | (47) |
| c.2132G>A | p.Arg711Gln | R711Q | 8 | Colon Adenocarcinoma | (63) |
| c.2132G>A | p.Arg711Gln | R711Q | 8 | Colon Adenocarcinoma | (63) |
| c.2132G>A | p.Arg711Gln | R711Q | 8 | Colon Adenocarcinoma | (11) |
| c.2132G>A | p.Arg711Gln | R711Q | 8 | Colorectal Adenocarcinoma | (14) |
| c.2135A>G | p.Tyr712Cys | Y712C | 3 | Uterine Carcinosarcoma/Uterine Malignant Mixed Mullerian Tumor | (29) |
| c.2134T>G | p.Tyr712Asp | Y712D | 3 | Stomach Adenocarcinoma | (82) |

| | | | | | |
|---|---|---|---|---|---|
| c.2134T>G | p.Tyr712Asp | Y712D | 3 | Stomach Adenocarcinoma | (83) |
| c.2138C>A | p.Ser713Tyr | S713Y | 1 | Cutaneous Melanoma | (18-23) |
| c.2141T>C | p.Met714Thr | M714T | 1 | Hepatocellular Carcinoma | (61) |
| c.2144A>G | p.Glu715Gly | E715G | 1 | Non-Small Cell Lung Cancer | (11) |
| c.2149A>T | p.Ile717Phe | I717F | 1 | Rectal Adenocarcinoma | (11) |
| c.2154G>C | p.Met718Ile | M718I | 1 | Lung Squamous Cell Carcinoma | (18-23,25) |
| c.2155C>A | p.Pro719Thr | P719T | 1 | Colon Adenocarcinoma | (11) |
| c.2165A>C | p.Glu722Ala | E722A | 1 | Hepatocellular Carcinoma | (18-23,25) |
| c.2168A>G | p.Tyr723Cys | Y723C | 1 | Rectal Adenocarcinoma | (11) |
| c.2199G>T | p.Lys733Asn | K733N | 1 | Colon Adenocarcinoma | (11) |
| c.2225C>T | p.Ser742Leu | S742L | 2 | Glioblastoma Multiforme | (13) |
| c.2225C>T | p.Ser742Leu | S742L | 2 | Cutaneous Squamous Cell Carcinoma | (13) |
| c.2234G>A | p.Cys745Tyr | C745Y | 2 | Endometrial Carcinoma | (30) |
| c.2234G>A | p.Cys745Tyr | C745Y | 2 | Colon Adenocarcinoma | (11) |
| c.2246G>A | p.Arg749Gln | R749Q | 10 | Colon Adenocarcinoma | (18-23,25) |
| c.2246G>A | p.Arg749Gln | R749Q | 10 | Cancer of Unknown Primary | (13) |
| c.2246G>A | p.Arg749Gln | R749Q | 10 | Colon Adenocarcinoma | (11) |
| c.2246G>A | p.Arg749Gln | R749Q | 10 | Colon Adenocarcinoma | (11) |
| c.2246G>A | p.Arg749Gln | R749Q | 10 | Colon Adenocarcinoma | (11) |
| c.2246G>A | p.Arg749Gln | R749Q | 10 | Colon Adenocarcinoma | (11) |
| c.2246G>A | p.Arg749Gln | R749Q | 10 | Uterine Endometrioid Carcinoma | (18-23,25) |
| c.2246G>A | p.Arg749Gln | R749Q | 10 | Uterine Endometrioid Carcinoma | (18-23,25) |
| c.2246G>A | p.Arg749Gln | R749Q | 10 | Uterine Endometrioid Carcinoma | (18-23,25) |
| c.2246G>A | p.Arg749Gln | R749Q | 10 | Cervical Squamous Cell Carcinoma | (18-23,25) |
| c.2255T>A | p.Leu752Gln | L752Q | 1 | Breast Invasive Ductal Carcinoma | (12) |
| c.2264T>C | p.Ile755Thr | I755T | 1 | Colon Adenocarcinoma | (18-23,25) |
| c.2266C>A | p.Leu756Ile | L756I | 1 | Colorectal Adenocarcinoma | (53) |
| c.2274G>T | p.Arg758Ser | R758S | 2 | Lung Adenocarcinoma | (11) |
| c.2274G>T | p.Arg758Ser | R758S | 2 | Rectal Adenocarcinoma | (11) |
| c.2281C>T | p.Leu761Phe | L761F | 2 | Uterine Endometrioid Carcinoma | (18-23,25) |
| c.2282T>C | p.Leu761Pro | L761P | 2 | Colon Adenocarcinoma | (11) |
| c.2288A>G | p.Glu763Gly | E763G | 8 | Colon Adenocarcinoma | (13) |
| c.2288A>G | p.Glu763Gly | E763G | 8 | Colon Adenocarcinoma | (11) |
| c.2287G>A | p.Glu763Lys | E763K | 8 | Colon Adenocarcinoma | (28) |
| c.2287G>A | p.Glu763Lys | E763K | 8 | Mucinous Adenocarcinoma of the Colon and Rectum | (18-23,25) |
| c.2287G>A | p.Glu763Lys | E763K | 8 | Mucinous Adenocarcinoma of the Colon and Rectum | (33) |
| c.2287G>A | p.Glu763Lys | E763K | 8 | Lung Adenocarcinoma | (11) |
| c.2287G>A | p.Glu763Lys | E763K | 8 | Uterine Endometrioid Carcinoma | (18-23,25) |
| c.2287G>A | p.Glu763Lys | E763K | 8 | Ampullary Carcinoma | (52) |
| c.2293C>T | p.Leu765Phe | L765F | 2 | Cutaneous Melanoma | (76) |
| c.2293C>T | p.Leu765Phe | L765F | 2 | Cutaneous Melanoma | (62) |
| c.2300C>T | p.Ser767Leu | S767L | 1 | Colorectal Adenocarcinoma | (14) |
| c.2309T>C | p.Leu770Ser | L770S | 1 | Uterine Clear Cell Carcinoma | (13) |
| c.2323G>C | p.Asp775His | D775H | 1 | Breast Invasive Ductal Carcinoma | (11) |
| c.2331A>C | p.Glu777Asp | E777D | 2 | Colorectal Adenocarcinoma | (13) |
| c.2331A>C | p.Glu777Asp | E777D | 2 | Colon Adenocarcinoma | (11) |
| c.2344G>T | p.Asp782Tyr | D782Y | 2 | Colorectal Adenocarcinoma | (11) |
| c.2344G>T | p.Asp782Tyr | D782Y | 2 | Lung Squamous Cell Carcinoma | (18-23,25) |
| c.2348A>C | p.Glu783Ala | E783A | 2 | Lung Squamous Cell Carcinoma | (18-23,25) |
| c.2347G>A | p.Glu783Lys | E783K | 2 | Esophageal Adenocarcinoma | (84) |
| c.2350G>A | p.Ala784Thr | A784T | 2 | Colon Adenocarcinoma | (11) |
| c.2351C>T | p.Ala784Val | A784V | 2 | Colon Adenocarcinoma | (11) |
| c.2366G>T | p.Arg789Leu | R789L | 9 | Clear Cell Ovarian Cancer | (81) |
| c.2366G>T | p.Arg789Leu | R789L | 9 | Head and Neck Squamous Cell Carcinoma | (18-23) |

| | | | | | |
|---|---|---|---|---|---|
| c.2366G>C | p.Arg789Pro | R789P | 9 | Cutaneous Melanoma | (13) |
| c.2366G>C | p.Arg789Pro | R789P | 9 | Breast Invasive Ductal Carcinoma | (11) |
| c.2366G>C | p.Arg789Pro | R789P | 9 | Endometrial Carcinoma | (34) |
| c.2366G>A | p.Arg789Gln | R789Q | 9 | Lung Adenosquamous Carcinoma | (13) |
| c.2366G>A | p.Arg789Gln | R789Q | 9 | Uterine Endometrioid Carcinoma | (13) |
| c.2366G>A | p.Arg789Gln | R789Q | 9 | Colorectal Adenocarcinoma | (11) |
| c.2366G>A | p.Arg789Gln | R789Q | 9 | Head and Neck Squamous Cell Carcinoma | (18-23) |
| c.2372C>T | p.Thr791Ile | T791I | 1 | Endometrial Carcinoma | (30) |
| c.2374A>G | p.Thr792Ala | T792A | 2 | Uterine Endometrioid Carcinoma | (18-23,25) |
| c.2375C>T | p.Thr792Ile | T792I | 2 | Colon Adenocarcinoma | (11) |
| c.2393T>A | p.Met798Lys | M798K | 2 | Lung Adenocarcinoma | (11) |
| c.2392A>T | p.Met798Leu | M798L | 2 | Invasive Breast Carcinoma | (11) |
| c.2406G>A | p.Met802Ile | M802I | 4 | Lung Squamous Cell Carcinoma | (33) |
| c.2406G>A | p.Met802Ile | M802I | 4 | Lung Squamous Cell Carcinoma | (18-23,25) |
| c.2406G>T | p.Met802Ile | M802I | 4 | Uterine Endometrioid Carcinoma | (18-23,25) |
| c.2406G>A | p.Met802Ile | M802I | 4 | Cervical Squamous Cell Carcinoma | (18-23,25) |
| c.2411C>A | p.Ala804Asp | A804D | 1 | Serous Ovarian Cancer | (33) |
| c.2434C>T | p.His812Tyr | H812Y | 2 | Renal Clear Cell Carcinoma | (18-23,25) |
| c.2434C>T | p.His812Tyr | H812Y | 2 | Renal Clear Cell Carcinoma | (33) |
| c.2438C>A | p.Ala813Asp | A813D | 1 | Lung Squamous Cell Carcinoma | (11) |
| c.2443A>G | p.Lys815Glu | K815E | 1 | Bladder Urothelial Carcinoma | (71) |
| c.2449T>C | p.Ser817Pro | S817P | 1 | Prostate Adenocarcinoma | (11) |
| c.2455T>G | p.Leu819Val | L819V | 1 | Basaloid Penile Squamous Cell Carcinoma | (13) |
| c.2460G>T | p.Lys820Asn | K820N | 1 | Colorectal Adenocarcinoma | (85) |
| c.2466G>T | p.Met822Ile | M822I | 1 | Cutaneous Melanoma | (18-23) |
| c.2478G>T | p.Gln826His | Q826H | 1 | Uterine Endometrioid Carcinoma | (18-23,25) |
| c.2480C>T | p.Ser827Phe | S827F | 2 | Basaloid Penile Squamous Cell Carcinoma | (13) |
| c.2479T>C | p.Ser827Pro | S827P | 2 | Colon Adenocarcinoma | (11) |
| c.2492G>A | p.Ser831Asn | S831N | 2 | Pancreatic Adenocarcinoma | (11) |
| c.2492G>C | p.Ser831Thr | S831T | 2 | Renal Clear Cell Carcinoma | (65) |
| c.2512A>C | p.Asn838His | N838H | 2 | Breast Invasive Ductal Carcinoma | (11) |
| c.2514T>A | p.Asn838Lys | N838K | 2 | Lung Adenocarcinoma | (11) |
| c.2515G>A | p.Glu839Lys | E839K | 2 | Chromophobe Renal Cell Carcinoma | (18-23,25) |
| c.2515G>A | p.Glu839Lys | E839K | 2 | Chromophobe Renal Cell Carcinoma | (33) |
| c.2521G>A | p.Val841Met | V841M | 1 | Colon Adenocarcinoma | (11) |
| c.2526C>A | p.Asn842Lys | N842K | 1 | Hepatocellular Adenoma | (48) |
| c.2528C>A | p.Thr843Asn | T843N | 1 | Oligodendroglioma | (18-23) |
| c.2530A>G | p.Asn844Asp | N844D | 1 | Endometrial Carcinoma | (30) |
| c.2536A>G | p.Thr846Ala | T846A | 3 | Bladder Urothelial Carcinoma | (71) |
| c.2536A>G | p.Thr846Ala | T846A | 3 | Upper Tract Urothelial Carcinoma | (71) |
| c.2536A>G | p.Thr846Ala | T846A | 3 | Upper Tract Urothelial Carcinoma | (60) |
| c.2541C>A | p.His847Gln | H847Q | 1 | Colon Adenocarcinoma | (11) |
| c.2543T>C | p.Leu848Pro | L848P | 3 | Esophageal Adenocarcinoma | (18-23) |
| c.2543T>A | p.Leu848Gln | L848Q | 3 | Hepatocellular Carcinoma | (86) |
| c.2543T>A | p.Leu848Gln | L848Q | 3 | Hepatocellular Carcinoma | (61) |
| c.2546T>C | p.Leu849Ser | L849S | 1 | Papillary Thyroid Cancer | (18-23,25) |
| c.2549A>G | p.Asn850Ser | N850S | 4 | Bladder Urothelial Carcinoma | (71) |
| c.2549A>G | p.Asn850Ser | N850S | 4 | Upper Tract Urothelial Carcinoma | (71) |
| c.2549A>G | p.Asn850Ser | N850S | 4 | Upper Tract Urothelial Carcinoma | (60) |
| c.2549A>G | p.Asn850Ser | N850S | 4 | Small Cell Lung Cancer | (87) |
| c.2552T>C | p.Ile851Thr | I851T | 1 | Colon Adenocarcinoma | (11) |
| c.2558C>T | p.Ser853Leu | S853L | 4 | Lung Adenocarcinoma | (13) |
| c.2558C>T | p.Ser853Leu | S853L | 4 | Lung Adenocarcinoma | (11) |
| c.2558C>T | p.Ser853Leu | S853L | 4 | Lung Adenocarcinoma | (11) |
| c.2558C>T | p.Ser853Leu | S853L | 4 | Lung Adenocarcinoma | (11) |
| c.2564T>C | p.Leu855Pro | L855P | 5 | Lung Squamous Cell Carcinoma | (88) |

| | | | | | |
|---|---|---|---|---|---|
| c.2564T>C | p.Leu855Pro | L855P | 5 | Lung Squamous Cell Carcinoma | (88) |
| c.2564T>C | p.Leu855Pro | L855P | 5 | Lung Squamous Cell Carcinoma | (88) |
| c.2564T>C | p.Leu855Pro | L855P | 5 | Lung Squamous Cell Carcinoma | (88) |
| c.2564T>G | p.Leu855Arg | L855R | 5 | Colon Adenocarcinoma | (63) |
| c.2571G>T | p.Glu857Asp | E857D | 2 | Uterine Endometrioid Carcinoma | (13) |
| c.2571G>C | p.Glu857Asp | E857D | 2 | Bladder Urothelial Carcinoma | (18-23) |
| c.2580C>A | p.Phe860Leu | F860L | 1 | Colon Adenocarcinoma | (11) |
| c.2592A>C | p.Glu864Asp | E864D | 1 | Uterine Endometrioid Carcinoma | (18-23,25) |
| c.2603C>T | p.Pro868Leu | P868L | 2 | Head and Neck Squamous Cell Carcinoma | (18-23) |
| c.2603C>T | p.Pro868Leu | P868L | 2 | Head and Neck Squamous Cell Carcinoma | (18-23) |
| c.2610G>T | p.Leu870Phe | L870F | 2 | Rectal Adenocarcinoma | (11) |
| c.2609T>G | p.Leu870Trp | L870W | 2 | Breast Invasive Ductal Carcinoma | (89) |
| c.2612G>T | p.Arg871Ile | R871I | 2 | Lung Squamous Cell Carcinoma | (11) |
| c.2612G>A | p.Arg871Lys | R871K | 2 | Uterine Endometrioid Carcinoma | (18-23,25) |
| c.2624G>C | p.Gly875Ala | G875A | 3 | Papillary Thyroid Cancer | (18-23,25) |
| c.2624G>A | p.Gly875Glu | G875E | 3 | Mucinous Adenocarcinoma of the Colon and Rectum | (13) |
| c.2624G>A | p.Gly875Glu | G875E | 3 | Lung Squamous Cell Carcinoma | (18-23,25) |
| c.2634G>T | p.Gln878His | Q878H | 1 | Uterine Endometrioid Carcinoma | (18-23,25) |
| c.2637A>C | p.Lys879Asn | K879N | 1 | Colorectal Adenocarcinoma | (11) |
| c.2639C>A | p.Ser880Tyr | S880Y | 1 | Basaloid Penile Squamous Cell Carcinoma | (13) |
| c.2641G>C | p.Val881Leu | V881L | 3 | Colon Adenocarcinoma | (28) |
| c.2641G>C | p.Val881Leu | V881L | 3 | Lung Squamous Cell Carcinoma | (11) |
| c.2641G>C | p.Val881Leu | V881L | 3 | Lung Adenocarcinoma | (18-23) |
| c.2651A>G | p.Lys884Arg | K884R | 1 | Breast Invasive Ductal Carcinoma | (12) |
| c.2655G>T | p.Trp885Cys | W885C | 1 | Cancer of Unknown Primary | (13) |
| c.2663A>C | p.Asn888Thr | N888T | 1 | Uterine Endometrioid Carcinoma | (18-23,25) |
| c.2669_2670delinsTT | p.Thr890Ile | T890I | 2 | Cancer of Unknown Primary | (13) |
| c.2669_2670delinsTT | p.Thr890Ile | T890I | 2 | Cancer of Unknown Primary | (13) |
| c.2671A>G | p.Met891Val | M891V | 1 | Lung Adenocarcinoma | (90) |
| c.2681G>T | p.Arg894Ile | R894I | 2 | Colon Adenocarcinoma | (11) |
| c.2681G>A | p.Arg894Lys | R894K | 2 | Colorectal Adenocarcinoma | (85) |
| c.2696T>C | p.Phe899Ser | F899S | 1 | Rectal Adenocarcinoma | (11) |
| c.2703T>A | p.Phe901Leu | F901L | 1 | Colon Adenocarcinoma | (11) |
| c.2705T>C | p.Leu902Pro | L902P | 3 | Lung Adenocarcinoma | (13) |
| c.2705T>C | p.Leu902Pro | L902P | 3 | Lung Adenocarcinoma | (11) |
| c.2704C>G | p.Leu902Val | L902V | 3 | Serous Ovarian Cancer | (18-23) |
| c.2708G>A | p.Arg903Gln | R903Q | 2 | Cholangiocarcinoma | (26) |
| c.2708G>A | p.Arg903Gln | R903Q | 2 | Uterine Endometrioid Carcinoma | (18-23,25) |
| c.2711T>G | p.Leu904Arg | L904R | 1 | Colorectal Adenocarcinoma | (50) |
| c.2714T>A | p.Ile905Asn | I905N | 1 | Lung Squamous Cell Carcinoma | (18-23,25) |
| c.2718T>G | p.Cys906Trp | C906W | 1 | Endometrial Carcinoma | (30) |
| c.2719C>T | p.Pro907Ser | P907S | 2 | Lung Adenocarcinoma | (90) |
| c.2719C>T | p.Pro907Ser | P907S | 2 | Non-Small Cell Lung Cancer | (11) |
| c.2722G>A | p.Ala908Thr | A908T | 2 | Colon Adenocarcinoma | (11) |
| c.2723C>T | p.Ala908Val | A908V | 2 | Cutaneous Squamous Cell Carcinoma | (91) |
| c.2729T>G | p.Leu910Arg | L910R | 2 | Cutaneous Melanoma | (18-23) |
| c.2729T>G | p.Leu910Arg | L910R | 2 | Lung Adenocarcinoma | (81) |
| c.2738G>A | p.Arg913Gln | R913Q | 4 | Cutaneous Melanoma | (18-23) |
| c.2738G>A | p.Arg913Gln | R913Q | 4 | Colon Adenocarcinoma | (11) |
| c.2738G>A | p.Arg913Gln | R913Q | 4 | Mucinous Adenocarcinoma of the Colon and Rectum | (11) |
| c.2737C>T | p.Arg913Trp | R913W | 4 | Gastrointestinal Stromal Tumor | (13) |
| c.2742G>A | p.Met914Ile | M914I | 1 | Plasma Cell Myeloma | (54) |

| | | | | | |
|---|---|---|---|---|---|
| c.2744T>G | p.Phe915Cys | F915C | 3 | Rectal Adenocarcinoma | (18-23,25) |
| c.2745C>A | p.Phe915Leu | F915L | 3 | Serous Ovarian Cancer | (18-23) |
| c.2743T>C | p.Phe915Leu | F915L | 3 | Uterine Endometrioid Carcinoma | (18-23,25) |
| c.2756C>T | p.Ser919Leu | S919L | 3 | Lung Adenocarcinoma | (13) |
| c.2756C>T | p.Ser919Leu | S919L | 3 | Lung Adenocarcinoma | (13) |
| c.2756C>T | p.Ser919Leu | S919L | 3 | Lung Adenocarcinoma | (11) |
| c.2762C>T | p.Ser921Phe | S921F | 4 | Renal Non-Clear Cell Carcinoma | (92) |
| c.2762C>T | p.Ser921Phe | S921F | 4 | Lung Adenocarcinoma | (11) |
| c.2762C>A | p.Ser921Tyr | S921Y | 4 | Breast Invasive Carcinoma, NOS | (13) |
| c.2762C>A | p.Ser921Tyr | S921Y | 4 | Uterine Endometrioid Carcinoma | (18-23,25) |
| c.2764C>T | p.Pro922Ser | P922S | 4 | Bladder Urothelial Carcinoma | (71) |
| c.2764C>T | p.Pro922Ser | P922S | 4 | Bladder Urothelial Carcinoma | (71) |
| c.2764C>T | p.Pro922Ser | P922S | 4 | Bladder Urothelial Carcinoma | (71) |
| c.2764C>T | p.Pro922Ser | P922S | 4 | Bladder Urothelial Carcinoma | (71) |
| c.2773A>G | p.Ile925Val | I925V | 1 | Bladder Urothelial Carcinoma | (40) |
| c.2782A>G | p.Arg928Gly | R928G | 3 | Lung Squamous Cell Carcinoma | (16,17) |
| c.2782A>G | p.Arg928Gly | R928G | 3 | Lung Squamous Cell Carcinoma | (16,17) |
| c.2783G>T | p.Arg928Ile | R928I | 3 | Rectal Adenocarcinoma | (18-23,25) |
| c.2785A>C | p.Thr929Pro | T929P | 1 | Cholangiocarcinoma | (26) |
| c.2788C>A | p.Leu930Met | L930M | 1 | Colon Adenocarcinoma | (11) |
| c.2792T>A | p.Ile931Lys | I931K | 2 | Colon Adenocarcinoma | (11) |
| c.2791A>G | p.Ile931Val | I931V | 2 | Hepatocellular Adenoma | (48) |
| c.2797G>C | p.Val933Leu | V933L | 1 | Uterine Endometrioid Carcinoma | (13) |
| c.2801C>T | p.Ala934Val | A934V | 1 | Skin Adnexal Carcinoma | (13) |
| c.2805A>C | p.Lys935Asn | K935N | 4 | Colon Adenocarcinoma | (13) |
| c.2805A>C | p.Lys935Asn | K935N | 4 | Colon Adenocarcinoma | (11) |
| c.2804A>C | p.Lys935Thr | K935T | 4 | Mucinous Adenocarcinoma of the Colon and Rectum | (18-23,25) |
| c.2804A>C | p.Lys935Thr | K935T | 4 | Mucinous Adenocarcinoma of the Colon and Rectum | (33) |
| c.2807C>T | p.Ser936Phe | S936F | 1 | Cutaneous Squamous Cell Carcinoma | (91) |
| c.2812C>A | p.Gln938Lys | Q938K | 1 | Hepatocellular Adenoma | (48) |
| c.2821G>T | p.Ala941Ser | A941S | 1 | Uterine Endometrioid Carcinoma | (18-23,25) |
| c.2824A>C | p.Asn942His | N942H | 3 | Colon Adenocarcinoma | (18-23,25) |
| c.2824A>C | p.Asn942His | N942H | 3 | Colon Adenocarcinoma | (63) |
| c.2824A>C | p.Asn942His | N942H | 3 | Uterine Endometrioid Carcinoma | (18-23,25) |
| c.2827C>T | p.Leu943Phe | L943F | 1 | Lung Adenocarcinoma | (11) |
| c.2833G>A | p.Glu945Lys | E945K | 1 | Colon Adenocarcinoma | (28) |
| c.2839G>A | p.Gly947Arg | G947R | 1 | Hepatocellular Carcinoma | (86) |
| c.2845A>G | p.Lys949Glu | K949E | 2 | Small Cell Lung Cancer | (70) |
| c.2846A>C | p.Lys949Thr | K949T | 2 | Lung Squamous Cell Carcinoma | (11) |
| c.2848G>A | p.Glu950Lys | E950K | 1 | Lung Squamous Cell Carcinoma | (11) |
| c.2857A>G | p.Met953Val | M953V | 1 | Lung Adenocarcinoma | (11) |
| c.2860G>A | p.Glu954Lys | E954K | 1 | Lung Squamous Cell Carcinoma | (16,17) |
| c.2864G>A | p.Gly955Asp | G955D | 1 | Uterine Endometrioid Carcinoma | (18-23,25) |
| c.2872C>G | p.Pro958Ala | P958A | 1 | Colorectal Adenocarcinoma | (14) |
| c.2877C>A | p.Phe959Leu | F959L | 2 | Head and Neck Squamous Cell Carcinoma | (93) |
| c.2877C>A | p.Phe959Leu | F959L | 2 | B-Lymphoblastic Leukemia/Lymphoma, BCR-ABL1 Like | (94) |
| c.2885G>T | p.Ser962Ile | S962I | 1 | Prostate Adenocarcinoma | (42) |
| c.2896C>T | p.Arg966Cys | R966C | 8 | Stomach Adenocarcinoma | (95) |
| c.2896C>T | p.Arg966Cys | R966C | 8 | Breast Invasive Ductal Carcinoma | (11) |
| c.2896C>T | p.Arg966Cys | R966C | 8 | Colon Adenocarcinoma | (11) |
| c.2896C>T | p.Arg966Cys | R966C | 8 | Colorectal Adenocarcinoma | (14) |
| c.2897G>A | p.Arg966His | R966H | 8 | Invasive Breast Carcinoma | (35) |
| c.2897G>A | p.Arg966His | R966H | 8 | Colon Adenocarcinoma | (11) |

| | | | | | |
|---|---|---|---|---|---|
| c.2897G>A | p.Arg966His | R966H | 8 | Uterine Endometrioid Carcinoma | (18-23,25) |
| c.2897G>A | p.Arg966His | R966H | 8 | Uterine Endometrioid Carcinoma | (18-23,25) |
| c.2901G>T | p.Met967Ile | M967I | 2 | Colorectal Adenocarcinoma | (44) |
| c.2900T>C | p.Met967Thr | M967T | 2 | Colon Adenocarcinoma | (11) |
| c.2904C>G | p.Ile968Met | I968M | 1 | Prostate Adenocarcinoma | (13) |
| c.2921T>C | p.Leu974Pro | L974P | 2 | Invasive Breast Carcinoma | (11) |
| c.2921T>G | p.Leu974Arg | L974R | 2 | Uterine Endometrioid Carcinoma | (18-23,25) |
| c.2924G>T | p.Gly975Val | G975V | 1 | Cutaneous Melanoma | (18-23) |
| c.2930T>C | p.Val977Ala | V977A | 1 | Colon Adenocarcinoma | (11) |
| c.2948C>A | p.Thr983Asn | T983N | 1 | Uterine Endometrioid Carcinoma | (18-23,25) |
| c.2960C>A | p.Ser987Tyr | S987Y | 1 | Colorectal Adenocarcinoma | (95) |
| c.2963G>T | p.Arg988Ile | R988I | 1 | Colon Adenocarcinoma | (11) |
| c.2966C>T | p.Thr989Met | T989M | 2 | Breast Invasive Lobular Carcinoma | (13) |
| c.2966C>T | p.Thr989Met | T989M | 2 | Melanoma | (33) |
| c.2971C>A | p.Leu991Met | L991M | 1 | Colon Adenocarcinoma | (11) |
| c.2975C>T | p.Ser992Phe | S992F | 2 | Bladder Urothelial Carcinoma | (13) |
| c.2974T>C | p.Ser992Pro | S992P | 2 | Mucinous Adenocarcinoma of the Colon and Rectum | (13) |
| c.2977C>T | p.Arg993Cys | R993C | 4 | Colorectal Adenocarcinoma | (11) |
| c.2977C>T | p.Arg993Cys | R993C | 4 | Intrahepatic Cholangiocarcinoma | (68) |
| c.2977C>G | p.Arg993Gly | R993G | 4 | Upper Tract Urothelial Carcinoma | (13) |
| c.2978G>A | p.Arg993His | R993H | 4 | Stomach Adenocarcinoma | (13) |
| c.2980G>A | p.Asp994Asn | D994N | 1 | Hepatocellular Carcinoma | (24) |
| c.2998G>A | p.Glu1000Lys | E1000K | 1 | Lung Squamous Cell Carcinoma | (11) |
| c.3001A>T | p.Ile1001Phe | I1001F | 1 | Colon Adenocarcinoma | (11) |
| c.3007G>A | p.Val1003Met | V1003M | 2 | Uterine Serous Carcinoma/Uterine Papillary Serous Carcinoma | (18-23,25) |
| c.3007G>A | p.Val1003Met | V1003M | 2 | Melanoma | (69) |
| c.3010G>T | p.Ala1004Ser | A1004S | 3 | Lung Adenocarcinoma | (11) |
| c.3010G>T | p.Ala1004Ser | A1004S | 3 | Myxofibrosarcoma | (18-23,25) |
| c.3011C>T | p.Ala1004Val | A1004V | 3 | Colon Adenocarcinoma | (18-23,25) |
| c.3014A>G | p.His1005Arg | H1005R | 4 | Glioblastoma Multiforme | (96) |
| c.3014A>G | p.His1005Arg | H1005R | 4 | Glioblastoma Multiforme | (13) |
| c.3013C>T | p.His1005Tyr | H1005Y | 4 | Invasive Breast Carcinoma | (11) |
| c.3013C>T | p.His1005Tyr | H1005Y | 4 | Lung Neuroendocrine Tumor | (11) |
| c.3017C>T | p.Ser1006Leu | S1006L | 1 | Uterine Endometrioid Carcinoma | (18-23,25) |
| c.3023A>G | p.Glu1008Gly | E1008G | 1 | Lung Adenocarcinoma | (11) |
| c.3026T>G | p.Leu1009Arg | L1009R | 2 | Adenoid Cystic Carcinoma | (67) |
| c.3026T>G | p.Leu1009Arg | L1009R | 2 | Adenoid Cystic Carcinoma | (13) |
| c.3029G>A | p.Arg1010Gln | R1010Q | 7 | Colon Adenocarcinoma | (63) |
| c.3029G>A | p.Arg1010Gln | R1010Q | 7 | Upper Tract Urothelial Carcinoma | (13) |
| c.3029G>A | p.Arg1010Gln | R1010Q | 7 | Uterine Endometrioid Carcinoma | (13) |
| c.3029G>A | p.Arg1010Gln | R1010Q | 7 | Lung Adenocarcinoma | (11) |
| c.3029G>A | p.Arg1010Gln | R1010Q | 7 | Colon Adenocarcinoma | (11) |
| c.3029G>A | p.Arg1010Gln | R1010Q | 7 | Colon Adenocarcinoma | (11) |
| c.3029G>A | p.Arg1010Gln | R1010Q | 7 | Colorectal Adenocarcinoma | (14) |
| c.3032C>T | p.Thr1011Met | T1011M | 1 | Rectal Adenocarcinoma | (11) |
| c.3037A>G | p.Ser1013Gly | S1013G | 5 | Intrahepatic Cholangiocarcinoma | (13) |
| c.3037A>G | p.Ser1013Gly | S1013G | 5 | Hepatocellular Carcinoma | (33) |
| c.3037A>G | p.Ser1013Gly | S1013G | 5 | Hepatocellular Carcinoma | (18-23,25) |
| c.3038G>A | p.Ser1013Asn | S1013N | 5 | Skin | (58) |
| c.3038G>C | p.Ser1013Thr | S1013T | 5 | Endometrial Carcinoma | (30) |
| c.3046C>T | p.Arg1016Cys | R1016C | 9 | Prostate Adenocarcinoma | (97) |
| c.3046C>T | p.Arg1016Cys | R1016C | 9 | Cutaneous Squamous Cell Carcinoma | (13) |
| c.3046C>T | p.Arg1016Cys | R1016C | 9 | Colon Adenocarcinoma | (11) |
| c.3046C>T | p.Arg1016Cys | R1016C | 9 | Medullary Carcinoma of the Colon | (11) |

| | | | | | |
|---|---|---|---|---|---|
| c.3046C>T | p.Arg1016Cys | R1016C | 9 | Mucinous Adenocarcinoma of the Colon and Rectum | (11) |
| c.3047G>A | p.Arg1016His | R1016H | 9 | Uterine Endometrioid Carcinoma | (13) |
| c.3047G>A | p.Arg1016His | R1016H | 9 | Rectal Adenocarcinoma | (11) |
| c.3047G>A | p.Arg1016His | R1016H | 9 | Head and Neck Squamous Cell Carcinoma | (98) |
| c.3047G>A | p.Arg1016His | R1016H | 9 | Uterine Endometrioid Carcinoma | (18-23,25) |
| c.3050G>T | p.Gly1017Val | G1017V | 2 | Breast Invasive Lobular Carcinoma | (13) |
| c.3050G>T | p.Gly1017Val | G1017V | 2 | Breast Invasive Lobular Carcinoma | (11) |
| c.3053C>T | p.Ala1018Val | A1018V | 1 | Rectal Adenocarcinoma | (11) |
| c.3056A>T | p.Gln1019Leu | Q1019L | 1 | Lung Adenocarcinoma | (11) |
| c.3060G>C | p.Gln1020His | Q1020H | 4 | Colon Adenocarcinoma | (13) |
| c.3060G>C | p.Gln1020His | Q1020H | 4 | Colon Adenocarcinoma | (11) |
| c.3060G>C | p.Gln1020His | Q1020H | 4 | Lung Squamous Cell Carcinoma | (18-23,25) |
| c.3058C>A | p.Gln1020Lys | Q1020K | 4 | Colorectal Adenocarcinoma | (14) |
| c.3063C>G | p.His1021Gln | H1021Q | 2 | Lung Adenocarcinoma | (11) |
| c.3061C>T | p.His1021Tyr | H1021Y | 2 | Cancer of Unknown Primary | (13) |
| c.3064G>A | p.Val1022Ile | V1022I | 2 | Colon Adenocarcinoma | (63) |
| c.3064G>A | p.Val1022Ile | V1022I | 2 | Uterine Endometrioid Carcinoma | (18-23,25) |
| c.3082G>A | p.Ala1028Thr | A1028T | 1 | Endometrial Carcinoma | (47) |
| c.3085A>G | p.Ile1029Val | I1029V | 2 | Anaplastic Oligoastrocytoma | (13) |
| c.3085A>G | p.Ile1029Val | I1029V | 2 | Head and Neck Squamous Cell Carcinoma | (18-23) |
| c.3091G>A | p.Glu1031Lys | E1031K | 1 | Colon Adenocarcinoma | (11) |
| c.3095T>C | p.Leu1032Pro | L1032P | 1 | Melanoma | (99) |
| c.3100C>G | p.Gln1034Glu | Q1034E | 3 | Renal Clear Cell Carcinoma | (13) |
| c.3101A>C | p.Gln1034Pro | Q1034P | 3 | Colon Adenocarcinoma | (13) |
| c.3101A>C | p.Gln1034Pro | Q1034P | 3 | Colon Adenocarcinoma | (11) |
| c.3106A>C | p.Lys1036Gln | K1036Q | 1 | Colon Adenocarcinoma | (11) |
| c.3109C>G | p.Gln1037Glu | Q1037E | 3 | Breast Invasive Ductal Carcinoma | (13) |
| c.3109C>G | p.Gln1037Glu | Q1037E | 3 | Breast Invasive Ductal Carcinoma | (11) |
| c.3111A>C | p.Gln1037His | Q1037H | 3 | Tubular Stomach Adenocarcinoma | (18-23,25) |
| c.3113A>G | p.Asn1038Ser | N1038S | 1 | Lung Adenocarcinoma | (18-23) |
| c.3117G>C | p.Gln1039His | Q1039H | 1 | Mucinous Adenocarcinoma of the Colon and Rectum | (11) |
| c.3137T>C | p.Val1046Ala | V1046A | 1 | Colon Adenocarcinoma | (63) |
| c.3140G>A | p.Arg1047Lys | R1047K | 1 | Uterine Endometrioid Carcinoma | (18-23,25) |

**Supplementary Table 3. Mutations observed in both vascular malformations and cancer.**

| Residue # | Number of Vascular Mutations | Vascular Mutation | Number of Cancer Mutations | Cancer Mutation(s) |
|---|---|---|---|---|
| 70 | 1 | E70Q | 3 | E70D/G/Q |
| 99 | 4 | A99V | 2 | A99V |
| 156 | 1 | G156V | 1 | G156C |
| 426 | 8 | Y426C | 2 | Y426C/H |
| 451 | 1 | N451S | 1 | N451S |
| 528 | 4 | Y528C | 1 | Y528H |
| 530 | 6 | V530D | 1 | V530I |
| 707 | 4 | R707C | 9 | R707C/H/L/P |
| 763 | 3 | E763V | 8 | E763G/K |

**Supplementary Table 4. Thermal shift assay values.** $\Delta T_m$ indicates difference compared to wild type (WT) for that construct. SD indicates standard deviation. N=3.

| RasGAP Construct | $T_m$ Mean (°C) | $T_m$ SD (°C) | $\Delta T_m$ Mean (°C) | $\Delta T_m$ SD (°C) |
|---|---:|---:|---:|---:|
| WT | 44.35 | 0.68 | - | 0.68 |
| R194C | 42.21 | 0.65 | -2.14 | 0.65 |
| R283H | 39.62 | 0.56 | -4.73 | 0.56 |
| R427Q | 44.01 | 0.79 | -0.34 | 0.79 |
| Y528C | 40.92 | 0.45 | -3.43 | 0.45 |
| R591C | 43.55 | 0.93 | -0.80 | 0.93 |
| H604N | 44.02 | 0.43 | -0.33 | 0.43 |
| T612A | 44.26 | 0.69 | -0.09 | 0.69 |
| W689R | 42.99 | 0.28 | -1.36 | 0.28 |
| H695L | 44.81 | 0.37 | 0.46 | 0.37 |
| S705F | 44.61 | 0.35 | 0.26 | 0.35 |
| R707H | 41.15 | 0.31 | -3.20 | 0.31 |

**Supplementary Table 5. Isothermal titration calorimetry analyses for RasGAP interactions with phosphorylated p190RhoGAP.** Synthesized p190RhoGAP peptide (residues 1083-1111, phosphorylated on pY1087 and pY1105) titrated against RasGAP SH2-SH3-SH2 construct (residues 174-444). Parameters from individual replicates are shown with their corresponding panel in **Supplementary Figure 6**. Averages are shown as mean ± standard deviation (SD).

| Construct | Figure S6 Panel | Protein Conc-entration in Cell (μM) | Peptide Conc-entration in Syringe (μM) | $K_D$ (nM) | n | ΔH (kJ/mol) | ΔS (J/mol•K) | Blank Constant (μJ) | $K_A$ (nM$^{-1}$) | TΔS (kJ/mol) | ΔG (kJ/mol) |
|---|---|---|---|---|---|---|---|---|---|---|---|
| WT | A | 4.94 | 36.3 | 6.49 | 1.356 | -63.72 | -59.96 | -2.2 | 0.1542 | -16.98 | -46.74 |
| WT | B | 4.94 | 36.3 | 3.59 | 1.453 | -61.36 | -44.13 | -2.9 | 0.2789 | -13.16 | -48.21 |
| WT Average | – | – | – | 5.04 ± 2.05 | 1.405 ± 0.069 | -62.54 ± 1.67 | -52.05 ± 11.19 | -2.55 ± 0.49 | 0.2166 ± 0.0882 | -15.07 ± 2.70 | -47.48 ± 1.04 |
| R194C | C | 4.39 | 24.8 | 3.65 | 1.038 | -118.3 | -235.2 | -3 | 0.2470 | -70.13 | -48.16 |
| R194C | D | 4.41 | 24.8 | 7.23 | 0.981 | -86.06 | -132.8 | -5 | 0.1383 | -39.60 | -46.47 |
| R194C | E | 4.41 | 24.8 | 1.20 | 0.897 | -96.4 | -152.6 | -5 | 0.8329 | -45.48 | -50.92 |
| R194C Average | – | – | – | 4.03 ± 3.03 | 0.972 ± 0.071 | -100.3 ± 16.5 | -173.5 ± 54.3 | -4.33 ± 1.15 | 0.4061 ± 0.3736 | -51.74 ± 16.20 | -48.26 ± 2.25 |
| R427Q | F | 4.24 | 24.8 | 8.14 | 0.967 | -104.8 | -196.6 | -3 | 0.1229 | -58.62 | -46.17 |
| R427Q | G | 4.24 | 24.8 | 2.08 | 0.932 | -102.9 | -178.8 | -3.5 | 0.4808 | -53.30 | -49.56 |
| R427Q Average | – | – | – | 5.11 ± 4.29 | 0.950 ± 0.025 | -103.9 ± 1.3 | -187.7 ± 12.6 | -3.25 ± 0.35 | 0.3019 ± 0.2531 | -55.96 ± 3.76 | -47.87 ± 2.40 |

**Supplementary Table 6. Michaelis-Menten kinetics assessment of RasGAP and disease associated RasGAP mutants.** Mean and standard deviation (SD) indicated for each sample. N=5.

| **RasGAP Construct** | **$k_{cat}$ Mean ($s^{-1}$)** | **$k_{cat}$ SD ($s^{-1}$)** | **$K_M$ Mean (μM)** | **$K_M$ SD (μM)** | **$k_{cat}/K_M$ Mean ($M^{-1}s^{-1}$)** | **$k_{cat}/K_M$ SD ($M^{-1}s^{-1}$)** |
|---|---|---|---|---|---|---|
| WT | 19.0 | 5.0 | 46.2 | 10.3 | 425,520 | 121,760 |
| R194C | 16.9 | 6.5 | 186.3 | 128.7 | 136,734 | 108,155 |
| R283H | 5.0 | 0.9 | 99.1 | 40.1 | 57,470 | 23,165 |
| R427Q | 13.9 | 3.1 | 49.6 | 17.1 | 291,720 | 73,738 |
| Y528C | 12.2 | 1.8 | 51.7 | 6.8 | 235,880 | 28,978 |
| R591C | 9.4 | 2.6 | 116.2 | 26.7 | 82,816 | 19,636 |
| H604N | 11.8 | 3.1 | 143.4 | 28.0 | 81,800 | 9,447 |
| T612A | 19.0 | 4.7 | 58.5 | 9.7 | 334,440 | 108,161 |
| W689R | 15.5 | 5.4 | 102.1 | 52.3 | 169,860 | 49,463 |
| H695L | 27.9 | 11.4 | 65.7 | 29.7 | 429,200 | 45,196 |
| S705F | 15.1 | 3.6 | 125.9 | 60.3 | 130,698 | 32,769 |
| R707H | 6.7 | 4.7 | 731.1 | 441.5 | 8,952 | 746 |

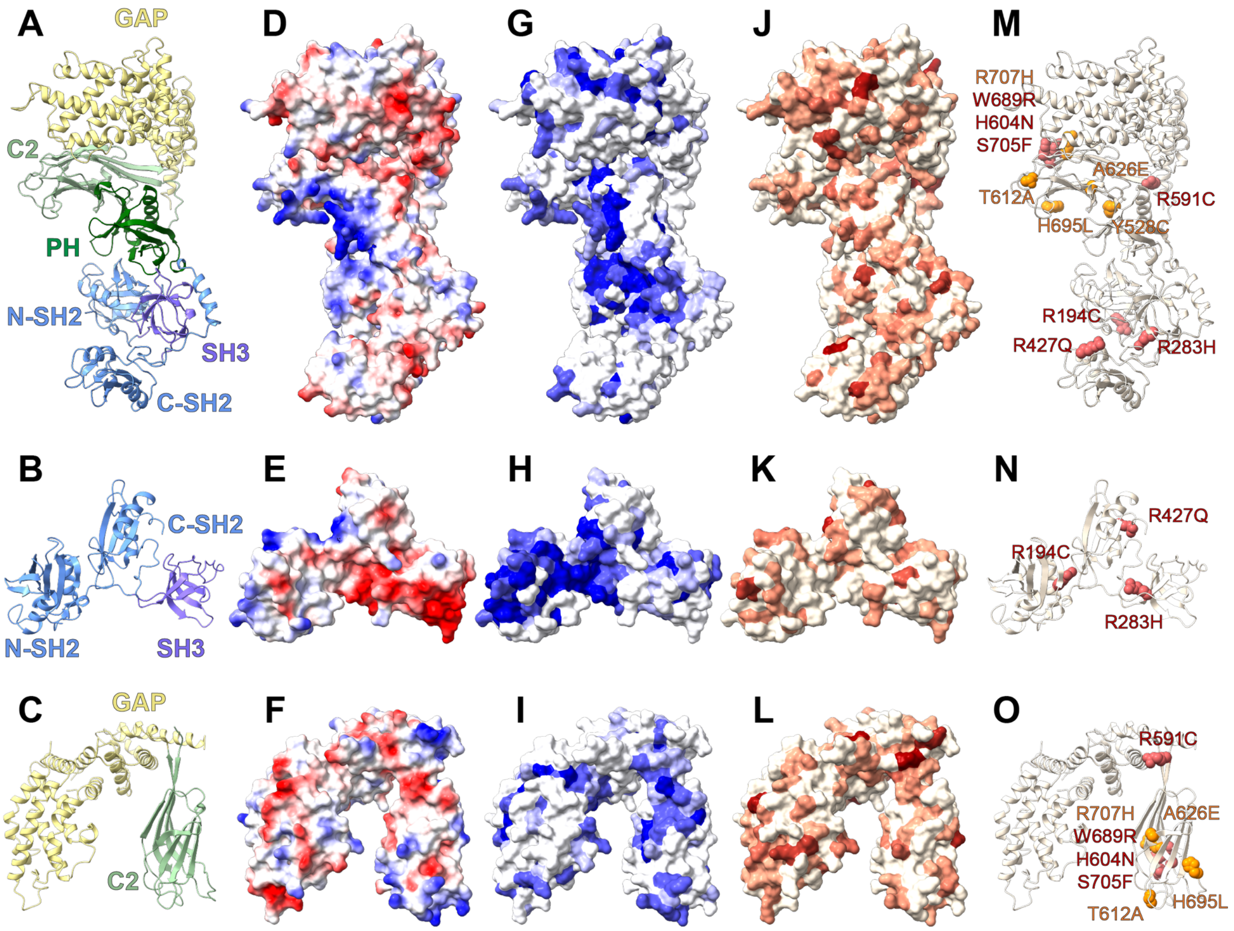


**Supplementary Figure 1. Reverse views mapping *RASA1* mutations onto RasGAP.** The figure depicts the AlphaFold prediction of ordered domains of RasGAP (**A**, **D**, **G**, **J**, **M**, top row) (AlphaFold ID: AF-P20936-F1-v4), crystal structure of the SH2-SH3-SH2 region (**B**, **E**, **H**, **K**, **N**, middle row) (PDB accession number: 8DGQ (100)), and crystal structure of the C2-GAP region (**C**, **F**, **I**, **L**, **O**, bottom row) (PDB accession number: 9BZ4 (101)). **A-C)** Ribbon diagrams colored by domain. **D-F)** Electrostatics of the protein surface. Positive (blue) and negative (red) charge indicated. **G-I)** Sequence conservation from low (white) to high (blue) mapped onto the protein surface. **J-L)** *RASA1* mutations mapped onto the protein surface. No reported mutations colored white, most frequent colored dark red. **M-O)** Mutations chosen for further study shown as spheres. Red indicates cancer-associated, orange indicates vascular malformations associated.

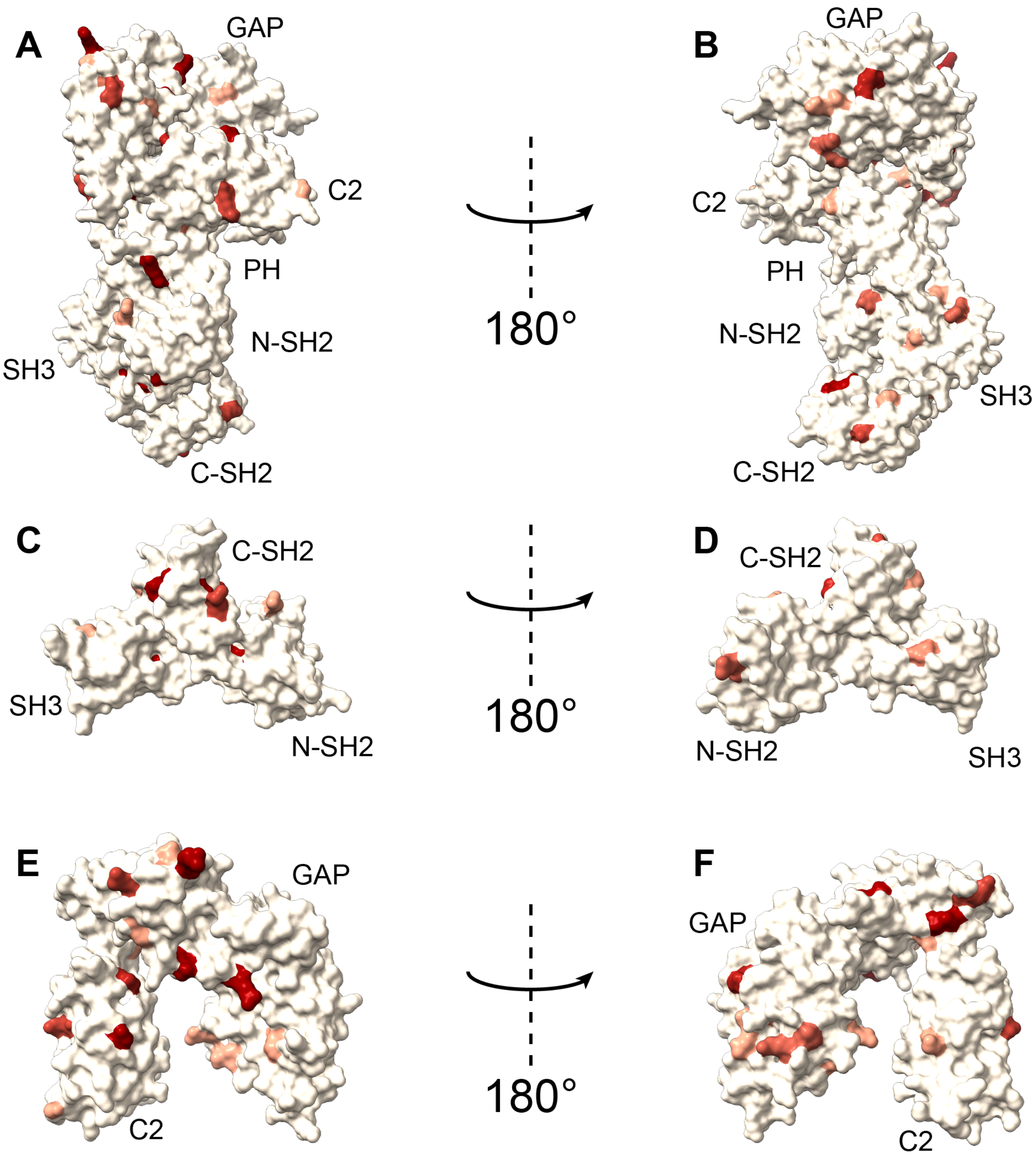


**Supplementary Figure 2. Mapping hotspots of *RASA1* mutations onto RasGAP.** Mutations observed more than four times depicted in red on the AlphaFold prediction of ordered domains of RasGAP (**A**, **B**, top row) (AlphaFold ID: AF-P20936-F1-v4), the crystal structure of the SH2-SH3-SH2 region (**C**, **D**, middle row) (PDB accession number: 8DGQ (100)), and the crystal structure of the C2-GAP region (**E**, **F**, bottom row) (PDB accession number: 9BZ4 (101)).

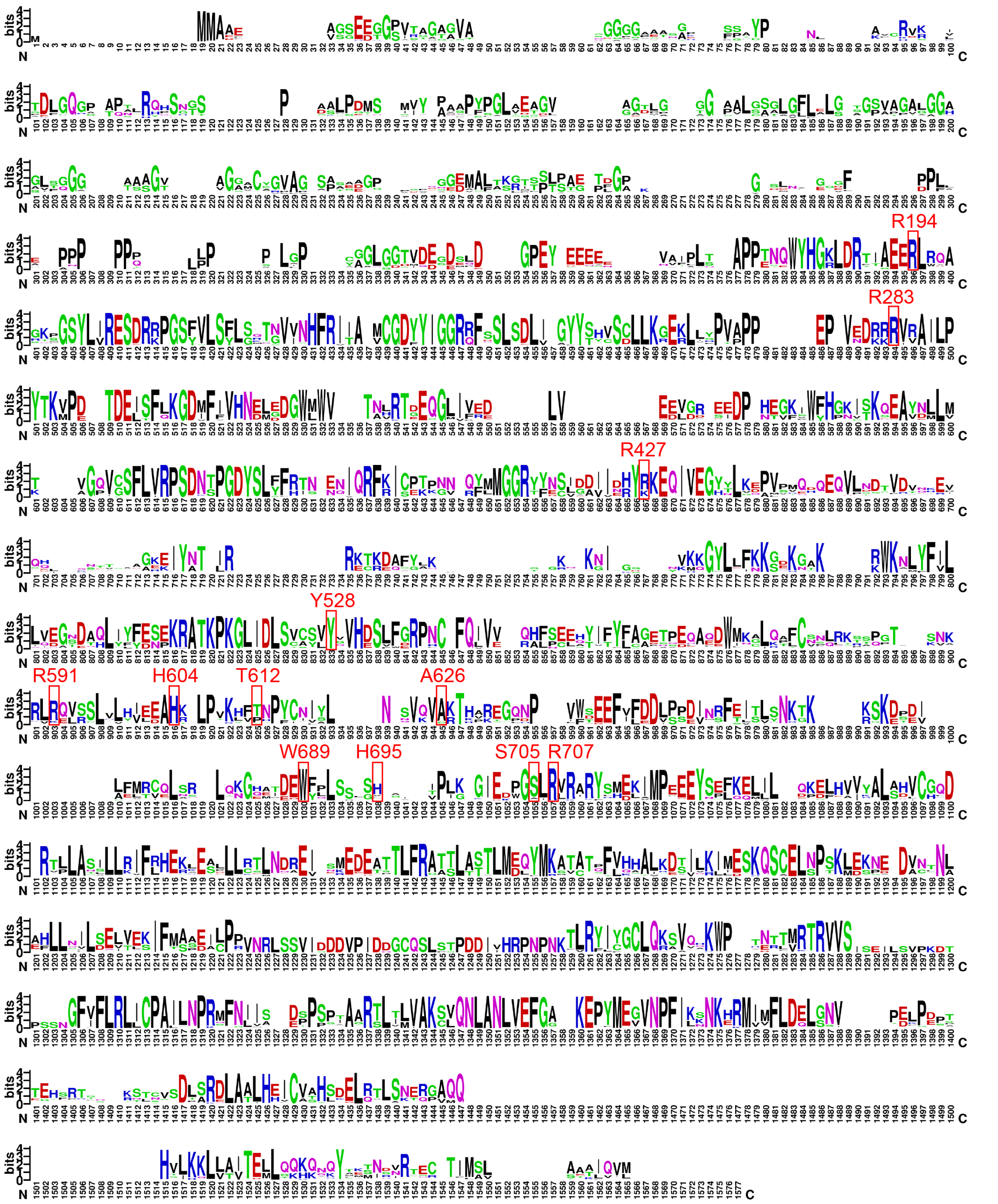


**Supplementary Figure 3. Complete sequence alignment/conservation of p120RasGAP.** Sequence logo based on an alignment of 209 RasGAP homologue sequences from sponges to humans. Numbering of residues is based on the overall alignment and does not correspond directly to the human p120RasGAP sequence. Height on y-axis indicates the relative frequency of each amino acid at that position. Basic residues are blue, acidic residues are red, hydrophobic residues are black, and polar residues are green or purple. Locations of mutants studied are indicated. Figure created using WebLogo (102).

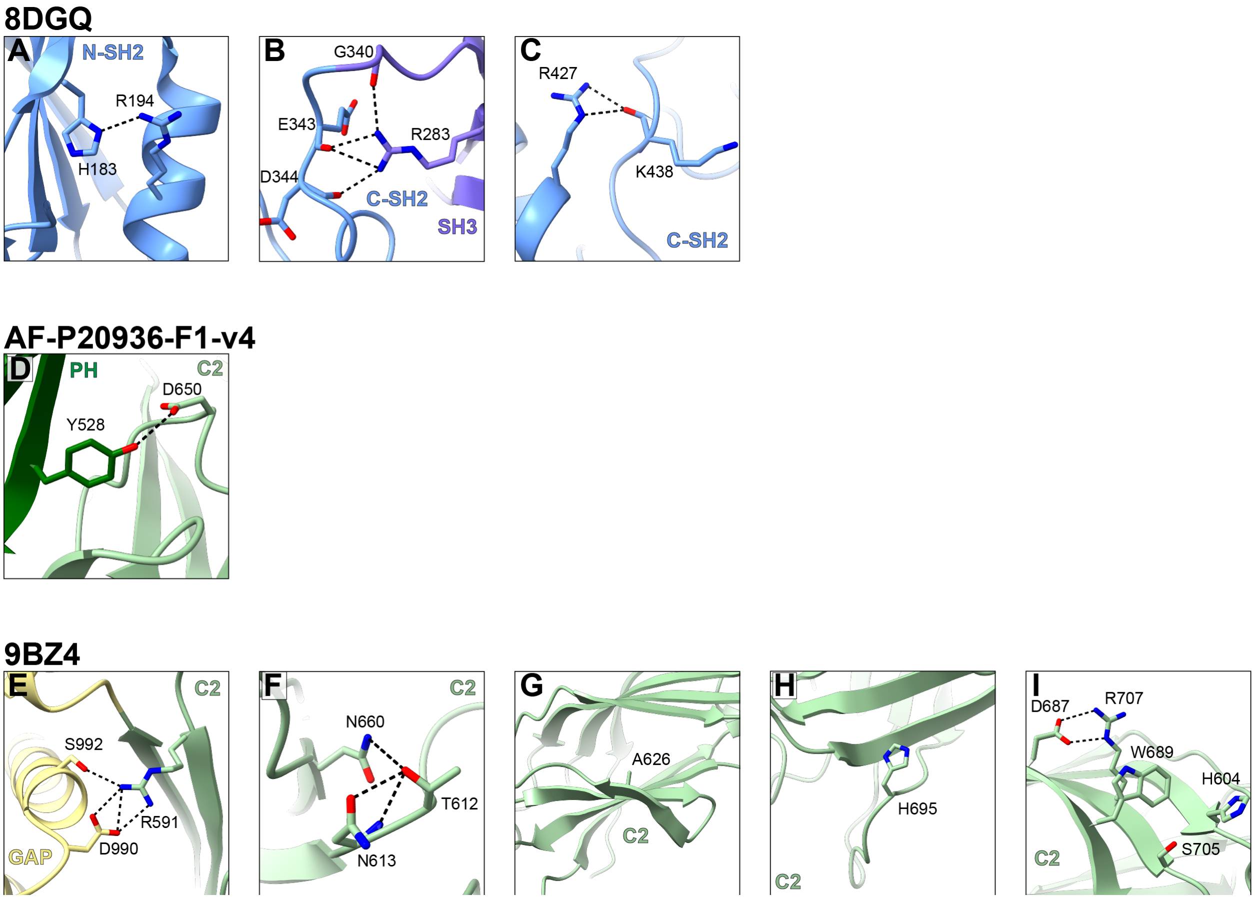


**Supplementary Figure 4. Hydrogen bonding of RasGAP mutants.** Hydrogen bonding of the twelve disease-associated missense mutants studied are shown as black dashed lines and engaged neighboring residues indicated. **A-C)** Crystal structure (8DGQ (100)) shown for residues R194, R283 and R427. **D)** AlphaFold structure prediction shown for residue Y528. **E-I)** Crystal structure (9BZ4 (101)) shown for residues R591, T612, A626, H695, R707, H604, W689, S705.

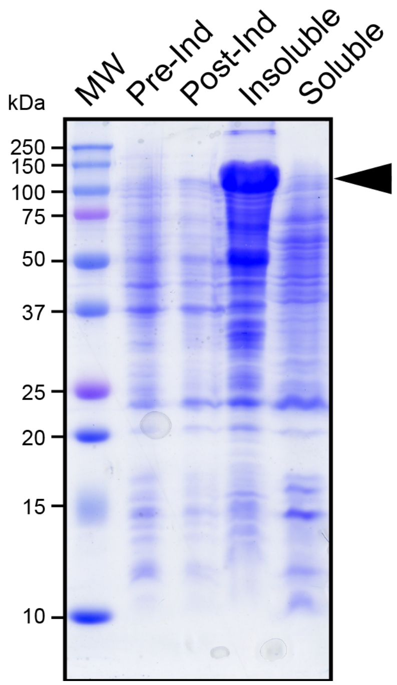


**Supplementary Figure 5. Expression and solubility of RasGAP A626E mutant.** Expression of p120RasGAP ∆N A626E shown by SDS-PAGE gel (Coomassie). Lane labels: MW, molecular weight; Pre-Ind, pre-induction; Post-Ind, post-induction; Insoluble, pelleted cell extract; Soluble, soluble cell extract. Arrow indicates expected molecular weight (103.5 kDa).

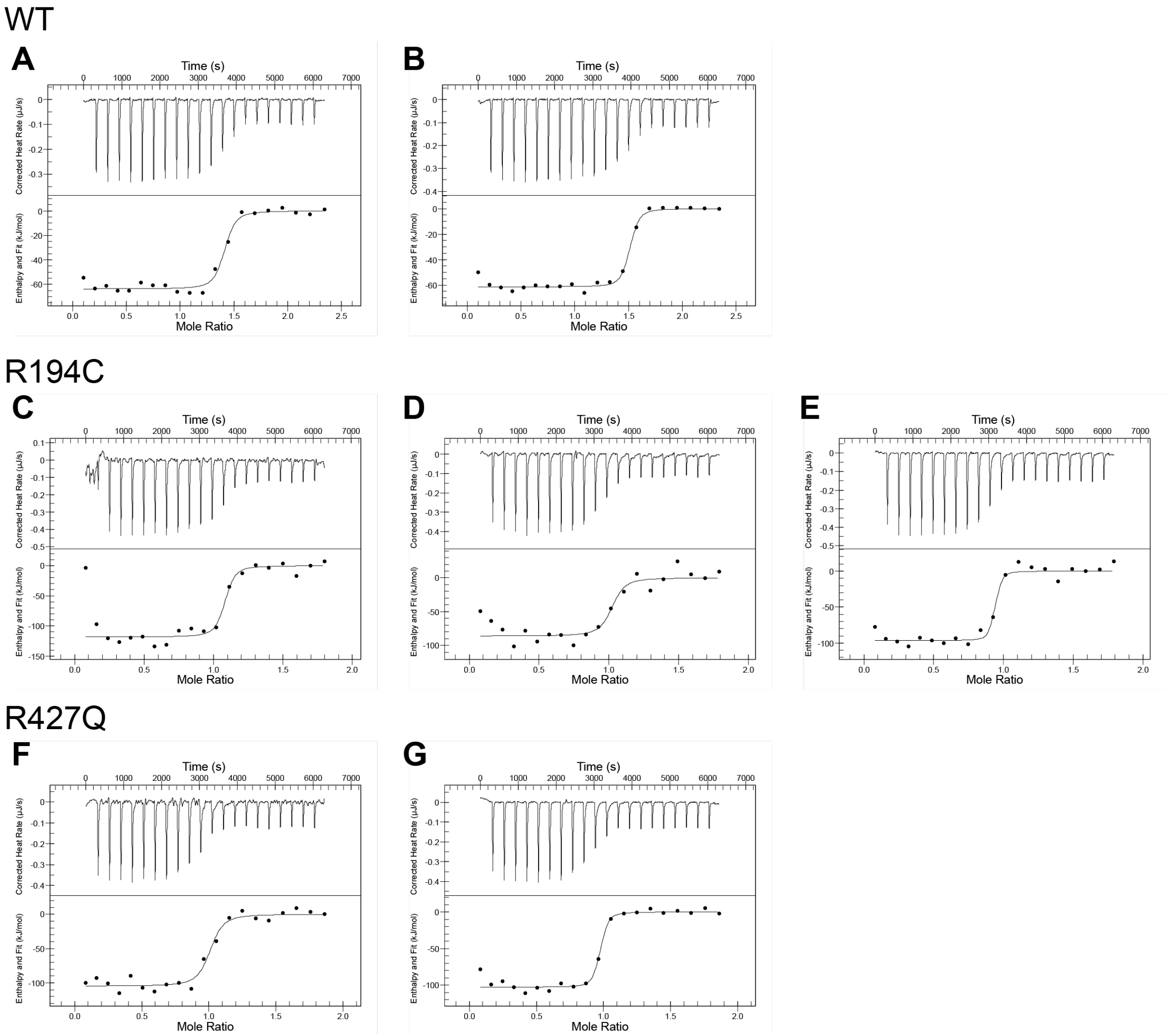


**Supplementary Figure 6. Isothermal titration calorimetry traces and isotherms for RasGAP interactions with phosphorylated p190RhoGAP.** Isothermal titration calorimetry traces (top) and isotherms (bottom) for individual replicates. Parameters obtained from the curve fit for each replicate and averages for each construct are presented in **Supplementary Table 5**.

10. Rizvi, H., Sanchez-Vega, F., La, K., Chatila, W., Jonsson, P., Halpenny, D., Plodkowski, A., Long, N., Sauter, J. L., Rekhtman, N., Hollmann, T., Schalper, K. A., Gainor, J. F., Shen, R., Ni, A., Arbour, K. C., Merghoub, T., Wolchok, J., Snyder, A., Chaft, J. E., Kris, M. G., Rudin, C. M., Socci, N. D., Berger, M. F., Taylor, B. S., Zehir, A., Solit, D. B., Arcila, M. E., Ladanyi, M., Riely, G. J., Schultz, N., and Hellmann, M. D. (2018) Molecular Determinants of Response to Anti-Programmed Cell Death (PD)-1 and Anti-Programmed Death-Ligand 1 (PD-L1) Blockade in Patients With Non-Small-Cell Lung Cancer Profiled With Targeted Next-Generation Sequencing. *J Clin Oncol* **36**, 633-641
11. Jee, J., Fong, C., Pichotta, K., Tran, T. N., Luthra, A., Waters, M., Fu, C., Altoe, M., Liu, S. Y., Maron, S. B., Ahmed, M., Kim, S., Pirun, M., Chatila, W. K., de Bruijn, I., Pasha, A., Kundra, R., Gross, B., Mastrogiacomo, B., Aprati, T. J., Liu, D., Gao, J., Capelletti, M., Pekala, K., Loudon, L., Perry, M., Bandlamudi, C., Donoghue, M., Satravada, B. A., Martin, A., Shen, R., Chen, Y., Brannon, A. R., Chang, J., Braunstein, L., Li, A., Safonov, A., Stonestrom, A., Sanchez-Vela, P., Wilhelm, C., Robson, M., Scher, H., Ladanyi, M., Reis-Filho, J. S., Solit, D. B., Jones, D. R., Gomez, D., Yu, H., Chakravarty, D., Yaeger, R., Abida, W., Park, W., O'Reilly, E. M., Garcia-Aguilar, J., Socci, N., Sanchez-Vega, F., Carrot-Zhang, J., Stetson, P. D., Levine, R., Rudin, C. M., Berger, M. F., Shah, S. P., Schrag, D., Razavi, P., Kehl, K. L., Li, B. T., Riely, G. J., Schultz, N., and Group, M. S. K. C. D. S. I. (2024) Automated real-world data integration improves cancer outcome prediction. *Nature* **636**, 728-736
12. Jain, E., Zañudo, J. G. T., McGillicuddy, M., Abravanel, D. L., Thomas, B. S., Kim, D., Balch, S., Navarro, J., Weiss, J. H., Hernandez, T. G., Dunphy, M., Tomson, B. N., Buendia-Buendia, J., Alao, O., Damon, A. L., Di Lascio, S., Shah, S., Small, I. K., Sosa, D., Sterlin, L., Boykin, I., Stoddard, R. E., Tsegai, N., Ulysse, U. F., Phelps, K., Frank, E., Kumari, P., Maiwald, S., Larkin, K., Pollock, S., Zimmer, A., Chastain, P. S., Cusher, T., Nguyen, C., Winnicki, S., Anastasio, E., Van Allen, E. M., Lander, E. S., Golub, T. R., Painter, C. A., and Wagle, N. (2023) The Metastatic Breast Cancer Project: leveraging patient-partnered research to expand the clinical and genomic landscape of metastatic breast cancer and accelerate discoveries. *medRxiv*, 2023.2006.2007.23291117
13. Zehir, A., Benayed, R., Shah, R. H., Syed, A., Middha, S., Kim, H. R., Srinivasan, P., Gao, J., Chakravarty, D., Devlin, S. M., Hellmann, M. D., Barron, D. A., Schram, A. M., Hameed, M., Dogan, S., Ross, D. S., Hechtman, J. F., DeLair, D. F., Yao, J., Mandelker, D. L., Cheng, D. T., Chandramohan, R., Mohanty, A. S., Ptashkin, R. N., Jayakumaran, G., Prasad, M., Syed, M. H., Rema, A. B., Liu, Z. Y., Nafa, K., Borsu, L., Sadowska, J., Casanova, J., Bacares, R., Kiecka, I. J., Razumova, A., Son, J. B., Stewart, L., Baldi, T., Mullaney, K. A., Al-Ahmadie, H., Vakiani, E., Abeshouse, A. A., Penson, A. V., Jonsson, P., Camacho, N., Chang, M. T., Won, H. H., Gross, B. E., Kundra, R., Heins, Z. J., Chen, H. W., Phillips, S., Zhang, H., Wang, J., Ochoa, A., Wills, J., Eubank, M., Thomas, S. B., Gardos, S. M., Reales, D. N., Galle, J., Durany, R., Cambria, R., Abida, W., Cercek, A., Feldman, D. R., Gounder, M. M., Hakimi, A. A., Harding, J. J., Iyer, G., Janjigian, Y. Y., Jordan, E. J., Kelly, C. M., Lowery, M. A., Morris, L. G. T., Omuro, A. M., Raj, N., Razavi, P., Shoushtari, A. N., Shukla, N., Soumerai, T. E., Varghese, A. M., Yaeger, R., Coleman, J., Bochner, B., Riely, G. J., Saltz, L. B., Scher, H. I., Sabbatini, P. J., Robson, M. E., Klimstra, D. S., Taylor, B. S., Baselga, J., Schultz, N., Hyman, D. M., Arcila, M. E., Solit, D. B., Ladanyi, M., and Berger, M. F. (2017) Mutational landscape of metastatic cancer revealed from prospective clinical sequencing of 10,000 patients. *Nat Med* **23**, 703-713
14. Giannakis, M., Mu, X. J., Shukla, S. A., Qian, Z. R., Cohen, O., Nishihara, R., Bahl, S., Cao, Y., Amin-Mansour, A., Yamauchi, M., Sukawa, Y., Stewart, C., Rosenberg, M., Mima, K., Inamura, K., Nosho, K., Nowak, J. A., Lawrence, M. S., Giovannucci, E. L., Chan, A. T., Ng, K., Meyerhardt, J. A., Van Allen, E. M., Getz, G., Gabriel, S. B., Lander,

E. S., Wu, C. J., Fuchs, C. S., Ogino, S., and Garraway, L. A. (2016) Genomic Correlates of Immune-Cell Infiltrates in Colorectal Carcinoma. *Cell Rep* **17**, 1206
15. Witkiewicz, A. K., McMillan, E. A., Balaji, U., Baek, G., Lin, W. C., Mansour, J., Mollaee, M., Wagner, K. U., Koduru, P., Yopp, A., Choti, M. A., Yeo, C. J., McCue, P., White, M. A., and Knudsen, E. S. (2015) Whole-exome sequencing of pancreatic cancer defines genetic diversity and therapeutic targets. *Nat Commun* **6**, 6744
16. Abbosh, C., Birkbak, N. J., Wilson, G. A., Jamal-Hanjani, M., Constantin, T., Salari, R., Le Quesne, J., Moore, D. A., Veeriah, S., Rosenthal, R., Marafioti, T., Kirkizlar, E., Watkins, T. B. K., McGranahan, N., Ward, S., Martinson, L., Riley, J., Fraioli, F., Al Bakir, M., Gronroos, E., Zambrana, F., Endozo, R., Bi, W. L., Fennessy, F. M., Sponer, N., Johnson, D., Laycock, J., Shafi, S., Czyzewska-Khan, J., Rowan, A., Chambers, T., Matthews, N., Turajlic, S., Hiley, C., Lee, S. M., Forster, M. D., Ahmad, T., Falzon, M., Borg, E., Lawrence, D., Hayward, M., Kolvekar, S., Panagiotopoulos, N., Janes, S. M., Thakrar, R., Ahmed, A., Blackhall, F., Summers, Y., Hafez, D., Naik, A., Ganguly, A., Kareht, S., Shah, R., Joseph, L., Marie Quinn, A., Crosbie, P. A., Naidu, B., Middleton, G., Langman, G., Trotter, S., Nicolson, M., Remmen, H., Kerr, K., Chetty, M., Gomersall, L., Fennell, D. A., Nakas, A., Rathinam, S., Anand, G., Khan, S., Russell, P., Ezhil, V., Ismail, B., Irvin-Sellers, M., Prakash, V., Lester, J. F., Kornaszewska, M., Attanoos, R., Adams, H., Davies, H., Oukrif, D., Akarca, A. U., Hartley, J. A., Lowe, H. L., Lock, S., Iles, N., Bell, H., Ngai, Y., Elgar, G., Szallasi, Z., Schwarz, R. F., Herrero, J., Stewart, A., Quezada, S. A., Peggs, K. S., Van Loo, P., Dive, C., Lin, C. J., Rabinowitz, M., Aerts, H., Hackshaw, A., Shaw, J. A., Zimmermann, B. G., consortium, T. R., consortium, P., and Swanton, C. (2017) Phylogenetic ctDNA analysis depicts early-stage lung cancer evolution. *Nature* **545**, 446-451
17. Jamal-Hanjani, M., Wilson, G. A., McGranahan, N., Birkbak, N. J., Watkins, T. B. K., Veeriah, S., Shafi, S., Johnson, D. H., Mitter, R., Rosenthal, R., Salm, M., Horswell, S., Escudero, M., Matthews, N., Rowan, A., Chambers, T., Moore, D. A., Turajlic, S., Xu, H., Lee, S. M., Forster, M. D., Ahmad, T., Hiley, C. T., Abbosh, C., Falzon, M., Borg, E., Marafioti, T., Lawrence, D., Hayward, M., Kolvekar, S., Panagiotopoulos, N., Janes, S. M., Thakrar, R., Ahmed, A., Blackhall, F., Summers, Y., Shah, R., Joseph, L., Quinn, A. M., Crosbie, P. A., Naidu, B., Middleton, G., Langman, G., Trotter, S., Nicolson, M., Remmen, H., Kerr, K., Chetty, M., Gomersall, L., Fennell, D. A., Nakas, A., Rathinam, S., Anand, G., Khan, S., Russell, P., Ezhil, V., Ismail, B., Irvin-Sellers, M., Prakash, V., Lester, J. F., Kornaszewska, M., Attanoos, R., Adams, H., Davies, H., Dentro, S., Taniere, P., O'Sullivan, B., Lowe, H. L., Hartley, J. A., Iles, N., Bell, H., Ngai, Y., Shaw, J. A., Herrero, J., Szallasi, Z., Schwarz, R. F., Stewart, A., Quezada, S. A., Le Quesne, J., Van Loo, P., Dive, C., Hackshaw, A., Swanton, C., and Consortium, T. R. (2017) Tracking the Evolution of Non-Small-Cell Lung Cancer. *N Engl J Med* **376**, 2109-2121
18. Hoadley, K. A., Yau, C., Hinoue, T., Wolf, D. M., Lazar, A. J., Drill, E., Shen, R., Taylor, A. M., Cherniack, A. D., Thorsson, V., Akbani, R., Bowlby, R., Wong, C. K., Wiznerowicz, M., Sanchez-Vega, F., Robertson, A. G., Schneider, B. G., Lawrence, M. S., Noushmehr, H., Malta, T. M., Cancer Genome Atlas, N., Stuart, J. M., Benz, C. C., and Laird, P. W. (2018) Cell-of-Origin Patterns Dominate the Molecular Classification of 10,000 Tumors from 33 Types of Cancer. *Cell* **173**, 291-304 e296
19. Liu, J., Lichtenberg, T., Hoadley, K. A., Poisson, L. M., Lazar, A. J., Cherniack, A. D., Kovatich, A. J., Benz, C. C., Levine, D. A., Lee, A. V., Omberg, L., Wolf, D. M., Shriver, C. D., Thorsson, V., Cancer Genome Atlas Research, N., and Hu, H. (2018) An Integrated TCGA Pan-Cancer Clinical Data Resource to Drive High-Quality Survival Outcome Analytics. *Cell* **173**, 400-416 e411
20. Gao, Q., Liang, W. W., Foltz, S. M., Mutharasu, G., Jayasinghe, R. G., Cao, S., Liao, W. W., Reynolds, S. M., Wyczalkowski, M. A., Yao, L., Yu, L., Sun, S. Q., Fusion Analysis

Working, G., Cancer Genome Atlas Research, N., Chen, K., Lazar, A. J., Fields, R. C., Wendl, M. C., Van Tine, B. A., Vij, R., Chen, F., Nykter, M., Shmulevich, I., and Ding, L. (2018) Driver Fusions and Their Implications in the Development and Treatment of Human Cancers. *Cell Rep* **23**, 227-238 e223

21. Sanchez-Vega, F., Mina, M., Armenia, J., Chatila, W. K., Luna, A., La, K. C., Dimitriadoy, S., Liu, D. L., Kantheti, H. S., Saghafinia, S., Chakravarty, D., Daian, F., Gao, Q., Bailey, M. H., Liang, W. W., Foltz, S. M., Shmulevich, I., Ding, L., Heins, Z., Ochoa, A., Gross, B., Gao, J., Zhang, H., Kundra, R., Kandoth, C., Bahceci, I., Dervishi, L., Dogrusoz, U., Zhou, W., Shen, H., Laird, P. W., Way, G. P., Greene, C. S., Liang, H., Xiao, Y., Wang, C., Iavarone, A., Berger, A. H., Bivona, T. G., Lazar, A. J., Hammer, G. D., Giordano, T., Kwong, L. N., McArthur, G., Huang, C., Tward, A. D., Frederick, M. J., McCormick, F., Meyerson, M., Cancer Genome Atlas Research, N., Van Allen, E. M., Cherniack, A. D., Ciriello, G., Sander, C., and Schultz, N. (2018) Oncogenic Signaling Pathways in The Cancer Genome Atlas. *Cell* **173**, 321-337 e310
22. Taylor, A. M., Shih, J., Ha, G., Gao, G. F., Zhang, X., Berger, A. C., Schumacher, S. E., Wang, C., Hu, H., Liu, J., Lazar, A. J., Cancer Genome Atlas Research, N., Cherniack, A. D., Beroukhim, R., and Meyerson, M. (2018) Genomic and Functional Approaches to Understanding Cancer Aneuploidy. *Cancer Cell* **33**, 676-689 e673
23. Ellrott, K., Bailey, M. H., Saksena, G., Covington, K. R., Kandoth, C., Stewart, C., Hess, J., Ma, S., Chiotti, K. E., McLellan, M., Sofia, H. J., Hutter, C., Getz, G., Wheeler, D., Ding, L., Group, M. C. W., and Cancer Genome Atlas Research, N. (2018) Scalable Open Science Approach for Mutation Calling of Tumor Exomes Using Multiple Genomic Pipelines. *Cell Syst* **6**, 271-281 e277
24. Oberg, J. A., Glade Bender, J. L., Sulis, M. L., Pendrick, D., Sireci, A. N., Hsiao, S. J., Turk, A. T., Dela Cruz, F. S., Hibshoosh, H., Remotti, H., Zylber, R. J., Pang, J., Diolaiti, D., Koval, C., Andrews, S. J., Garvin, J. H., Yamashiro, D. J., Chung, W. K., Emerson, S. G., Nagy, P. L., Mansukhani, M. M., and Kung, A. L. (2016) Implementation of next generation sequencing into pediatric hematology-oncology practice: moving beyond actionable alterations. *Genome Med* **8**, 133
25. Bhandari, V., Hoey, C., Liu, L. Y., Lalonde, E., Ray, J., Livingstone, J., Lesurf, R., Shiah, Y. J., Vujcic, T., Huang, X., Espiritu, S. M. G., Heisler, L. E., Yousif, F., Huang, V., Yamaguchi, T. N., Yao, C. Q., Sabelnykova, V. Y., Fraser, M., Chua, M. L. K., van der Kwast, T., Liu, S. K., Boutros, P. C., and Bristow, R. G. (2019) Molecular landmarks of tumor hypoxia across cancer types. *Nat Genet* **51**, 308-318
26. Jusakul, A., Cutcutache, I., Yong, C. H., Lim, J. Q., Huang, M. N., Padmanabhan, N., Nellore, V., Kongpetch, S., Ng, A. W. T., Ng, L. M., Choo, S. P., Myint, S. S., Thanan, R., Nagarajan, S., Lim, W. K., Ng, C. C. Y., Boot, A., Liu, M., Ong, C. K., Rajasegaran, V., Lie, S., Lim, A. S. T., Lim, T. H., Tan, J., Loh, J. L., McPherson, J. R., Khuntikeo, N., Bhudhisawasdi, V., Yongvanit, P., Wongkham, S., Totoki, Y., Nakamura, H., Arai, Y., Yamasaki, S., Chow, P. K., Chung, A. Y. F., Ooi, L., Lim, K. H., Dima, S., Duda, D. G., Popescu, I., Broet, P., Hsieh, S. Y., Yu, M. C., Scarpa, A., Lai, J., Luo, D. X., Carvalho, A. L., Vettore, A. L., Rhee, H., Park, Y. N., Alexandrov, L. B., Gordan, R., Rozen, S. G., Shibata, T., Pairojkul, C., Teh, B. T., and Tan, P. (2017) Whole-Genome and Epigenomic Landscapes of Etiologically Distinct Subtypes of Cholangiocarcinoma. *Cancer Discov* **7**, 1116-1135
27. Bonilla, X., Parmentier, L., King, B., Bezrukov, F., Kaya, G., Zoete, V., Seplyarskiy, V. B., Sharpe, H. J., McKee, T., Letourneau, A., Ribaux, P. G., Popadin, K., Basset-Seguin, N., Ben Chaabene, R., Santoni, F. A., Andrianova, M. A., Guipponi, M., Garieri, M., Verdan, C., Grosdemange, K., Sumara, O., Eilers, M., Aifantis, I., Michielin, O., de Sauvage, F. J., Antonarakis, S. E., and Nikolaev, S. I. (2016) Genomic analysis identifies

new drivers and progression pathways in skin basal cell carcinoma. *Nat Genet* **48**, 398-406

28. Vasaikar, S., Huang, C., Wang, X., Petyuk, V. A., Savage, S. R., Wen, B., Dou, Y., Zhang, Y., Shi, Z., Arshad, O. A., Gritsenko, M. A., Zimmerman, L. J., McDermott, J. E., Clauss, T. R., Moore, R. J., Zhao, R., Monroe, M. E., Wang, Y. T., Chambers, M. C., Slebos, R. J. C., Lau, K. S., Mo, Q., Ding, L., Ellis, M., Thiagarajan, M., Kinsinger, C. R., Rodriguez, H., Smith, R. D., Rodland, K. D., Liebler, D. C., Liu, T., Zhang, B., and Clinical Proteomic Tumor Analysis, C. (2019) Proteogenomic Analysis of Human Colon Cancer Reveals New Therapeutic Opportunities. *Cell* **177**, 1035-1049 e1019
29. Jones, S., Stransky, N., McCord, C. L., Cerami, E., Lagowski, J., Kelly, D., Angiuoli, S. V., Sausen, M., Kann, L., Shukla, M., Makar, R., Wood, L. D., Diaz, L. A., Jr., Lengauer, C., and Velculescu, V. E. (2014) Genomic analyses of gynaecologic carcinosarcomas reveal frequent mutations in chromatin remodelling genes. *Nat Commun* **5**, 5006
30. Manning-Geist, B. L., Liu, Y. L., Devereaux, K. A., Paula, A. D. C., Zhou, Q. C., Ma, W., Selenica, P., Ceyhan-Birsoy, O., Moukarzel, L. A., Hoang, T., Gordhandas, S., Rubinstein, M. M., Friedman, C. F., Aghajanian, C., Abu-Rustum, N. R., Stadler, Z. K., Reis-Filho, J. S., Iasonos, A., Zamarin, D., Ellenson, L. H., Lakhman, Y., Mandelker, D. L., and Weigelt, B. (2022) Microsatellite Instability-High Endometrial Cancers with MLH1 Promoter Hypermethylation Have Distinct Molecular and Clinical Profiles. *Clin Cancer Res* **28**, 4302-4311
31. Morin, R. D., Mungall, K., Pleasance, E., Mungall, A. J., Goya, R., Huff, R. D., Scott, D. W., Ding, J., Roth, A., Chiu, R., Corbett, R. D., Chan, F. C., Mendez-Lago, M., Trinh, D. L., Bolger-Munro, M., Taylor, G., Hadj Khodabakhshi, A., Ben-Neriah, S., Pon, J., Meissner, B., Woolcock, B., Farnoud, N., Rogic, S., Lim, E. L., Johnson, N. A., Shah, S., Jones, S., Steidl, C., Holt, R., Birol, I., Moore, R., Connors, J. M., Gascoyne, R. D., and Marra, M. A. (2013) Mutational and structural analysis of diffuse large B-cell lymphoma using whole-genome sequencing. *Blood* **122**, 1256-1265
32. Robinson, D. R., Wu, Y. M., Lonigro, R. J., Vats, P., Cobain, E., Everett, J., Cao, X., Rabban, E., Kumar-Sinha, C., Raymond, V., Schuetze, S., Alva, A., Siddiqui, J., Chugh, R., Worden, F., Zalupski, M. M., Innis, J., Mody, R. J., Tomlins, S. A., Lucas, D., Baker, L. H., Ramnath, N., Schott, A. F., Hayes, D. F., Vijai, J., Offit, K., Stoffel, E. M., Roberts, J. S., Smith, D. C., Kunju, L. P., Talpaz, M., Cieslik, M., and Chinnaiyan, A. M. (2017) Integrative clinical genomics of metastatic cancer. *Nature* **548**, 297-303
33. Consortium, I. T. P.-C. A. o. W. G. (2020) Pan-cancer analysis of whole genomes. *Nature* **578**, 82-93
34. Ashley, C. W., Selenica, P., Patel, J., Wu, M., Nincevic, J., Lakhman, Y., Zhou, Q., Shah, R. H., Berger, M. F., Da Cruz Paula, A., Brown, D. N., Marra, A., Iasonos, A., Momeni-Boroujeni, A., Alektiar, K. M., Long Roche, K., Zivanovic, O., Mueller, J. J., Zamarin, D., Broach, V. A., Sonoda, Y., Leitao, M. M., Friedman, C. F., Jewell, E., Reis-Filho, J. S., Ellenson, L. H., Aghajanian, C., Abu-Rustum, N. R., Cadoo, K., and Weigelt, B. (2023) High-Sensitivity Mutation Analysis of Cell-Free DNA for Disease Monitoring in Endometrial Cancer. *Clin Cancer Res* **29**, 410-421
35. Nixon, M. J., Formisano, L., Mayer, I. A., Estrada, M. V., Gonzalez-Ericsson, P. I., Isakoff, S. J., Forero-Torres, A., Won, H., Sanders, M. E., Solit, D. B., Berger, M. F., Cantley, L. C., Winer, E. P., Arteaga, C. L., and Balko, J. M. (2019) PIK3CA and MAP3K1 alterations imply luminal A status and are associated with clinical benefit from pan-PI3K inhibitor buparlisib and letrozole in ER+ metastatic breast cancer. *NPJ Breast Cancer* **5**, 31
36. Hodis, E., Watson, I. R., Kryukov, G. V., Arold, S. T., Imielinski, M., Theurillat, J. P., Nickerson, E., Auclair, D., Li, L., Place, C., Dicara, D., Ramos, A. H., Lawrence, M. S., Cibulskis, K., Sivachenko, A., Voet, D., Saksena, G., Stransky, N., Onofrio, R. C.,

Winckler, W., Ardlie, K., Wagle, N., Wargo, J., Chong, K., Morton, D. L., Stemke-Hale, K., Chen, G., Noble, M., Meyerson, M., Ladbury, J. E., Davies, M. A., Gershenwald, J. E., Wagner, S. N., Hoon, D. S., Schadendorf, D., Lander, E. S., Gabriel, S. B., Getz, G., Garraway, L. A., and Chin, L. (2012) A landscape of driver mutations in melanoma. *Cell* **150**, 251-263

37. Gardner, E. E., Lok, B. H., Schneeberger, V. E., Desmeules, P., Miles, L. A., Arnold, P. K., Ni, A., Khodos, I., de Stanchina, E., Nguyen, T., Sage, J., Campbell, J. E., Ribich, S., Rekhtman, N., Dowlati, A., Massion, P. P., Rudin, C. M., and Poirier, J. T. (2017) Chemosensitive Relapse in Small Cell Lung Cancer Proceeds through an EZH2-SLFN11 Axis. *Cancer Cell* **31**, 286-299
38. Hugo, W., Zaretsky, J. M., Sun, L., Song, C., Moreno, B. H., Hu-Lieskovan, S., Berent-Maoz, B., Pang, J., Chmielowski, B., Cherry, G., Seja, E., Lomeli, S., Kong, X., Kelley, M. C., Sosman, J. A., Johnson, D. B., Ribas, A., and Lo, R. S. (2016) Genomic and Transcriptomic Features of Response to Anti-PD-1 Therapy in Metastatic Melanoma. *Cell* **165**, 35-44
39. Faltas, B. M., Prandi, D., Tagawa, S. T., Molina, A. M., Nanus, D. M., Sternberg, C., Rosenberg, J., Mosquera, J. M., Robinson, B., Elemento, O., Sboner, A., Beltran, H., Demichelis, F., and Rubin, M. A. (2016) Clonal evolution of chemotherapy-resistant urothelial carcinoma. *Nat Genet* **48**, 1490-1499
40. Van Allen, E. M., Mouw, K. W., Kim, P., Iyer, G., Wagle, N., Al-Ahmadie, H., Zhu, C., Ostrovnaya, I., Kryukov, G. V., O'Connor, K. W., Sfakianos, J., Garcia-Grossman, I., Kim, J., Guancial, E. A., Bambury, R., Bahl, S., Gupta, N., Farlow, D., Qu, A., Signoretti, S., Barletta, J. A., Reuter, V., Boehm, J., Lawrence, M., Getz, G., Kantoff, P., Bochner, B. H., Choueiri, T. K., Bajorin, D. F., Solit, D. B., Gabriel, S., D'Andrea, A., Garraway, L. A., and Rosenberg, J. E. (2014) Somatic ERCC2 mutations correlate with cisplatin sensitivity in muscle-invasive urothelial carcinoma. *Cancer Discov* **4**, 1140-1153
41. Chen, K., Yang, D., Li, X., Sun, B., Song, F., Cao, W., Brat, D. J., Gao, Z., Li, H., Liang, H., Zhao, Y., Zheng, H., Li, M., Buckner, J., Patterson, S. D., Ye, X., Reinhard, C., Bhathena, A., Joshi, D., Mischel, P. S., Croce, C. M., Wang, Y. M., Raghavakaimal, S., Li, H., Lu, X., Pan, Y., Chang, H., Ba, S., Luo, L., Cavenee, W. K., Zhang, W., and Hao, X. (2015) Mutational landscape of gastric adenocarcinoma in Chinese: implications for prognosis and therapy. *Proc Natl Acad Sci U S A* **112**, 1107-1112
42. Abida, W., Cyrta, J., Heller, G., Prandi, D., Armenia, J., Coleman, I., Cieslik, M., Benelli, M., Robinson, D., Van Allen, E. M., Sboner, A., Fedrizzi, T., Mosquera, J. M., Robinson, B. D., De Sarkar, N., Kunju, L. P., Tomlins, S., Wu, Y. M., Nava Rodrigues, D., Loda, M., Gopalan, A., Reuter, V. E., Pritchard, C. C., Mateo, J., Bianchini, D., Miranda, S., Carreira, S., Rescigno, P., Filipenko, J., Vinson, J., Montgomery, R. B., Beltran, H., Heath, E. I., Scher, H. I., Kantoff, P. W., Taplin, M. E., Schultz, N., deBono, J. S., Demichelis, F., Nelson, P. S., Rubin, M. A., Chinnaiyan, A. M., and Sawyers, C. L. (2019) Genomic correlates of clinical outcome in advanced prostate cancer. *Proc Natl Acad Sci U S A* **116**, 11428-11436
43. Xue, R., Chen, L., Zhang, C., Fujita, M., Li, R., Yan, S. M., Ong, C. K., Liao, X., Gao, Q., Sasagawa, S., Li, Y., Wang, J., Guo, H., Huang, Q. T., Zhong, Q., Tan, J., Qi, L., Gong, W., Hong, Z., Li, M., Zhao, J., Peng, T., Lu, Y., Lim, K. H. T., Boot, A., Ono, A., Chayama, K., Zhang, Z., Rozen, S. G., Teh, B. T., Wang, X. W., Nakagawa, H., Zeng, M. S., Bai, F., and Zhang, N. (2019) Genomic and Transcriptomic Profiling of Combined Hepatocellular and Intrahepatic Cholangiocarcinoma Reveals Distinct Molecular Subtypes. *Cancer Cell* **35**, 932-947 e938
44. Seshagiri, S., Stawiski, E. W., Durinck, S., Modrusan, Z., Storm, E. E., Conboy, C. B., Chaudhuri, S., Guan, Y., Janakiraman, V., Jaiswal, B. S., Guillory, J., Ha, C., Dijkgraaf, G. J., Stinson, J., Gnad, F., Huntley, M. A., Degenhardt, J. D., Haverty, P. M., Bourgon,

R., Wang, W., Koeppen, H., Gentleman, R., Starr, T. K., Zhang, Z., Largaespada, D. A., Wu, T. D., and de Sauvage, F. J. (2012) Recurrent R-spondin fusions in colon cancer. *Nature* **488**, 660-664

45. Rodriguez-Calero, A., Gallon, J., Akhoundova, D., Maletti, S., Ferguson, A., Cyrta, J., Amstutz, U., Garofoli, A., Paradiso, V., Tomlins, S. A., Hewer, E., Genitsch, V., Fleischmann, A., Vassella, E., Rushing, E. J., Grobholz, R., Fischer, I., Jochum, W., Cathomas, G., Osunkoya, A. O., Bubendorf, L., Moch, H., Thalmann, G., Ng, C. K. Y., Gillessen, S., Piscuoglio, S., and Rubin, M. A. (2022) Alterations in homologous recombination repair genes in prostate cancer brain metastases. *Nat Commun* **13**, 2400
46. Krauthammer, M., Kong, Y., Ha, B. H., Evans, P., Bacchiocchi, A., McCusker, J. P., Cheng, E., Davis, M. J., Goh, G., Choi, M., Ariyan, S., Narayan, D., Dutton-Regester, K., Capatana, A., Holman, E. C., Bosenberg, M., Sznol, M., Kluger, H. M., Brash, D. E., Stern, D. F., Materin, M. A., Lo, R. S., Mane, S., Ma, S., Kidd, K. K., Hayward, N. K., Lifton, R. P., Schlessinger, J., Boggon, T. J., and Halaban, R. (2012) Exome sequencing identifies recurrent somatic RAC1 mutations in melanoma. *Nat Genet* **44**, 1006-1014
47. Dou, Y., Kawaler, E. A., Cui Zhou, D., Gritsenko, M. A., Huang, C., Blumenberg, L., Karpova, A., Petyuk, V. A., Savage, S. R., Satpathy, S., Liu, W., Wu, Y., Tsai, C. F., Wen, B., Li, Z., Cao, S., Moon, J., Shi, Z., Cornwell, M., Wyczalkowski, M. A., Chu, R. K., Vasaikar, S., Zhou, H., Gao, Q., Moore, R. J., Li, K., Sethuraman, S., Monroe, M. E., Zhao, R., Heiman, D., Krug, K., Clauser, K., Kothadia, R., Maruvka, Y., Pico, A. R., Oliphant, A. E., Hoskins, E. L., Pugh, S. L., Beecroft, S. J. I., Adams, D. W., Jarman, J. C., Kong, A., Chang, H. Y., Reva, B., Liao, Y., Rykunov, D., Colaprico, A., Chen, X. S., Czekanski, A., Jedryka, M., Matkowski, R., Wiznerowicz, M., Hiltke, T., Boja, E., Kinsinger, C. R., Mesri, M., Robles, A. I., Rodriguez, H., Mutch, D., Fuh, K., Ellis, M. J., DeLair, D., Thiagarajan, M., Mani, D. R., Getz, G., Noble, M., Nesvizhskii, A. I., Wang, P., Anderson, M. L., Levine, D. A., Smith, R. D., Payne, S. H., Ruggles, K. V., Rodland, K. D., Ding, L., Zhang, B., Liu, T., Fenyo, D., and Clinical Proteomic Tumor Analysis, C. (2020) Proteogenomic Characterization of Endometrial Carcinoma. *Cell* **180**, 729-748 e726
48. Ahn, S. M., Jang, S. J., Shim, J. H., Kim, D., Hong, S. M., Sung, C. O., Baek, D., Haq, F., Ansari, A. A., Lee, S. Y., Chun, S. M., Choi, S., Choi, H. J., Kim, J., Kim, S., Hwang, S., Lee, Y. J., Lee, J. E., Jung, W. R., Jang, H. Y., Yang, E., Sung, W. K., Lee, N. P., Mao, M., Lee, C., Zucman-Rossi, J., Yu, E., Lee, H. C., and Kong, G. (2014) Genomic portrait of resectable hepatocellular carcinomas: implications of RB1 and FGF19 aberrations for patient stratification. *Hepatology* **60**, 1972-1982
49. Barthel, F. P., Johnson, K. C., Varn, F. S., Moskalik, A. D., Tanner, G., Kocakavuk, E., Anderson, K. J., Abiola, O., Aldape, K., Alfaro, K. D., Alpar, D., Amin, S. B., Ashley, D. M., Bandopadhayay, P., Barnholtz-Sloan, J. S., Beroukhim, R., Bock, C., Brastianos, P. K., Brat, D. J., Brodbelt, A. R., Bruns, A. F., Bulsara, K. R., Chakrabarty, A., Chakravarti, A., Chuang, J. H., Claus, E. B., Cochran, E. J., Connelly, J., Costello, J. F., Finocchiaro, G., Fletcher, M. N., French, P. J., Gan, H. K., Gilbert, M. R., Gould, P. V., Grimmer, M. R., Iavarone, A., Ismail, A., Jenkinson, M. D., Khasraw, M., Kim, H., Kouwenhoven, M. C. M., LaViolette, P. S., Li, M., Lichter, P., Ligon, K. L., Lowman, A. K., Malta, T. M., Mazor, T., McDonald, K. L., Molinaro, A. M., Nam, D. H., Nayyar, N., Ng, H. K., Ngan, C. Y., Niclou, S. P., Niers, J. M., Noushmehr, H., Noorbakhsh, J., Ormond, D. R., Park, C. K., Poisson, L. M., Rabadan, R., Radlwimmer, B., Rao, G., Reifenberger, G., Sa, J. K., Schuster, M., Shaw, B. L., Short, S. C., Smitt, P. A. S., Sloan, A. E., Smits, M., Suzuki, H., Tabatabai, G., Van Meir, E. G., Watts, C., Weller, M., Wesseling, P., Westerman, B. A., Widhalm, G., Woehrer, A., Yung, W. K. A., Zadeh, G., Huse, J. T., De Groot, J. F., Stead, L. F., Verhaak, R. G. W., and Consortium, G. (2019) Longitudinal molecular trajectories of diffuse glioma in adults. *Nature* **576**, 112-120

50. Alatise, O. I., Knapp, G. C., Sharma, A., Chatila, W. K., Arowolo, O. A., Olasehinde, O., Famurewa, O. C., Omisore, A. D., Komolafe, A. O., Olaofe, O. O., Katung, A. I., Ibikunle, D. E., Egberongbe, A. A., Olatoke, S. A., Agodirin, S. O., Adesiyun, O. A., Adeyeye, A., Kolawole, O. A., Olakanmi, A. O., Arora, K., Constable, J., Shah, R., Basunia, A., Sylvester, B., Wu, C., Weiser, M. R., Seier, K., Gonen, M., Stadler, Z. K., Kemel, Y., Vakiani, E., Berger, M. F., Chan, T. A., Solit, D. B., Shia, J., Sanchez-Vega, F., Schultz, N., Brennan, M., Smith, J. J., and Kingham, T. P. (2021) Molecular and phenotypic profiling of colorectal cancer patients in West Africa reveals biological insights. *Nat Commun* **12**, 6821
51. Satpathy, S., Krug, K., Jean Beltran, P. M., Savage, S. R., Petralia, F., Kumar-Sinha, C., Dou, Y., Reva, B., Kane, M. H., Avanessian, S. C., Vasaikar, S. V., Krek, A., Lei, J. T., Jaehnig, E. J., Omelchenko, T., Geffen, Y., Bergstrom, E. J., Stathias, V., Christianson, K. E., Heiman, D. I., Cieslik, M. P., Cao, S., Song, X., Ji, J., Liu, W., Li, K., Wen, B., Li, Y., Gumus, Z. H., Selvan, M. E., Soundararajan, R., Visal, T. H., Raso, M. G., Parra, E. R., Babur, O., Vats, P., Anand, S., Schraink, T., Cornwell, M., Rodrigues, F. M., Zhu, H., Mo, C. K., Zhang, Y., da Veiga Leprevost, F., Huang, C., Chinnaiyan, A. M., Wyczalkowski, M. A., Omenn, G. S., Newton, C. J., Schurer, S., Ruggles, K. V., Fenyo, D., Jewell, S. D., Thiagarajan, M., Mesri, M., Rodriguez, H., Mani, S. A., Udeshi, N. D., Getz, G., Suh, J., Li, Q. K., Hostetter, G., Paik, P. K., Dhanasekaran, S. M., Govindan, R., Ding, L., Robles, A. I., Clauser, K. R., Nesvizhskii, A. I., Wang, P., Carr, S. A., Zhang, B., Mani, D. R., Gillette, M. A., and Clinical Proteomic Tumor Analysis, C. (2021) A proteogenomic portrait of lung squamous cell carcinoma. *Cell* **184**, 4348-4371 e4340
52. Gingras, M. C., Covington, K. R., Chang, D. K., Donehower, L. A., Gill, A. J., Ittmann, M. M., Creighton, C. J., Johns, A. L., Shinbrot, E., Dewal, N., Fisher, W. E., Australian Pancreatic Cancer Genome, I., Pilarsky, C., Grutzmann, R., Overman, M. J., Jamieson, N. B., Van Buren, G., 2nd, Drummond, J., Walker, K., Hampton, O. A., Xi, L., Muzny, D. M., Doddapaneni, H., Lee, S. L., Bellair, M., Hu, J., Han, Y., Dinh, H. H., Dahdouli, M., Samra, J. S., Bailey, P., Waddell, N., Pearson, J. V., Harliwong, I., Wang, H., Aust, D., Oien, K. A., Hruban, R. H., Hodges, S. E., McElhany, A., Saengboonmee, C., Duthie, F. R., Grimmond, S. M., Biankin, A. V., Wheeler, D. A., and Gibbs, R. A. (2016) Ampullary Cancers Harbor ELF3 Tumor Suppressor Gene Mutations and Exhibit Frequent WNT Dysregulation. *Cell Rep* **14**, 907-919
53. Chen, B., Scurrah, C. R., McKinley, E. T., Simmons, A. J., Ramirez-Solano, M. A., Zhu, X., Markham, N. O., Heiser, C. N., Vega, P. N., Rolong, A., Kim, H., Sheng, Q., Drewes, J. L., Zhou, Y., Southard-Smith, A. N., Xu, Y., Ro, J., Jones, A. L., Revetta, F., Berry, L. D., Niitsu, H., Islam, M., Pelka, K., Hofree, M., Chen, J. H., Sarkizova, S., Ng, K., Giannakis, M., Boland, G. M., Aguirre, A. J., Anderson, A. C., Rozenblatt-Rosen, O., Regev, A., Hacohen, N., Kawasaki, K., Sato, T., Goettel, J. A., Grady, W. M., Zheng, W., Washington, M. K., Cai, Q., Sears, C. L., Goldenring, J. R., Franklin, J. L., Su, T., Huh, W. J., Vandekar, S., Roland, J. T., Liu, Q., Coffey, R. J., Shrubsole, M. J., and Lau, K. S. (2021) Differential pre-malignant programs and microenvironment chart distinct paths to malignancy in human colorectal polyps. *Cell* **184**, 6262-6280 e6226
54. Lohr, J. G., Stojanov, P., Carter, S. L., Cruz-Gordillo, P., Lawrence, M. S., Auclair, D., Sougnez, C., Knoechel, B., Gould, J., Saksena, G., Cibulskis, K., McKenna, A., Chapman, M. A., Straussman, R., Levy, J., Perkins, L. M., Keats, J. J., Schumacher, S. E., Rosenberg, M., Multiple Myeloma Research, C., Getz, G., and Golub, T. R. (2014) Widespread genetic heterogeneity in multiple myeloma: implications for targeted therapy. *Cancer Cell* **25**, 91-101
55. Tang, J., Fewings, E., Chang, D., Zeng, H., Liu, S., Jorapur, A., Belote, R. L., McNeal, A. S., Tan, T. M., Yeh, I., Arron, S. T., Judson-Torres, R. L., Bastian, B. C., and Shain, A.

H. (2020) The genomic landscapes of individual melanocytes from human skin. *Nature* **586**, 600-605
56. Gerami, P., Tandukar, B., Deivendran, D., Olivares, S., Chen, L., Tang, J., Tan, T., Sharma, H., Bandari, A. K., Cruz-Pacheco, N., Chang, D., Marty, A., Olshen, A., Murad, N. F., Song, J., Lee, J., Yeh, I., and Hunter Shain, A. (2024) Molecular effects of indoor tanning. *bioRxiv*
57. Tandukar, B., Deivendran, D., Chen, L., Bahrani, N., Weier, B., Sharma, H., Cruz-Pacheco, N., Hu, M., Marks, K., Zitnay, R. G., Bandari, A. K., Nekoonam, R., Yeh, I., Judson-Torres, R., and Shain, A. H. (2025) Somatic mutations distinguish melanocyte subpopulations in human skin. *bioRxiv*
58. Tandukar, B., Deivendran, D., Chen, L., Cruz-Pacheco, N., Sharma, H., Xu, A., Bandari, A. K., Chen, D. B., George, C., Marty, A., Cho, R. J., Cheng, J., Saylor, D., Gerami, P., Arron, S. T., Bastian, B. C., and Shain, A. H. (2024) Genetic evolution of keratinocytes to cutaneous squamous cell carcinoma. *bioRxiv*
59. Le Gallo, M., Rudd, M. L., Urick, M. E., Hansen, N. F., Zhang, S., Program, N. C. S., Lozy, F., Sgroi, D. C., Vidal Bel, A., Matias-Guiu, X., Broaddus, R. R., Lu, K. H., Levine, D. A., Mutch, D. G., Goodfellow, P. J., Salvesen, H. B., Mullikin, J. C., and Bell, D. W. (2017) Somatic mutation profiles of clear cell endometrial tumors revealed by whole exome and targeted gene sequencing. *Cancer* **123**, 3261-3268
60. Kim, K., Hu, W., Audenet, F., Almassi, N., Hanrahan, A. J., Murray, K., Bagrodia, A., Wong, N., Clinton, T. N., Dason, S., Mohan, V., Jebiwott, S., Nagar, K., Gao, J., Penson, A., Hughes, C., Gordon, B., Chen, Z., Dong, Y., Watson, P. A., Alvim, R., Elzein, A., Gao, S. P., Cocco, E., Santin, A. D., Ostrovnaya, I., Hsieh, J. J., Sagi, I., Pietzak, E. J., Hakimi, A. A., Rosenberg, J. E., Iyer, G., Vargas, H. A., Scaltriti, M., Al-Ahmadie, H., Solit, D. B., and Coleman, J. A. (2020) Modeling biological and genetic diversity in upper tract urothelial carcinoma with patient derived xenografts. *Nat Commun* **11**, 1975
61. Chen, L., Zhang, C., Xue, R., Liu, M., Bai, J., Bao, J., Wang, Y., Jiang, N., Li, Z., Wang, W., Wang, R., Zheng, B., Yang, A., Hu, J., Liu, K., Shen, S., Zhang, Y., Bai, M., Wang, Y., Zhu, Y., Yang, S., Gao, Q., Gu, J., Gao, D., Wang, X. W., Nakagawa, H., Zhang, N., Wu, L., Rozen, S. G., Bai, F., and Wang, H. (2024) Deep whole-genome analysis of 494 hepatocellular carcinomas. *Nature* **627**, 586-593
62. Miao, D., Margolis, C. A., Vokes, N. I., Liu, D., Taylor-Weiner, A., Wankowicz, S. M., Adeegbe, D., Keliher, D., Schilling, B., Tracy, A., Manos, M., Chau, N. G., Hanna, G. J., Polak, P., Rodig, S. J., Signoretti, S., Sholl, L. M., Engelman, J. A., Getz, G., Janne, P. A., Haddad, R. I., Choueiri, T. K., Barbie, D. A., Haq, R., Awad, M. M., Schadendorf, D., Hodi, F. S., Bellmunt, J., Wong, K. K., Hammerman, P., and Van Allen, E. M. (2018) Genomic correlates of response to immune checkpoint blockade in microsatellite-stable solid tumors. *Nat Genet* **50**, 1271-1281
63. Roelands, J., Kuppen, P. J. K., Ahmed, E. I., Mall, R., Masoodi, T., Singh, P., Monaco, G., Raynaud, C., de Miranda, N., Ferraro, L., Carneiro-Lobo, T. C., Syed, N., Rawat, A., Awad, A., Decock, J., Mifsud, W., Miller, L. D., Sherif, S., Mohamed, M. G., Rinchai, D., Van den Eynde, M., Sayaman, R. W., Ziv, E., Bertucci, F., Petkar, M. A., Lorenz, S., Mathew, L. S., Wang, K., Murugesan, S., Chaussabel, D., Vahrmeijer, A. L., Wang, E., Ceccarelli, A., Fakhro, K. A., Zoppoli, G., Ballestrero, A., Tollenaar, R., Marincola, F. M., Galon, J., Khodor, S. A., Ceccarelli, M., Hendrickx, W., and Bedognetti, D. (2023) An integrated tumor, immune and microbiome atlas of colon cancer. *Nat Med* **29**, 1273-1286
64. Al Shihabi, A., Tebon, P. J., Nguyen, H. T. L., Chantharasamee, J., Sartini, S., Davarifar, A., Jensen, A. Y., Diaz-Infante, M., Cox, H., Gonzalez, A. E., Norris, S., Sperry, J., Nakashima, J., Tavanaie, N., Winata, H., Fitz-Gibbon, S. T., Yamaguchi, T. N., Jeong, J. H., Dry, S., Singh, A. S., Chmielowski, B., Crompton, J. G., Kalbasi, A. K., Eilber, F. C.,

Hornicek, F., Bernthal, N. M., Nelson, S. D., Boutros, P. C., Federman, N. C., Yanagawa, J., and Soragni, A. (2024) The landscape of drug sensitivity and resistance in sarcoma. *Cell Stem Cell* **31**, 1524-1542 e1524

65. Miao, D., Margolis, C. A., Gao, W., Voss, M. H., Li, W., Martini, D. J., Norton, C., Bosse, D., Wankowicz, S. M., Cullen, D., Horak, C., Wind-Rotolo, M., Tracy, A., Giannakis, M., Hodi, F. S., Drake, C. G., Ball, M. W., Allaf, M. E., Snyder, A., Hellmann, M. D., Ho, T., Motzer, R. J., Signoretti, S., Kaelin, W. G., Jr., Choueiri, T. K., and Van Allen, E. M. (2018) Genomic correlates of response to immune checkpoint therapies in clear cell renal cell carcinoma. *Science* **359**, 801-806
66. Martelotto, L. G., De Filippo, M. R., Ng, C. K., Natrajan, R., Fuhrmann, L., Cyrta, J., Piscuoglio, S., Wen, H. C., Lim, R. S., Shen, R., Schultheis, A. M., Wen, Y. H., Edelweiss, M., Mariani, O., Stenman, G., Chan, T. A., Colombo, P. E., Norton, L., Vincent-Salomon, A., Reis-Filho, J. S., and Weigelt, B. (2015) Genomic landscape of adenoid cystic carcinoma of the breast. *J Pathol* **237**, 179-189
67. Ho, A. S., Ochoa, A., Jayakumaran, G., Zehir, A., Valero Mayor, C., Tepe, J., Makarov, V., Dalin, M. G., He, J., Bailey, M., Montesion, M., Ross, J. S., Miller, V. A., Chan, L., Ganly, I., Dogan, S., Katabi, N., Tsipouras, P., Ha, P., Agrawal, N., Solit, D. B., Futreal, P. A., El Naggar, A. K., Reis-Filho, J. S., Weigelt, B., Ho, A. L., Schultz, N., Chan, T. A., and Morris, L. G. (2019) Genetic hallmarks of recurrent/metastatic adenoid cystic carcinoma. *J Clin Invest* **129**, 4276-4289
68. Zou, S., Li, J., Zhou, H., Frech, C., Jiang, X., Chu, J. S., Zhao, X., Li, Y., Li, Q., Wang, H., Hu, J., Kong, G., Wu, M., Ding, C., Chen, N., and Hu, H. (2014) Mutational landscape of intrahepatic cholangiocarcinoma. *Nat Commun* **5**, 5696
69. Van Allen, E. M., Wagle, N., Sucker, A., Treacy, D. J., Johannessen, C. M., Goetz, E. M., Place, C. S., Taylor-Weiner, A., Whittaker, S., Kryukov, G. V., Hodis, E., Rosenberg, M., McKenna, A., Cibulskis, K., Farlow, D., Zimmer, L., Hillen, U., Gutzmer, R., Goldinger, S. M., Ugurel, S., Gogas, H. J., Egberts, F., Berking, C., Trefzer, U., Loquai, C., Weide, B., Hassel, J. C., Gabriel, S. B., Carter, S. L., Getz, G., Garraway, L. A., Schadendorf, D., and Dermatologic Cooperative Oncology Group of, G. (2014) The genetic landscape of clinical resistance to RAF inhibition in metastatic melanoma. *Cancer Discov* **4**, 94-109
70. George, J., Lim, J. S., Jang, S. J., Cun, Y., Ozretic, L., Kong, G., Leenders, F., Lu, X., Fernandez-Cuesta, L., Bosco, G., Muller, C., Dahmen, I., Jahchan, N. S., Park, K. S., Yang, D., Karnezis, A. N., Vaka, D., Torres, A., Wang, M. S., Korbel, J. O., Menon, R., Chun, S. M., Kim, D., Wilkerson, M., Hayes, N., Engelmann, D., Putzer, B., Bos, M., Michels, S., Vlasic, I., Seidel, D., Pinther, B., Schaub, P., Becker, C., Altmuller, J., Yokota, J., Kohno, T., Iwakawa, R., Tsuta, K., Noguchi, M., Muley, T., Hoffmann, H., Schnabel, P. A., Petersen, I., Chen, Y., Soltermann, A., Tischler, V., Choi, C. M., Kim, Y. H., Massion, P. P., Zou, Y., Jovanovic, D., Kontic, M., Wright, G. M., Russell, P. A., Solomon, B., Koch, I., Lindner, M., Muscarella, L. A., la Torre, A., Field, J. K., Jakopovic, M., Knezevic, J., Castanos-Velez, E., Roz, L., Pastorino, U., Brustugun, O. T., Lund-Iversen, M., Thunnissen, E., Kohler, J., Schuler, M., Botling, J., Sandelin, M., Sanchez-Cespedes, M., Salvesen, H. B., Achter, V., Lang, U., Bogus, M., Schneider, P. M., Zander, T., Ansen, S., Hallek, M., Wolf, J., Vingron, M., Yatabe, Y., Travis, W. D., Nurnberg, P., Reinhardt, C., Perner, S., Heukamp, L., Buttner, R., Haas, S. A., Brambilla, E., Peifer, M., Sage, J., and Thomas, R. K. (2015) Comprehensive genomic profiles of small cell lung cancer. *Nature* **524**, 47-53
71. Lee, S. H., Hu, W., Matulay, J. T., Silva, M. V., Owczarek, T. B., Kim, K., Chua, C. W., Barlow, L. J., Kandoth, C., Williams, A. B., Bergren, S. K., Pietzak, E. J., Anderson, C. B., Benson, M. C., Coleman, J. A., Taylor, B. S., Abate-Shen, C., McKiernan, J. M., Al-

Ahmadie, H., Solit, D. B., and Shen, M. M. (2018) Tumor Evolution and Drug Response in Patient-Derived Organoid Models of Bladder Cancer. *Cell* **173**, 515-528 e517

72. Petralia, F., Tignor, N., Reva, B., Koptyra, M., Chowdhury, S., Rykunov, D., Krek, A., Ma, W., Zhu, Y., Ji, J., Calinawan, A., Whiteaker, J. R., Colaprico, A., Stathias, V., Omelchenko, T., Song, X., Raman, P., Guo, Y., Brown, M. A., Ivey, R. G., Szpyt, J., Guha Thakurta, S., Gritsenko, M. A., Weitz, K. K., Lopez, G., Kalayci, S., Gumus, Z. H., Yoo, S., da Veiga Leprevost, F., Chang, H. Y., Krug, K., Katsnelson, L., Wang, Y., Kennedy, J. J., Voytovich, U. J., Zhao, L., Gaonkar, K. S., Ennis, B. M., Zhang, B., Baubet, V., Tauhid, L., Lilly, J. V., Mason, J. L., Farrow, B., Young, N., Leary, S., Moon, J., Petyuk, V. A., Nazarian, J., Adappa, N. D., Palmer, J. N., Lober, R. M., Rivero-Hinojosa, S., Wang, L. B., Wang, J. M., Broberg, M., Chu, R. K., Moore, R. J., Monroe, M. E., Zhao, R., Smith, R. D., Zhu, J., Robles, A. I., Mesri, M., Boja, E., Hiltke, T., Rodriguez, H., Zhang, B., Schadt, E. E., Mani, D. R., Ding, L., Iavarone, A., Wiznerowicz, M., Schurer, S., Chen, X. S., Heath, A. P., Rokita, J. L., Nesvizhskii, A. I., Fenyo, D., Rodland, K. D., Liu, T., Gygi, S. P., Paulovich, A. G., Resnick, A. C., Storm, P. B., Rood, B. R., Wang, P., Children's Brain Tumor, N., and Clinical Proteomic Tumor Analysis, C. (2020) Integrated Proteogenomic Characterization across Major Histological Types of Pediatric Brain Cancer. *Cell* **183**, 1962-1985 e1931
73. Painter, C. A., Jain, E., Tomson, B. N., Dunphy, M., Stoddard, R. E., Thomas, B. S., Damon, A. L., Shah, S., Kim, D., Gomez Tejeda Zanudo, J., Hornick, J. L., Chen, Y. L., Merriam, P., Raut, C. P., Demetri, G. D., Van Tine, B. A., Lander, E. S., Golub, T. R., and Wagle, N. (2020) The Angiosarcoma Project: enabling genomic and clinical discoveries in a rare cancer through patient-partnered research. *Nat Med* **26**, 181-187
74. Gillette, M. A., Satpathy, S., Cao, S., Dhanasekaran, S. M., Vasaikar, S. V., Krug, K., Petralia, F., Li, Y., Liang, W. W., Reva, B., Krek, A., Ji, J., Song, X., Liu, W., Hong, R., Yao, L., Blumenberg, L., Savage, S. R., Wendl, M. C., Wen, B., Li, K., Tang, L. C., MacMullan, M. A., Avanessian, S. C., Kane, M. H., Newton, C. J., Cornwell, M., Kothadia, R. B., Ma, W., Yoo, S., Mannan, R., Vats, P., Kumar-Sinha, C., Kawaler, E. A., Omelchenko, T., Colaprico, A., Geffen, Y., Maruvka, Y. E., da Veiga Leprevost, F., Wiznerowicz, M., Gumus, Z. H., Veluswamy, R. R., Hostetter, G., Heiman, D. I., Wyczalkowski, M. A., Hiltke, T., Mesri, M., Kinsinger, C. R., Boja, E. S., Omenn, G. S., Chinnaiyan, A. M., Rodriguez, H., Li, Q. K., Jewell, S. D., Thiagarajan, M., Getz, G., Zhang, B., Fenyo, D., Ruggles, K. V., Cieslik, M. P., Robles, A. I., Clauser, K. R., Govindan, R., Wang, P., Nesvizhskii, A. I., Ding, L., Mani, D. R., Carr, S. A., and Clinical Proteomic Tumor Analysis, C. (2020) Proteogenomic Characterization Reveals Therapeutic Vulnerabilities in Lung Adenocarcinoma. *Cell* **182**, 200-225 e235
75. Nguyen, B., Fong, C., Luthra, A., Smith, S. A., DiNatale, R. G., Nandakumar, S., Walch, H., Chatila, W. K., Madupuri, R., Kundra, R., Bielski, C. M., Mastrogiacomo, B., Donoghue, M. T. A., Boire, A., Chandarlapaty, S., Ganesh, K., Harding, J. J., Iacobuzio-Donahue, C. A., Razavi, P., Reznik, E., Rudin, C. M., Zamarin, D., Abida, W., Abou-Alfa, G. K., Aghajanian, C., Cercek, A., Chi, P., Feldman, D., Ho, A. L., Iyer, G., Janjigian, Y. Y., Morris, M., Motzer, R. J., O'Reilly, E. M., Postow, M. A., Raj, N. P., Riely, G. J., Robson, M. E., Rosenberg, J. E., Safonov, A., Shoushtari, A. N., Tap, W., Teo, M. Y., Varghese, A. M., Voss, M., Yaeger, R., Zauderer, M. G., Abu-Rustum, N., Garcia-Aguilar, J., Bochner, B., Hakimi, A., Jarnagin, W. R., Jones, D. R., Molena, D., Morris, L., Rios-Doria, E., Russo, P., Singer, S., Strong, V. E., Chakravarty, D., Ellenson, L. H., Gopalan, A., Reis-Filho, J. S., Weigelt, B., Ladanyi, M., Gonen, M., Shah, S. P., Massague, J., Gao, J., Zehir, A., Berger, M. F., Solit, D. B., Bakhoum, S. F., Sanchez-Vega, F., and Schultz, N. (2022) Genomic characterization of metastatic patterns from prospective clinical sequencing of 25,000 patients. *Cell* **185**, 563-575 e511

76. Van Allen, E. M., Miao, D., Schilling, B., Shukla, S. A., Blank, C., Zimmer, L., Sucker, A., Hillen, U., Foppen, M. H. G., Goldinger, S. M., Utikal, J., Hassel, J. C., Weide, B., Kaehler, K. C., Loquai, C., Mohr, P., Gutzmer, R., Dummer, R., Gabriel, S., Wu, C. J., Schadendorf, D., and Garraway, L. A. (2015) Genomic correlates of response to CTLA-4 blockade in metastatic melanoma. *Science* **350**, 207-211

77. Song, Y., Li, L., Ou, Y., Gao, Z., Li, E., Li, X., Zhang, W., Wang, J., Xu, L., Zhou, Y., Ma, X., Liu, L., Zhao, Z., Huang, X., Fan, J., Dong, L., Chen, G., Ma, L., Yang, J., Chen, L., He, M., Li, M., Zhuang, X., Huang, K., Qiu, K., Yin, G., Guo, G., Feng, Q., Chen, P., Wu, Z., Wu, J., Ma, L., Zhao, J., Luo, L., Fu, M., Xu, B., Chen, B., Li, Y., Tong, T., Wang, M., Liu, Z., Lin, D., Zhang, X., Yang, H., Wang, J., and Zhan, Q. (2014) Identification of genomic alterations in oesophageal squamous cell cancer. *Nature* **509**, 91-95

78. Bailey, P., Chang, D. K., Nones, K., Johns, A. L., Patch, A. M., Gingras, M. C., Miller, D. K., Christ, A. N., Bruxner, T. J., Quinn, M. C., Nourse, C., Murtaugh, L. C., Harliwong, I., Idrisoglu, S., Manning, S., Nourbakhsh, E., Wani, S., Fink, L., Holmes, O., Chin, V., Anderson, M. J., Kazakoff, S., Leonard, C., Newell, F., Waddell, N., Wood, S., Xu, Q., Wilson, P. J., Cloonan, N., Kassahn, K. S., Taylor, D., Quek, K., Robertson, A., Pantano, L., Mincarelli, L., Sanchez, L. N., Evers, L., Wu, J., Pinese, M., Cowley, M. J., Jones, M. D., Colvin, E. K., Nagrial, A. M., Humphrey, E. S., Chantrill, L. A., Mawson, A., Humphris, J., Chou, A., Pajic, M., Scarlett, C. J., Pinho, A. V., Giry-Laterriere, M., Rooman, I., Samra, J. S., Kench, J. G., Lovell, J. A., Merrett, N. D., Toon, C. W., Epari, K., Nguyen, N. Q., Barbour, A., Zeps, N., Moran-Jones, K., Jamieson, N. B., Graham, J. S., Duthie, F., Oien, K., Hair, J., Grutzmann, R., Maitra, A., Iacobuzio-Donahue, C. A., Wolfgang, C. L., Morgan, R. A., Lawlor, R. T., Corbo, V., Bassi, C., Rusev, B., Capelli, P., Salvia, R., Tortora, G., Mukhopadhyay, D., Petersen, G. M., Australian Pancreatic Cancer Genome, I., Munzy, D. M., Fisher, W. E., Karim, S. A., Eshleman, J. R., Hruban, R. H., Pilarsky, C., Morton, J. P., Sansom, O. J., Scarpa, A., Musgrove, E. A., Bailey, U. M., Hofmann, O., Sutherland, R. L., Wheeler, D. A., Gill, A. J., Gibbs, R. A., Pearson, J. V., Waddell, N., Biankin, A. V., and Grimmond, S. M. (2016) Genomic analyses identify molecular subtypes of pancreatic cancer. *Nature* **531**, 47-52

79. Zhang, J., Bajari, R., Andric, D., Gerthoffert, F., Lepsa, A., Nahal-Bose, H., Stein, L. D., and Ferretti, V. (2019) The International Cancer Genome Consortium Data Portal. *Nat Biotechnol* **37**, 367-369

80. Zhang, T., Joubert, P., Ansari-Pour, N., Zhao, W., Hoang, P. H., Lokanga, R., Moye, A. L., Rosenbaum, J., Gonzalez-Perez, A., Martinez-Jimenez, F., Castro, A., Muscarella, L. A., Hofman, P., Consonni, D., Pesatori, A. C., Kebede, M., Li, M., Gould Rothberg, B. E., Peneva, I., Schabath, M. B., Poeta, M. L., Costantini, M., Hirsch, D., Heselmeyer-Haddad, K., Hutchinson, A., Olanich, M., Lawrence, S. M., Lenz, P., Duggan, M., Bhawsar, P. M. S., Sang, J., Kim, J., Mendoza, L., Saini, N., Klimczak, L. J., Islam, S. M. A., Otlu, B., Khandekar, A., Cole, N., Stewart, D. R., Choi, J., Brown, K. M., Caporaso, N. E., Wilson, S. H., Pommier, Y., Lan, Q., Rothman, N., Almeida, J. S., Carter, H., Ried, T., Kim, C. F., Lopez-Bigas, N., Garcia-Closas, M., Shi, J., Bosse, Y., Zhu, B., Gordenin, D. A., Alexandrov, L. B., Chanock, S. J., Wedge, D. C., and Landi, M. T. (2021) Genomic and evolutionary classification of lung cancer in never smokers. *Nat Genet* **53**, 1348-1359

81. Pleasance, E., Titmuss, E., Williamson, L., Kwan, H., Culibrk, L., Zhao, E. Y., Dixon, K., Fan, K., Bowlby, R., Jones, M. R., Shen, Y., Grewal, J. K., Ashkani, J., Wee, K., Grisdale, C. J., Thibodeau, M. L., Bozoky, Z., Pearson, H., Majounie, E., Vira, T., Shenwai, R., Mungall, K. L., Chuah, E., Davies, A., Warren, M., Reisle, C., Bonakdar, M., Taylor, G. A., Csizmok, V., Chan, S. K., Zong, Z., Bilobram, S., Muhammadzadeh, A., D'Souza, D., Corbett, R. D., MacMillan, D., Carreira, M., Choo, C., Bleile, D., Sadeghi, S., Zhang, W., Wong, T., Cheng, D., Brown, S. D., Holt, R. A., Moore, R. A.,

Mungall, A. J., Zhao, Y., Nelson, J., Fok, A., Ma, Y., Lee, M. K. C., Lavoie, J. M., Mendis, S., Karasinska, J. M., Deol, B., Fisic, A., Schaeffer, D. F., Yip, S., Schrader, K., Regier, D. A., Weymann, D., Chia, S., Gelmon, K., Tinker, A., Sun, S., Lim, H., Renouf, D. J., Laskin, J., Jones, S. J. M., and Marra, M. A. (2020) Pan-cancer analysis of advanced patient tumors reveals interactions between therapy and genomic landscapes. *Nat Cancer* **1**, 452-468

82. Guo, Y. A., Chang, M. M., Huang, W., Ooi, W. F., Xing, M., Tan, P., and Skanderup, A. J. (2018) Mutation hotspots at CTCF binding sites coupled to chromosomal instability in gastrointestinal cancers. *Nat Commun* **9**, 1520
83. Wang, K., Yuen, S. T., Xu, J., Lee, S. P., Yan, H. H., Shi, S. T., Siu, H. C., Deng, S., Chu, K. M., Law, S., Chan, K. H., Chan, A. S., Tsui, W. Y., Ho, S. L., Chan, A. K., Man, J. L., Foglizzo, V., Ng, M. K., Chan, A. S., Ching, Y. P., Cheng, G. H., Xie, T., Fernandez, J., Li, V. S., Clevers, H., Rejto, P. A., Mao, M., and Leung, S. Y. (2014) Whole-genome sequencing and comprehensive molecular profiling identify new driver mutations in gastric cancer. *Nat Genet* **46**, 573-582
84. Dulak, A. M., Stojanov, P., Peng, S., Lawrence, M. S., Fox, C., Stewart, C., Bandla, S., Imamura, Y., Schumacher, S. E., Shefler, E., McKenna, A., Carter, S. L., Cibulskis, K., Sivachenko, A., Saksena, G., Voet, D., Ramos, A. H., Auclair, D., Thompson, K., Sougnez, C., Onofrio, R. C., Guiducci, C., Beroukhim, R., Zhou, Z., Lin, L., Lin, J., Reddy, R., Chang, A., Landrenau, R., Pennathur, A., Ogino, S., Luketich, J. D., Golub, T. R., Gabriel, S. B., Lander, E. S., Beer, D. G., Godfrey, T. E., Getz, G., and Bass, A. J. (2013) Exome and whole-genome sequencing of esophageal adenocarcinoma identifies recurrent driver events and mutational complexity. *Nat Genet* **45**, 478-486
85. Li, C., Sun, Y. D., Yu, G. Y., Cui, J. R., Lou, Z., Zhang, H., Huang, Y., Bai, C. G., Deng, L. L., Liu, P., Zheng, K., Wang, Y. H., Wang, Q. Q., Li, Q. R., Wu, Q. Q., Liu, Q., Shyr, Y., Li, Y. X., Chen, L. N., Wu, J. R., Zhang, W., and Zeng, R. (2020) Integrated Omics of Metastatic Colorectal Cancer. *Cancer Cell* **38**, 734-747 e739
86. Schulze, K., Imbeaud, S., Letouze, E., Alexandrov, L. B., Calderaro, J., Rebouissou, S., Couchy, G., Meiller, C., Shinde, J., Soysouvanh, F., Calatayud, A. L., Pinyol, R., Pelletier, L., Balabaud, C., Laurent, A., Blanc, J. F., Mazzaferro, V., Calvo, F., Villanueva, A., Nault, J. C., Bioulac-Sage, P., Stratton, M. R., Llovet, J. M., and Zucman-Rossi, J. (2015) Exome sequencing of hepatocellular carcinomas identifies new mutational signatures and potential therapeutic targets. *Nat Genet* **47**, 505-511
87. Rudin, C. M., Durinck, S., Stawiski, E. W., Poirier, J. T., Modrusan, Z., Shames, D. S., Bergbower, E. A., Guan, Y., Shin, J., Guillory, J., Rivers, C. S., Foo, C. K., Bhatt, D., Stinson, J., Gnad, F., Haverty, P. M., Gentleman, R., Chaudhuri, S., Janakiraman, V., Jaiswal, B. S., Parikh, C., Yuan, W., Zhang, Z., Koeppen, H., Wu, T. D., Stern, H. M., Yauch, R. L., Huffman, K. E., Paskulin, D. D., Illei, P. B., Varella-Garcia, M., Gazdar, A. F., de Sauvage, F. J., Bourgon, R., Minna, J. D., Brock, M. V., and Seshagiri, S. (2012) Comprehensive genomic analysis identifies SOX2 as a frequently amplified gene in small-cell lung cancer. *Nat Genet* **44**, 1111-1116
88. Um, S. W., Joung, J. G., Lee, H., Kim, H., Kim, K. T., Park, J., Hayes, D. N., and Park, W. Y. (2016) Molecular Evolution Patterns in Metastatic Lymph Nodes Reflect the Differential Treatment Response of Advanced Primary Lung Cancer. *Cancer Res* **76**, 6568-6576
89. Kan, Z., Ding, Y., Kim, J., Jung, H. H., Chung, W., Lal, S., Cho, S., Fernandez-Banet, J., Lee, S. K., Kim, S. W., Lee, J. E., Choi, Y. L., Deng, S., Kim, J. Y., Ahn, J. S., Sha, Y., Mu, X. J., Nam, J. Y., Im, Y. H., Lee, S., Park, W. Y., Nam, S. J., and Park, Y. H. (2018) Multi-omics profiling of younger Asian breast cancers reveals distinctive molecular signatures. *Nat Commun* **9**, 1725

90. Chen, J., Yang, H., Teo, A. S. M., Amer, L. B., Sherbaf, F. G., Tan, C. Q., Alvarez, J. J. S., Lu, B., Lim, J. Q., Takano, A., Nahar, R., Lee, Y. Y., Phua, C. Z. J., Chua, K. P., Suteja, L., Chen, P. J., Chang, M. M., Koh, T. P. T., Ong, B. H., Anantham, D., Hsu, A. A. L., Gogna, A., Too, C. W., Aung, Z. W., Lee, Y. F., Wang, L., Lim, T. K. H., Wilm, A., Choi, P. S., Ng, P. Y., Toh, C. K., Lim, W. T., Ma, S., Lim, B., Liu, J., Tam, W. L., Skanderup, A. J., Yeong, J. P. S., Tan, E. H., Creasy, C. L., Tan, D. S. W., Hillmer, A. M., and Zhai, W. (2020) Genomic landscape of lung adenocarcinoma in East Asians. *Nat Genet* **52**, 177-186

91. Pickering, C. R., Zhou, J. H., Lee, J. J., Drummond, J. A., Peng, S. A., Saade, R. E., Tsai, K. Y., Curry, J. L., Tetzlaff, M. T., Lai, S. Y., Yu, J., Muzny, D. M., Doddapaneni, H., Shinbrot, E., Covington, K. R., Zhang, J., Seth, S., Caulin, C., Clayman, G. L., El-Naggar, A. K., Gibbs, R. A., Weber, R. S., Myers, J. N., Wheeler, D. A., and Frederick, M. J. (2014) Mutational landscape of aggressive cutaneous squamous cell carcinoma. *Clin Cancer Res* **20**, 6582-6592

92. Durinck, S., Stawiski, E. W., Pavia-Jimenez, A., Modrusan, Z., Kapur, P., Jaiswal, B. S., Zhang, N., Toffessi-Tcheuyap, V., Nguyen, T. T., Pahuja, K. B., Chen, Y. J., Saleem, S., Chaudhuri, S., Heldens, S., Jackson, M., Pena-Llopis, S., Guillory, J., Toy, K., Ha, C., Harris, C. J., Holloman, E., Hill, H. M., Stinson, J., Rivers, C. S., Janakiraman, V., Wang, W., Kinch, L. N., Grishin, N. V., Haverty, P. M., Chow, B., Gehring, J. S., Reeder, J., Pau, G., Wu, T. D., Margulis, V., Lotan, Y., Sagalowsky, A., Pedrosa, I., de Sauvage, F. J., Brugarolas, J., and Seshagiri, S. (2015) Spectrum of diverse genomic alterations define non-clear cell renal carcinoma subtypes. *Nat Genet* **47**, 13-21

93. Stransky, N., Egloff, A. M., Tward, A. D., Kostic, A. D., Cibulskis, K., Sivachenko, A., Kryukov, G. V., Lawrence, M. S., Sougnez, C., McKenna, A., Shefler, E., Ramos, A. H., Stojanov, P., Carter, S. L., Voet, D., Cortes, M. L., Auclair, D., Berger, M. F., Saksena, G., Guiducci, C., Onofrio, R. C., Parkin, M., Romkes, M., Weissfeld, J. L., Seethala, R. R., Wang, L., Rangel-Escareno, C., Fernandez-Lopez, J. C., Hidalgo-Miranda, A., Melendez-Zajgla, J., Winckler, W., Ardlie, K., Gabriel, S. B., Meyerson, M., Lander, E. S., Getz, G., Golub, T. R., Garraway, L. A., and Grandis, J. R. (2011) The mutational landscape of head and neck squamous cell carcinoma. *Science* **333**, 1157-1160

94. Rokita, J. L., Rathi, K. S., Cardenas, M. F., Upton, K. A., Jayaseelan, J., Cross, K. L., Pfeil, J., Egolf, L. E., Way, G. P., Farrel, A., Kendsersky, N. M., Patel, K., Gaonkar, K. S., Modi, A., Berko, E. R., Lopez, G., Vaksman, Z., Mayoh, C., Nance, J., McCoy, K., Haber, M., Evans, K., McCalmont, H., Bendak, K., Bohm, J. W., Marshall, G. M., Tyrrell, V., Kalletla, K., Braun, F. K., Qi, L., Du, Y., Zhang, H., Lindsay, H. B., Zhao, S., Shu, J., Baxter, P., Morton, C., Kurmashev, D., Zheng, S., Chen, Y., Bowen, J., Bryan, A. C., Leraas, K. M., Coppens, S. E., Doddapaneni, H., Momin, Z., Zhang, W., Sacks, G. I., Hart, L. S., Krytska, K., Mosse, Y. P., Gatto, G. J., Sanchez, Y., Greene, C. S., Diskin, S. J., Vaske, O. M., Haussler, D., Gastier-Foster, J. M., Kolb, E. A., Gorlick, R., Li, X. N., Reynolds, C. P., Kurmasheva, R. T., Houghton, P. J., Smith, M. A., Lock, R. B., Raman, P., Wheeler, D. A., and Maris, J. M. (2019) Genomic Profiling of Childhood Tumor Patient-Derived Xenograft Models to Enable Rational Clinical Trial Design. *Cell Rep* **29**, 1675-1689 e1679

95. Hyman, D. M., Piha-Paul, S. A., Won, H., Rodon, J., Saura, C., Shapiro, G. I., Juric, D., Quinn, D. I., Moreno, V., Doger, B., Mayer, I. A., Boni, V., Calvo, E., Loi, S., Lockhart, A. C., Erinjeri, J. P., Scaltriti, M., Ulaner, G. A., Patel, J., Tang, J., Beer, H., Selcuklu, S. D., Hanrahan, A. J., Bouvier, N., Melcer, M., Murali, R., Schram, A. M., Smyth, L. M., Jhaveri, K., Li, B. T., Drilon, A., Harding, J. J., Iyer, G., Taylor, B. S., Berger, M. F., Cutler, R. E., Jr., Xu, F., Butturini, A., Eli, L. D., Mann, G., Farrell, C., Lalani, A. S., Bryce, R. P., Arteaga, C. L., Meric-Bernstam, F., Baselga, J., and Solit, D. B. (2018)

HER kinase inhibition in patients with HER2- and HER3-mutant cancers. *Nature* **554**, 189-194
